# Mutational Profiling of SARS-CoV-2 PLpro in human cells reveals requirements for function, structure, and drug escape

**DOI:** 10.1101/2024.03.11.584210

**Authors:** Xinyu Wu, Margareta Go, Julie V. Nguyen, Nathan W. Kuchel, Bernadine G.C. Lu, Kym N. Lowes, Dale J. Calleja, Jeffrey P. Mitchell, Guillaume Lessene, David Komander, Matthew E. Call, Melissa J. Call

## Abstract

SARS-CoV-2, the causative agent of COVID-19, is responsible for the recent global pandemic and remains a major source of mortality. Papain-like protease (PLpro) is a target for SARS-CoV-2 inhibitor development, as it is not only essential for viral replication through cleavage of the viral poly-proteins pp1a and pp1ab, but also has de-ubiquitylation and de-ISGylation activities, which can affect innate immune responses. To understand the features of PLpro that dictate activity and anticipate how emerging PLpro variants will affect function, we employed Deep Mutational Scanning to evaluate the mutational effects on enzymatic activity and protein stability in mammalian cells. We confirm features of the active site and identify all mutations in neighboring residues that support or ablate activity. We characterize residues responsible for substrate binding and demonstrate that although the blocking loop is remarkably tolerant to nearly all mutations, its flexibility is important for enzymatic function. We additionally find a connected network of mutations affecting function but not structure that extends far from the active site. Using our DMS libraries we were able to identify drug-escape variants to a common PLpro inhibitor scaffold and predict that plasticity in both the S4 pocket and blocking loop sequence should be considered during the drug design process.

## Introduction

COVID-19, caused by SARS-CoV-2 has swiftly become one of the leading causes of death globally and, despite robust population-level immunity, remains so four years after SARS-CoV-2 emerged in the human population. SARS-CoV-2 can be treated by inhibition of crucial viral proteases. SARS-CoV-2, like SARS-CoV and other coronaviruses, relies on the Main protease (Mpro) and Papain-Like protease (PLpro) to cleave virally encoded poly-protein chains that contain non-structural proteins required for viral replication. Development of Pfizer’s SARS-CoV-2 Mpro inhibitor, Nirmatrelvir (Owen et al. 2021) relied on lead compounds to a highly homologous coronavirus, SARS-CoV, which emerged in 2004. Further development was abandoned when SARS-CoV virus was eliminated in the human population but resumed in 2020 during the COVID-19 pandemic. Nirmatrelvir demonstrates that protease inhibition is effective in SARS-CoV-2 treatment, but viral escape from existing single-agent therapeutics remains a concern.

Based on recent findings and knowledge of SARS-CoV PLpro function, PLpro is an active enzymatic domain of non-structural protein 3 (Nsp3) (Fig. **1a**). Like Mpro, PLpro is an attractive target for inhibitor development because of its proteolytic role in releasing Nsp1, Nsp2, Nsp3, and in conjunction with Mpro, Nsp4 from the viral poly-proteins pp1a and pp1ab (Harcourt et al. 2004). Additionally, PLpro efficiently removes the post-translational modifications of ubiquitin and interferon-stimulated gene 15 (ISG15) from viral and host proteins (Barretto et al. 2005; Lindner et al. 2005; Lindner et al. 2007; Klemm et al. 2020), a mechanism that likely protects viral proteins (Durfee et al. 2010) and hampers immune signaling pathways (Perng and Lenschow 2018; Gold et al. 2022; Skaug and Chen 2010; Shin et al. 2020; Zhang et al. 2021). Proteolytic cleavage, de-ubiquitination and de-ISGylation activity relies on the motif common among all substrates, Leu_P4_X_P3_Gly_P2_Gly_P1_↓ where X is commonly Lys or Arg and the arrow represents the site of cleavage (Rut et al. 2020).

**Fig. 1.**
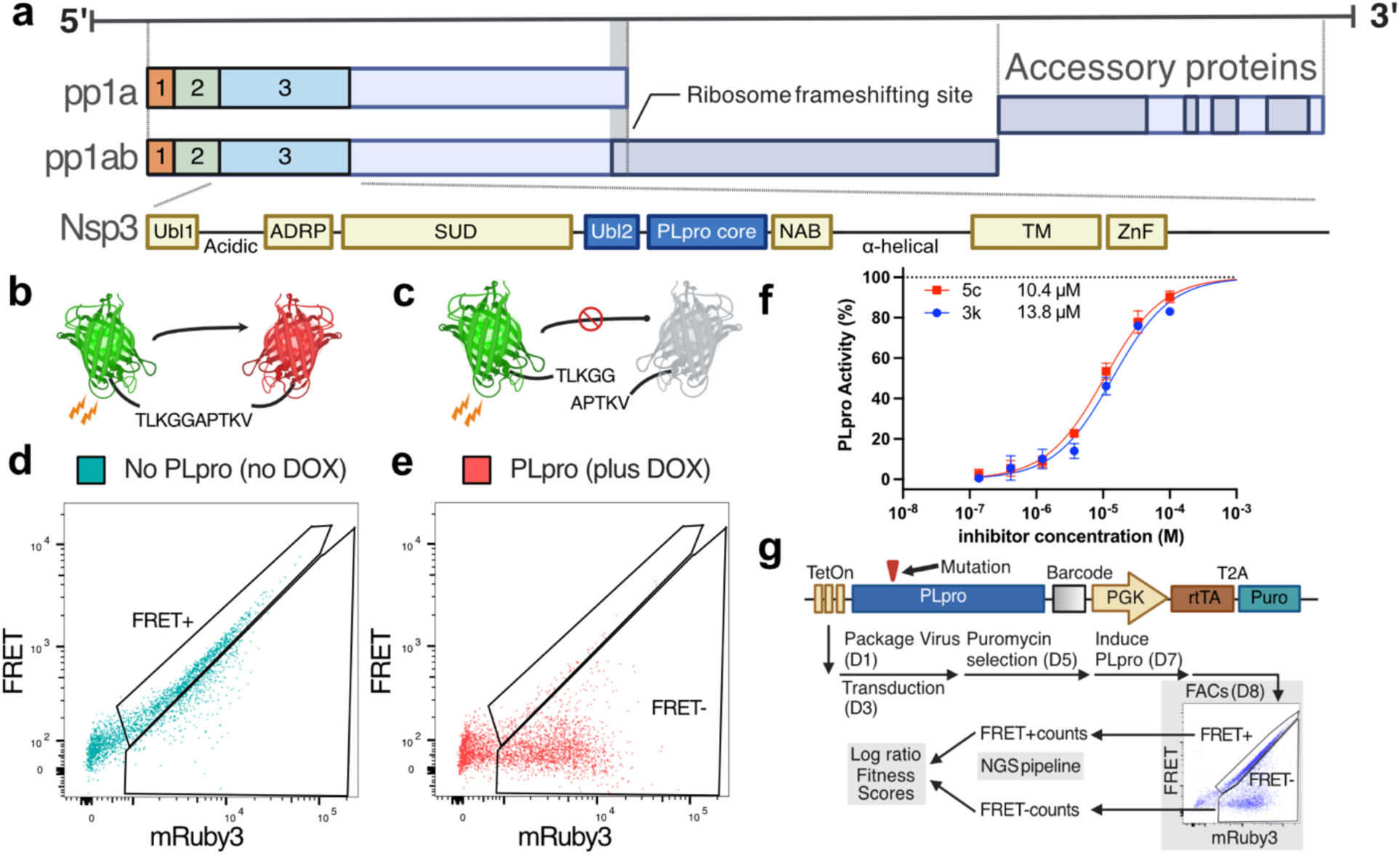
SARS-CoV-2 genome and cellular assays for PLpro Activity. **a)** SARS-CoV-2 genome indicating position of PLpro. Numbered domains 1, 2 and 3 refer to the first three non-structural proteins that are released by PLpro. PLpro is a domain of Non-Structural Protein 3 (Nsp3). The other domains of Nsp3 are ubiquitin-like domain 1 (Ubl1), ADP-ribose phosphatase (ADRP), SARS-unique domain (SUD), ubiquitin-like domain 2 (Ubl2), PLpro core, nucleic acid binding domain (NAB), transmembrane domain (TM) and zinc finger motif (ZnF) (Armstrong et al. 2021) **b)** Illustration of the FRET biosensor. mClover3 (in green) is covalently linked to mRuby3 (in red) with the linker sequence TLKGGAPTKV. Excitation of mClover3 transfers energy to mRuby3. **c)** Cleavage of the linker sequence by PLpro releases mRuby3 from mClover3 donor fluorophore, reducing FRET. **d)** Flow cytometry plot of cells containing the FRET biosensor prior to induction of PLpro expression with dox. The x-axis shows emission from the 610/20 filter after excitation with the 561 nm laser (mRuby3), the y-axis shows emission from the 610/20 filter after excitation with the 488 nm laser (FRET). **e)** The same cells in (d) 24 hours after dox addition to induce PLpro expression, showing cleavage of the FRET biosensor. **f)** Dose response of 3k and 5c in the cellular assay described in d) & e). Data were normalized so the top was 100 and bottom was 0 after fitting the 5c dose response curve that had a Hill slope of 1. EC50 values for this representative experiment were calculated as: 13.8 µM (3k); 10.4 µM (5c), n = 3 and error bars are mean ± SD. Note: the 5c EC50 over 8 biological replicates is 15.5 ± 1.3 SEM (*Extended data Fig. 5c*) **g)** Schematic of the Activity DMS workflow. The linear vector map shows the PLpro expression construct with TetOn promoter element driving PLpro expression. A 16bp barcode is installed 3’ of the PLpro coding sequence. This is followed by a constitutive PGK promoter that drives rtTA and Puromycin resistance. The workflow describes the PLpro Activity DMS assay, starting from viral packaging, reporter cell transduction with the library, dox induction, selection by FACS and data processing. Panels a), b) and c) were created with BioRender.

The PLpro core has a domain architecture that resembles human de-ubiquitinases from the USP family and comprises “thumb”, “palm” and “fingers” domains. The active site residues are found on a flat surface, split among the thumb (Cys111) and palm (His272 and Asp286) domains, while the fingers domain contains a zinc binding site coordinated by Cys189, Cys192, Cys224, and Cys226 that is required for overall structural integrity (Barretto et al. 2005). Because the P1 and P2 positions of the cleavage motif are glycines, entry to the active site is narrow and the active site itself is shallow, making it spatially confined. Most well-characterized inhibitors of SARS-CoV-2 PLpro instead target the S4 pocket that accommodates the P4 Leu of the recognition sequence motif (Rut et al. 2020) and engage a flexible β-hairpin, termed the blocking loop, that flips up to cover the pocket (Calleja, Lessene, and Komander 2022; Ton et al. 2022; Lv et al. 2021).

Known inhibitors are focused on a few primary scaffolds exemplified by GRL0617 (Ratia et al. 2008b), and the 5c/3k series (Báez-Santos, Barraza, et al. 2014), which were both developed from hits in high-throughput screens against SARS-CoV PLpro. GRL0617 and 5c/3k share a naphthyl ring that binds the S4 pocket, with polar contacts further stabilizing binding through Asp164 and a substituted phenyl ring that induces a conformation change in Leu162 allowing the compounds to be accommodated (Calleja, Lessene, and Komander 2022). Recently reported compounds XR8-89 and Jun9-84-3 build on the GRL0617 scaffold. XR8-89 by replacing the naphthalene with a substituted phenylthiophene, and introducing the azetidine group to bind with Glu167(Shen et al. 2021) and Jun9-84-3 by replacing the phenyl ring with an indole ring, increasing potency (Ma et al. 2021; Xia et al. 2021). Modifications have also been performed on 5c and 3k scaffolds to optimize absorption, distribution, metabolism and excretion (ADME) properties (Calleja et al. 2022).

In this study, we establish an assay to measure PLpro activity in mammalian cells and construct a Deep Mutational Scanning (DMS) (Fowler and Fields 2014) library to assess the impact of every possible single-site substitution on proteolytic activity. We further examined the mutational effects on protein stability by linking a fluorescent reporter to PLpro to measure the abundance of each variant in cells, allowing variants that impact PLpro activity, but not its folding, to be elucidated. We also used our library to predict variants of PLpro that escape the PLpro inhibitor scaffold exemplified by small molecule compounds 3k and 5c (Báez-Santos, Barraza, et al. 2014; Klemm et al. 2020; Calleja et al. 2022). Top-ranked escape variants were validated individually in cellular assays and recombinant enzymatic assays. In combination, these data elucidate all regions of PLpro that are required for activity and indicate a role for PLpro flexibility in proteolytic cleavage and drug escape.

## Results

### Development of DMS-suitable cell-based assay to measure PLpro proteolytic activity

Deep mutational scanning examines the functional effects of *all* single-site substitutions in a region of interest of a protein. We chose to examine 315 residues of Nsp3 spanning residues 746 to 1060, which encompasses the PLpro core and the ubiquitin-like domain N-terminal to PLpro. This region can be expressed in isolation, remaining enzymatically active and in the remainder of the paper, we use PLpro numbering (1 to 315). Within our construct there are, including wildtype, 6301 theoretical protein variants, numbers which make individual assessment of each variant a formidable task. Instead, we used DMS to assess the function of each variant in a pooled experiment with next-generation sequencing used to count variants in selected populations. From these counts, ratiometic scores are calculated to elucidate the enrichment of each variant in the pool during selection. This requires careful assay design where genetic material encoding the variant and the variant protein itself must co-segregate in any selection step. One way to accomplish this is to measure activity in a cell, where the cell membrane encompasses both the protein variant and the genetic material encoding it and to sort cells with functional variants from those that are impaired.

We elected to develop an assay for PLpro activity in a human cell line to closely mimic the native environment during viral replication. To measure proteolytic activity in cells, we converted an existing luciferase-based reporter (Zhang et al. 2017) into a FRET-based biosensor compatible with fluorescence activated cell sorting (FACS). Our FRET-based biosensor is composed of N-terminal mClover3 donor and C-terminal mRuby3 acceptor fluorophores separated by a linker containing the Nsp2/3 PLpro cleavage motif T_P5_L_P4_K_P3_G_P2_G_P1_↓A_P-1_P_P-2_T_P-3_K_P-4_V_P-5_ (Fig. **1b**). We introduced this biosensor into 293T cells via retroviral transduction to create a stable PLpro reporter cell line. A FRET signal between mClover3 and mRuby3 was readily observable with excitation at 488 nm and emission at 610 nm (Extended data Fig. 1a).

To validate the biosensor’s ability to report on PLpro activity, we installed the PLpro coding sequence into a lentiviral Tet-On expression vector. Upon introduction of this construct into our 293T biosensor cell line, FRET emission was reduced in a doxycycline (dox) dependent manner, indicating biosensor cleavage (Fig. **1b-e**). Inhibition of PLpro protected the biosensor from cleavage: dose-response curves with two PLpro inhibitors 3k and 5c (Báez-Santos, Barraza, et al. 2014) reported EC50s of 13.8 µM and 10.4 µM (Fig. **1f**). We found this approach also worked to create cellular assays for PLpro of other coronaviruses. We designed biosensors that were able to report activities of SARS-CoV, and MERS PLpro, by adjusting the flexible linkers (Extended data Fig. 1b). Together, these data indicate that our SARS-CoV-2 PLpro biosensor is stable in cells until PLpro expression is induced and that inhibition of PLpro can be faithfully measured, providing a cellular assay that is suitable for DMS.

### Construction and validation of a DMS library

Several methods are reported to generate DMS libraries in high-throughput manner, including nicking mutagenesis (Wrenbeck et al. 2016; Steiner, Baumer, and Whitehead 2020), PCR-based site-directed mutagenesis (Shenoy and Visweswariah 2003; Trenker et al. 2021), codon-mutagenesis (Bloom 2014; Starr et al. 2020) and Mutagenesis by Integrated TilEs (MITE) (Melnikov et al. 2014). For residues close to the active site (encompassing residues 62, 69-70, 73-74, 77, 93, 104, 106-119, 151-174, 206-212, 243-253, 260-276, 285-286, 296-304), we ordered dsDNA with gateway adapters from Twist Bioscience and directly cloned these fragments into our lentiviral vector based on FU-tetO-Gateway (Addis et al. 2011) that had been modified to include a puromycin resistance cassette. To create the remainder of the library, we elected to adapt the MITE method (Melnikov et al. 2014) and directly cloned single-strand oligos in tiled cohorts to cover the entire length of PLpro. We additionally included a 16 bp barcode 3’ PLpro coding region to simplify the sequencing strategy (Extended data Fig. 2). Once mixed, the full library was characterised by PacBio long-read sequencing to map barcodes to variants. We found 6263 of the 6300 protein variants were present in the library representing 99.4% coverage and each variant was replicated with an average of 13 barcodes, except wildtype which aligned to 6657 barcodes.

### DMS pinpoints structurally and functionally conserved residues of PLpro

We transduced our DMS library into 20 million FRET-biosensor-containing 239T reporter cells at a multiplicity of infection of ∼0.2 to ensure that cells that received PLpro only had one copy. Lentiviral virions contain two RNA strands, so to prevent barcode swapping and double transduction of two different variants (Hill et al. 2018) we mixed our library with an unrelated lentiviral transfer vector containing BFP at a ratio of 1:2 before we made virus. Non-transduced cells were subsequently removed by puromycin treatment, leaving an estimated 2 million unique transductants in each replicate and providing approximately 10-fold representation of the library before selection.

To perform the screen based on biosensor cleavage, we induced PLpro with dox for 24 hours prior to sorting cells into FRET positive (inactive PLpro) and negative (active PLpro) gates (Fig. **1g**). mRNA was prepared from cells in each population, reverse transcribed and barcodes sequenced by Illumina sequencing. Ratiometric scores for each variant were calculated with DimSum (Faure et al. 2020) to determine which variants were enriched in the negative FRET gate that contained active PLpro variants.

The data quality was first checked by comparing the distribution of positive and negative controls, which were synonymous variants that encode wildtype protein but have variant DNA codons, and nonsense variants with premature stop codons, respectively (Fig. **2a**). We saw good separation of scores, indicating that selection was successful.

**Fig. 2.**
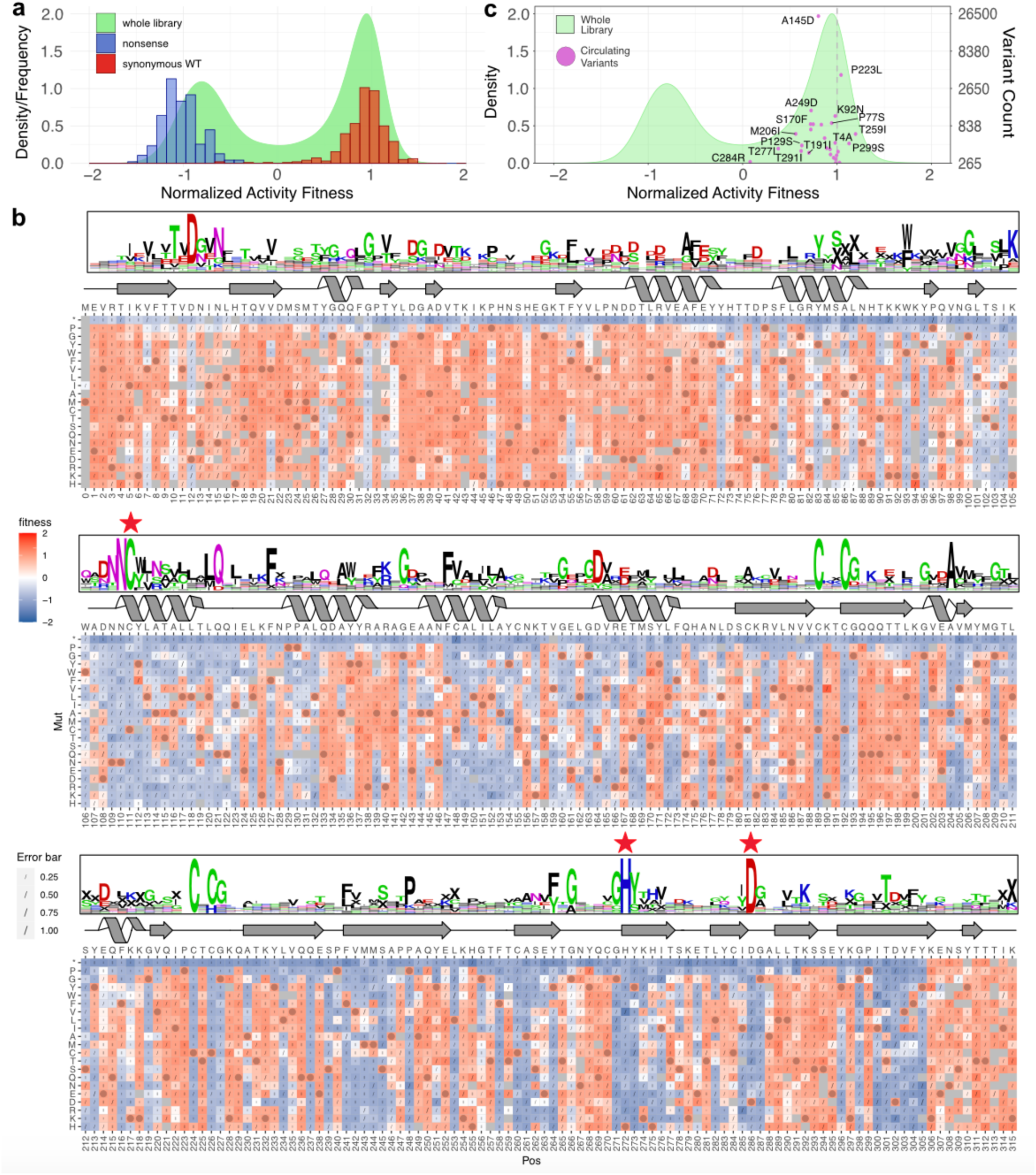
Sequence-Function map of PLpro activity and observed circulating variants. **a)** The distribution of normalized fitness scores for the whole library (density; green), overlaid with the frequency of scores from synonymous wildtype variants (red; set at 1) and nonsense variants at positions 1-305 (blue; set at −1). **b)** The normalized sequence-function map of activity data from 7 independent transductions. The bottom x-axis indicates the position; the top x-axis indicates WT sequence; and the y-axis indicates the mutation. Fitness is shown in a two-color gradient with red indicating active variants, blue inactive variants, and grey missing variants. Wildtype variants are marked with a circle. Errors (sigma) from DimSum are indicated with a slash, the size of which scales with the extent of the error. Secondary structure is depicted along the top of the map and based on (PDB: 7CJM) (Fu et al. 2021). Arrows denote β-sheet; helices denote α-helix and lines denote random coil. Three red stars highlight the active triad: C111, H272, D286. On top of the secondary structure diagram, a sequence alignment of 22 PLpro sequences from different coronavirus species (Supplementary Table 3) is shown in weblog format. Positively charged residues are shown in blue; negatively charged residues shown in red; polar residues are shown in green; and others in black. “X” was used to fill in gaps in the alignment. The bigger the letter is, the more conserved the residue. **c)** The density plot of the whole library in (a) was overlaid with the score (x-axis) and variant counts (right y-axis, in log scale) of circulating variants (magenta) identified from COVID-3D (Portelli et al. 2020). Some commonly observed variants are labeled; refer to Supplementary Table 4 for a full list of variants and scores used in this study. The least active variant C284R is also labeled. Interestingly, the canonical sequence of SARS-CoV PLpro also has an Arg at this position.

We normalized the dataset to place the mean of synonymous wildtype on 1 and nonsense variants on −1, filtered the data to remove variants with low counts and high errors (Extended data Fig. 3) and calculated a sequence-function heatmap to view the data in its entirety (Fig. **2b**). As expected, no mutations were permitted at the PLpro active site (Cys111, His272, Asp286). Additionally, the four cysteines (Cys189, Cys192, Cys224, Cys226) that tetrahedrally coordinate the zinc in the fingers domain (Barretto et al. 2005) were absolutely required except for Cys224, where another zinc coordinating residue (histidine) was tolerated. This may introduce a non-classical C2-CH zinc finger (Cassandri et al. 2017) and thereby maintain structural integrity. All nonsense mutations were inactive except for those that came after Tyr305, which is the end of a major β-sheet (Fig. **2b**) in the thumb domain, and residues following Tyr305 may not be required for PLpro structural integrity and hence functionality. Viral genome sequencing over the course of the pandemic has identified naturally occurring PLpro variants (Portelli et al. 2020). These variants all had scores indicating some level of activity, with all but a handful clustering close to wildtype (Fig. **2c**). Those with lower activity scores and frequency may be sequencing errors or may occur in combination with other mutations that restore activity.

To check that our DMS activity scores aligned well with conservation among the PLpro of different coronavirus species, we aligned multiple PLpro sequences from SARS-CoV-2, MERS and other coronaviruses (Chaudhuri et al. 2011; see supplementary materials) and calculated a WebLogo representation. Although sequence conservation is relatively low between species, some residues are highly conserved. In our DMS dataset these same residues are, almost without exception, intolerant of mutation (Fig. **2b**).

As variants with folding defects will have reduced activity, we expect that residues on the surface of PLpro will be more tolerant of mutation than those within the core of the protein. To assess this, we averaged the activity scores of all variants for each position and plotted the resulting score on a ribbon diagram of the PLpro structure, where red indicates tolerant and blue intolerant to mutation (Fig. **3a**). In addition to the active and zinc-coordination sites (Fig. **3a**) we saw that residues within the papain-like fold were largely intolerant of mutation (Fig. **3b**). In comparison, the N-terminal ubiquitin-like domain is largely tolerant. This agrees with findings from SARS-CoV PLpro showing that the ubiquitin-like domain can be truncated with minimal impact on activity (Báez-Santos, Mielech, et al. 2014).

**Fig. 3.**
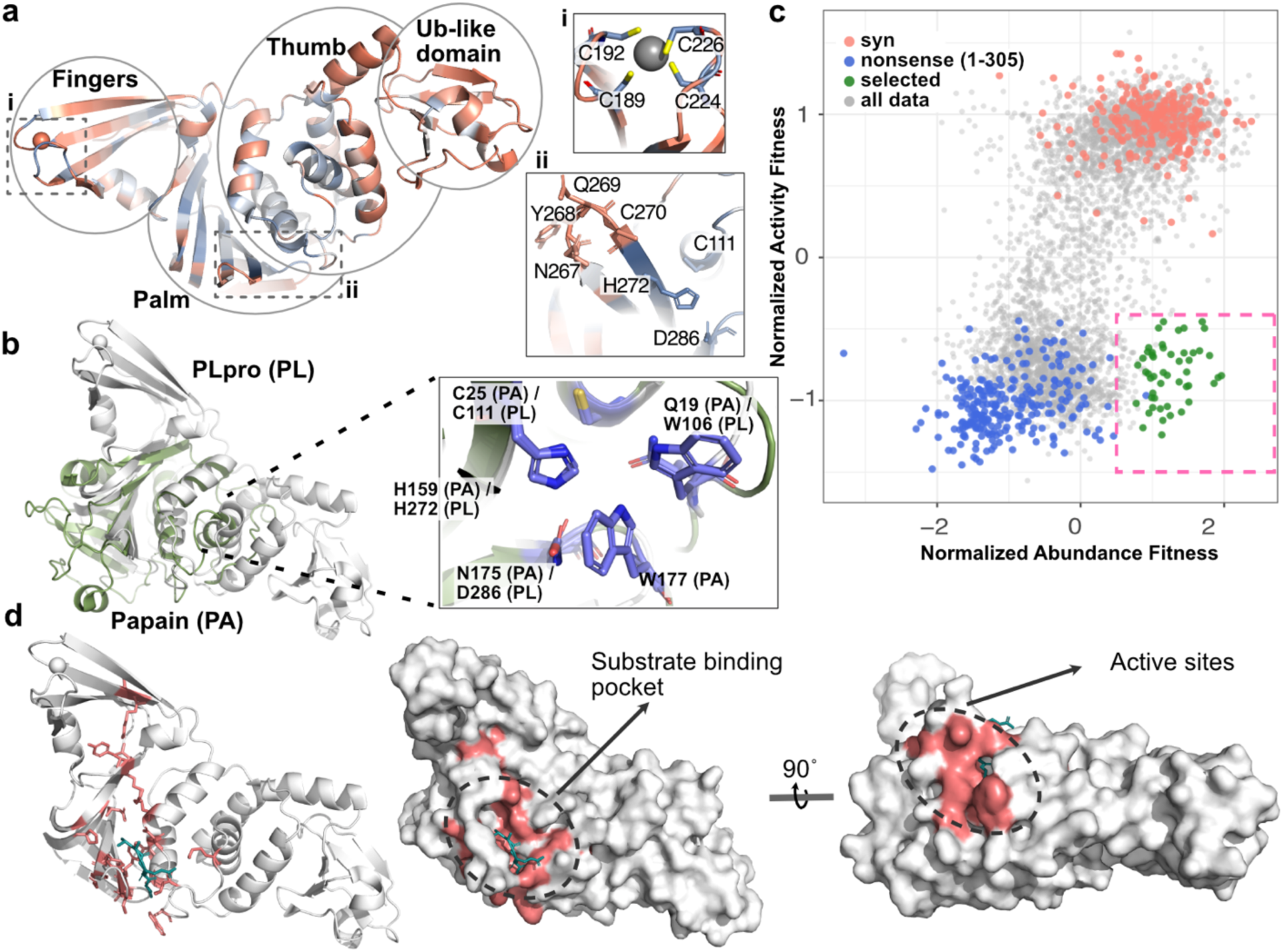
Structural analysis of activity fitness scores before and after filtering for abundance. **a)** The PLpro structure (PDB:6XAA) (Klemm et al. 2020) is represented in ribbon diagram with the fingers, palm, thumb and ubiquitin like domains labeled. Two additional regions containing the zinc-coordination site (**i**) and, active site and blocking loop (**ii**) are boxed and expanded to the right. The ribbon and highlighted areas in box i) and ii) are colored according to the average fitness score among all variants at each position. Red positions are tolerant to mutation and blue positions are intolerant. In box (i), C189, C192, C224, C226 are shown in stick and the zinc atom is shown as sphere. In box ii), residues on blocking loop and active triad (C111, H272, D286) are shown in stick. **b)** The papain active site (PDB: 9PAP) (Kamphuis et al. 1984) was aligned on the active site of PLpro (PDB: 6XAA, shown in white) and the regions of papain that showed good structural correlation are shown in green and account for ∼25% of the full-length papain protein. The overlay of the active site is shown on the right. Key PLpro (PL) and Papain (PA) residues are shown in stick and labeled accordingly. **c)** A scatterplot of Normalized Activity Fitness Scores against Normalized Abundance Fitness Scores (from two independent transductions, see *Extended data Fig. 4 for Abundance Sequence-Function map*.) identifies variants with normal abundance, but impaired function. Variants from the whole library are colored in grey, nonsense variants (positions 1-305) are colored in blue, and synonymous wildtype variants are colored in red. Variants with small enough errors to confidently place them in the box at the lower right of the scatter plot were selected as abundant but impaired and are colored in green. **d)** *left)* The position of abundant but impaired variants are indicated on the PLpro structure in stick and magenta. The last 4 residues of Ubiquitin are shown in green to highlight the substrate binding region. Comparison with b) above shows almost all abundant but impaired variants have mutations that fall within the area structurally conserved with Papain. *middle)* Surface representation of (d-left). *right)* PLpro structure in (d-middle) rotated 90° around *x* to show the active site.

### Identification of mutations that affect PLpro function but not abundance

The DMS dataset measuring PLpro activity has revealed all single-site substitutions that either maintain or impact PLpro activity. Impaired variants may carry a mutation that impacts abundance, most likely via folding or stability defects, or function, giving mechanistic insight. To identify abundant but functionally impaired variants, we reformatted the library to install a C-terminal mClover3 fusion (Extended data Fig. 4a), allowing PLpro abundance to be monitored by flow cytometry. Similar methods have been used to measure protein abundance in other systems (Matreyek et al. 2018; Thorn 2017; Csibra and Stan 2022). We expressed this library in parental 293T cells and sorted cells from high and low mClover3 gates, recovered mRNA to count variants, and calculated ratiometric scores to determine the abundance of each variant.

In reformatting our library, we lost some redundancy, but after filtering the data to remove variants with high errors, we were still able to calculate confident abundance scores for 4888 variants (∼80% of all potential 6301 variants) and saw segregation among synonymous and nonsense variants (Extended data Fig. 4b-e). When activity scores were plotted against abundance scores, we saw good correlation among most data points. We gated inactive variants with WT-like abundance and defined these variants, encompassing 22 residue positions, as structurally intact but functionally impaired (Fig. **3c**; Table 1).

**Table 1.** Classification of Functionally Important Residues. The classifications are performed based on Extended data Fig. 17. In the table, inactive variants are in bold; partially active variants are in italics; active variants are neither bolded nor in italics. For variants missing data, activity information is provided with the same font treatment, except for variants missing both activity and abundance scores, which are marked with an asterisk.

|  | Abundant |  | Partially Abundant |  | Low Abundance |  |  |  |
| --- | --- | --- | --- | --- | --- | --- | --- | --- |
| Position | Inactive | Active | Active | Partially active | Inactive | Active | Missing Data | Classification |
| W106 | **C,E,G,**<br>**H,K,L,**<br>**M,V** |  |  | *A,F,I,N,*<br>*S* |  |  | **D,T,R**<br>**Q\*,Y\*,**<br>**P\*** | Oxyanion hole (Báez-Santos, St John, and Mesecar 2015); similar position to Gln19 in papain (Ménard et al. 1991) |
| C111 | **E,G,H,**<br>**M,Q,S,**<br>**T** |  |  |  | **D,I,L,N,**<br>**R,V,W,**<br>**Y** |  | **K,F,A**<br>**P\*** | Active site |
| T115 | **E,Q** | A,C,V |  | *D,G* | **F,H,I,L,**<br>**M,P,W,**<br>**Y** |  | **N**<br>**K,R**<br>**S\*** | Second shell |
| L118 | **E** |  |  | *C,M,V* | **A,F,I,K,**<br>**N,Q,R,**<br>**S,T,W,**<br>**Y** |  | **P,G,H,**<br>**D** | Second shell similar position to Val32 in papain - Interdomain flexibility (Choudhury et al. 2010) |
| K157 | **Y** |  |  | *H,Q,R* | **A,C,D,**<br>**E,F,G,**<br>**N,S,T,V** |  | **W,L,P,I**<br>**M\*** | Unknown, may stabilise L162 |
| L162 | **D,I** | C,G,K,<br>M,R,T,<br>Y |  | *E,H,Q,*<br>*S,W* | **N** |  | **V,A,F**<br>**P\*** | Substrate binding (Ratia et al. 2006) |
| D164 | **K,T,V,Y** | C,E |  | *G,N* | **F,H,W** |  | **S**<br>**R,L,I,P**<br>**Q\*,M\*,**<br>**A\*** | Substrate binding (Fu et al. 2021; Perlinska et al. 2022) |
| R166 | **G,S,V,**<br>**W,Y** |  |  | *A,K* | **D,E,I,L,**<br>**M,N,P,**<br>**Q,T** |  | **C,H**<br>**F\*** | Lines the S4 pocket. Unknown, may stabilise D164 (Ma et al. 2021) |
| R183 | **F,I,L,V,**<br>**W** | G,S |  | *Q* | **E,H,K,**<br>**P,T,Y** |  | **A**<br>**M,C,N,**<br>**D** | Unknown |
| V184 | **H,W** | I,K,T |  | *C,L,M,*<br>*Q,R,S* | **A,D,F,**<br>**G,N,P,**<br>**Y** |  | **E\*** | Unknown |
| Y207 | **P** | A,G,M,<br>S,V,W |  | *D,I,L,N,*<br>*Q,T* | **R** |  | **E,C,H**<br>**K**<br>**F\*** | Unknown |
| V242 | **D,E** | I,L |  | *C,M* | **A,F,G,**<br>**H,N,P,**<br>**Q,R,S,**<br>**T,W,Y** |  | **K** | Unknown |
| M243 | **D,H,P,**<br>**W** |  |  | *C,I,L,V* | **A,E,F,**<br>**G,K,N,**<br>**Q,T** |  | **S,Y,R** | Unknown |
| P248 | **A,C,E,**<br>**G,R,S,**<br>**T** |  |  | *V* | **D,F,H,I,**<br>**K,L,Q,**<br>**W,Y** |  | **N,M** | Substrate binding (Rut et al. 2020; Shin et al. 2020) |
| Y264 | **H,M,R,**<br>**V** | F |  |  | **A,C,D,**<br>**E,G,I,N**<br>**,P,S,W** |  | **L,K,Q,**<br>**T** | Substrate binding (Zhao et al. 2021) |
| G266 | **L,M,N** | E |  | *C,D,F,*<br>*H,I,K,P,*<br>*Q,S,T,*<br>*V,W,Y* |  |  | **R**<br>**A\*** | BL2 flexibility |
| C270 | **P** | A,E,F,<br>G,H,I,K<br>,L,M,Q,<br>S,T |  |  |  | N | R,Y,V,<br>W<br>D* | second shell (Shao et al. 2023) |
| G271 | **F,H,P,**<br>**Q,R** |  |  |  | **A,C,D,**<br>**E,I,L,M,**<br>**S,T,V,**<br>**W,Y** |  | **N,K** | Substrate binding (Fu et al. 2021; Klemm et al. 2020) & blocking loop flexibility, second shell |
| H272 | **A,K** |  |  |  | **C,D,E,**<br>**F,G,I,L,**<br>**M,N,P,**<br>**Q,S,T,**<br>**V,W,Y** |  | **R** | Active site |
| Y273 | **C,H,I,W** |  |  | *F* | **A,D,E,**<br>**G,K,L,**<br>**M,N,P,**<br>**Q,R,S,**<br>**T,V** |  |  | Substrate binding(Zhao et al. 2021) |
| D286 | **Q** |  |  |  | **A,C,F,**<br>**G,H,K,**<br>**L,M,N,**<br>**R,T,W,**<br>**Y** |  | **S,V,P,I,**<br>**E** | Active site |
| D302 | **A,M,N,**<br>**S,T,V** |  |  | *C* | **E,F,G,I,**<br>**K,L,P,**<br>**Q,R,W,**<br>**Y** |  | **H** | Lines the S4 pocket<br>Unknown |

We depict these 22 residues on a PLpro structure that includes the LRGG PLpro cleavage motif taken from the PLpro:ubiquitin structure 6XAA (Klemm et al. 2020) (Fig. **3d**). Strikingly, mutations that impaired function without impacting abundance were not only found at the active site and substrate binding pockets along the interface of the thumb and palm domains, but also extended all the way up to the fingers domain in a connected pathway. The nature of these mutations and roles in PLpro function are reviewed in the Discussion.

### Predicting drug-escape variants with DMS

The blocking loop of PLpro (spanning residues 267-270) is flexible in its conformation, which likely allows substrates of various size to be accommodated (Báez-Santos, St. John, and Mesecar 2015). Our DMS data indicate that almost any mutation in the blocking loop is tolerated for function. Indeed, the MERS PLpro blocking loop differs in sequence and length, indicating that residues at these positions are not highly conserved (Lee et al. 2015). This property of PLpro may impact drug-discovery efforts as most reported inhibitors currently engage this flexible region (Brian Chia and Pheng Lim 2023; Calleja et al. 2022; Zhao et al. 2021). We therefore decided to employ our DMS screen to systematically investigate whether we could detect drug-escape variants to published PLpro inhibitors 3k, its analogue 5c (Báez-Santos, St John, and Mesecar 2015), GRL0617 (Ratia et al. 2008a), Jun9-84-3 (Ma et al. 2021) and XR8-89 (Shen et al. 2022).

We assessed each compound’s activity in our cellular assay (Extended data Fig. 5; Fig. **1f**) to determine an appropriate concentration with which to perform DMS, aiming for at least 80% inhibition. While XR8-89 had good activity in biochemical assays, we could only detect activity at 100 µM in cells, indicating this compound may be poorly membrane penetrant. GRL0617 had an EC50 of 37.3 µM, requiring more than 100 µM for an effective inhibition at EC80. At these high concentrations, GRL0617 starts to precipitate and impacts cell viability. Jun9-84-3 has a lower EC50 of 22.8 µM, but like GRL0617, we observed significant impacts on cell viability at high concentrations. Thus, we were only able to assess 3k and 5c in a physiologically relevant cell-based screen. We repeated the DMS screen in the presence of 65 µM 3k and 5c (EC75 - EC85), sorting FRET positive and FRET negative gates for comparison.

The resulting sequence-function maps were assessed for evidence of selection. If drug-treatment resulted in 100% inhibition of PLpro activity, we would see similar scores for synonymous wildtype and nonsense variants, as both variants would remain in the FRET positive gate. Instead, we saw that synonymous wildtype variants had higher scores than nonsense variants, consistent with incomplete inhibition (Extended data Fig. 6-7). Nevertheless, some positions showed elevated activity scores, and these were similar among the 3k and 5c datasets (Extended data Fig. 8). Because these compounds are almost identical (Extended data Fig. 5d), we combined the 3k and 5c datasets to gain precision and selected variants for individual testing that would be sensitive to the 3k/5c shared scaffold (Extended data Fig. 9).

After plotting activity scores ‘without compound’ against those ‘with compound’ (Fig. **4a**), we selected the following variants for testing in cells: wildtype-like variants L58V and Y268W (which was one of the few residues at position 268 that did not interfere with drug activity), escape variants D164S, D164C, M208A, M208W, and Y268R, as well as M208S, which differed in behavior between 3k and 5c. We transduced 293T sensor cells with each variant and tested activity before dox treatment (no PLpro), after dox treatment (PLpro alone), and with dox and 65 µM 3k (Extended data Fig. 10). Almost all inhibitor treatments behaved as expected, with L58V and Y268W remaining sensitive to compound and D164S, D164C, M208A and Y268R showing evidence of drug escape. M208S proved to also escape 3k, as seen in the 5c dataset, indicating that M208S provides an escape route for both compounds. The data from M208W, however, was unusual. The no dox treatment flow-cytometry plots indicated that this variant displayed substantial activity prior to dox-induced PLpro expression (Fig. **4b**). Indeed, 3k treatment inhibited dox-induced M208W PLpro back to pre-dox treatment levels (Fig. **4b**). This leads us to conclude that M208W was mis-characterized as a drug-escape variant in our screen because of poor dox control. To remove the effect of this artifact from our final list of escape variants, we repeated the screen without adding dox to characterize the landscape of leaky expression across our whole library (Extended data Fig. 11).

**Fig. 4.**
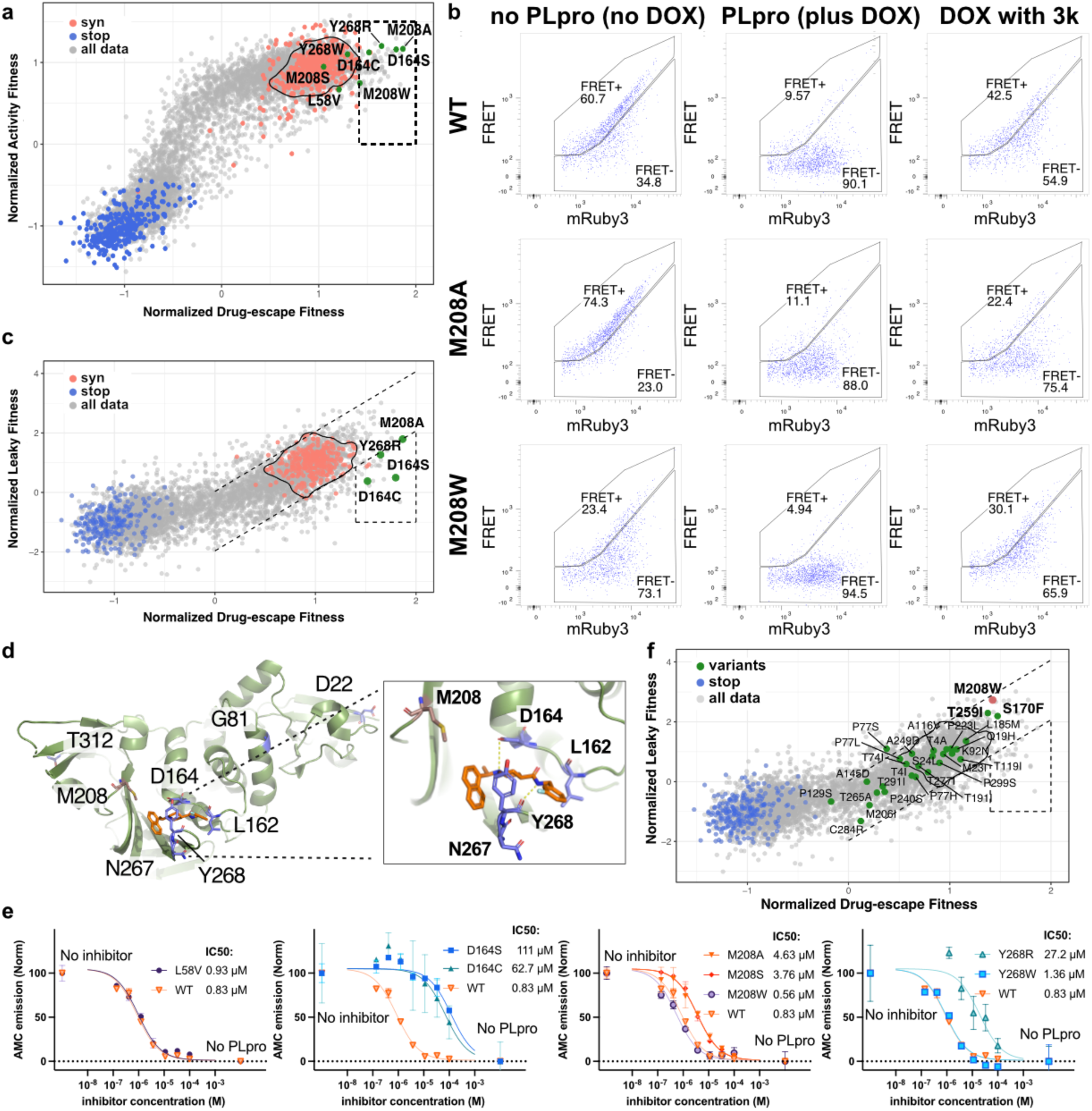
Identification of drug escape variants for the 3k/5c compound scaffold. **a)** A scatterplot of normalized activity scores against normalized 3k/5c combined drug-escape fitness scores (combined from five 5c and three 3k replicates from 5 independent transductions of the library; See *Extended data Fig. 9* for individual and combined drug-escape fitness sequence-function maps). Nonsense variants (positions 1-305) are shown in blue, synonymous wildtype variants in red, and the rest of the variants in grey. A density border around synonymous wildtype variants was calculated with ggplot2 function “geom_density2d” with a 0.2 contour encompassing approximately 90% of the datapoints. The dashed box contains variants with drug-escape fitness scores more than the main cluster of synonymous wildtype variants and activity scores similar to synonymous wildtype variants. Variants selected for downstream analysis are highlighted in green and labelled with their wildtype residue, position number and substitution. **b)** Representative flow cytometry plots of wildtype, M208A and M208W PLpro with 293T biosensor cells (transduced once, tested twice). Cells were measured without dox induction (no PLpro, left column), with dox treatment (PLpro, middle column) and with dox and 65 µM 3k (right column). The percentage of FRET positive and negative cells are shown. Wildtype PLpro displayed expected inhibition with 3k, M208A was resistant to 3k, M208W displayed poor dox control (leaky expression), but was unexpectedly sensitive to 3k inhibition. Other selected variants are shown in *Extended data Fig. 10*, were consistent with expectations. **c)** A scatterplot of normalized leaky (i.e., poor dox control) fitness scores (y-axis) against normalized 3k/5c combined drug-escape fitness scores (x-axis). See *Extended data Fig. 11* for leaky expression sequence-function map. All variants are shown in grey; nonsense variants (positions 1-305) are shown in blue and synonymous wildtype variants are shown in red. A density border around synonymous wildtype variants was calculated with ggplot2 function “geom_density2d” with a 0.2 contour encompassing approximately 90% of the datapoints. The dashed lines highlight the correlation between the two datasets. We identify variants below this correlation on the y-axis and above the cluster of synonymous wildtype variants on the x-axis (dashed polygon) as having good dox control and likely to be true drug-escape variants. Variants selected for downstream analysis are highlighted in green and labelled with their wildtype residue, position, and mutation. **d)** Positions identified in (c) as having drug-escape variants are highlighted in stick representation on the PLpro structure (PDB: 7TZJ) (Calleja et al. 2022). The zoomed view shows the compound 3k in orange and surrounding residues. Positions that have drug-escape variants and contact 3k are depicted in blue stick. M208, which is too far from 3k to make meaningful contact is highlighted in red stick. **e)** Recombinant protein was made for variants selected in (a) and tested against a dose response of 3k with the substrate Z-RLRGG-AMC, which becomes fluorescent upon cleavage. The data was scaled with AMC emission without PLpro defining 0% activity and AMC emission with no inhibitor defining 100% activity. The IC50s depicted in the legends were calculated from fitting dose response curves to normalized data with a hillslope of 1. A representative experiment is shown from 2 biological replicates. Error bars from technical triplicates indicate mean ± SD. IC50 values are the average of both biological replicates. **f)** The likelihood of drug escape in already circulating variants. Circulating variants (green) identified from COVID-3D (Portelli et al. 2020). were overlaid on the same scatterplot in (c). Synonymous wildtype variant datapoints were removed for clarity, but their border remains as a reference. Circulating variants are highlighted in green and labelled with wildtype residue, position and mutation. The M208W variant is highlighted in red and clusters near T259I and S170F indicating that these variants may be similarly stabilized.

We plotted leaky expression fitness scores versus drug escape fitness scores, allowing us to select variants with drug escape scores that were higher than synonymous wildtype variants and leaky expression scores that were in line with or lower than synonymous wildtype variants (Fig. **4c**; Extended data Fig. 12). The mechanism of drug escape for most variants can be explained by contacts to 3k (Fig. **4d**). However, some variants were far from the 3k binding site and may indicate noise in our dataset or an allosteric mechanism. The side chain of Met208 appears too far from 3k to make a meaningful contribution to compound binding, yet multiple mutations at this site affect 3k- and 5c-mediated inhibition without impacting PLpro function. This list of Met208 variants (Extended data Fig. 13-14) includes small side chain mutations, ruling out steric hindrance as a mechanism for drug escape.

### Activity and 3k responsiveness of recombinant PLpro variants

To determine the extent of drug escape, we elected to make recombinant PLpro variants and measure 3k dose-response curves to calculate IC50 scores for each variant. This orthogonal assay confirms that our DMS approach faithfully reports PLpro proteolytic activity and gives an indication of how sensitive our screen is to changes in IC50. It also provides control of PLpro concentration in subsequent assays, eliminating potential differences in expression as a source of drug-escape. We again selected L58V, D164S, D164C, M208A, M208S, M208W, Y268R, Y268W PLpro variants and expressed these variants and wildtype PLpro in *E. coli*. The yields of recombinant protein were similar to wildtype for variants at Leu58, Asp164 and Tyr268. M208A and M208S consistently gave yields around 3-fold lower than wildtype, while M208W yields were more than 3-fold higher.

The activity of each variant was tested in an assay that measured the cleavage of the commercially available fluorogenic substrate Z-RLRGG-AMC, which contains the PLpro recognition site and emits fluorescence upon cleavage. Unlike our cell-based assay, which measures biosensor cleavage at steady-state 24 hours after PLpro induction, this biochemical assay allows the initial rate of substrate cleavage to be measured for each variant and compared to wildtype. While all selected variants had activity scores similar to wildtype PLpro in our DMS assay, we saw impacts on the initial rate at which substrate was processed among variants: L58V and D164S and Y268W behaved similarly to wildtype, D164C and Y268R processed substrate at approximately 30% the rate of wildtype PLpro, and M208A and M208S processed substrate two and half times faster (Extended data Fig. 15). These results indicate that our DMS assay is powered to identify substantial, rather than mild, defects in activity that we predict will capture more physiologically relevant variants.

We tested each variant for its ability to be inhibited by 3k in the Z-RLRGG-AMC assay. Dose-response analyses were performed and IC50 values extracted from 2 independent experiments (Fig. **4e**). The IC50 of 3k against wildtype PLpro was 0.83 µM, while variants selected for wildtype-like activity L58V and Y268W exhibited IC50s of 0.93 µM and 1.36 µM, respectively. Y268R PLpro sensitivity to 3k was reduced, with an IC50 of 27.2 µM. D164S and D164C were the least responsive to 3k, with estimated IC50s of >100 µM and 62.7 µM, respectively. M208A and M208S showed modest loss of activity, with IC50s of 4.63 and 3.76 µM, respectively, while M208W gained ∼2-fold 3k sensitivity, with an IC50 of 0.56 µM. Together, these data show that we can identify drug-escape variants with as little as 4-fold loss in sensitivity to drug in our DMS analysis.

We identified circulating variants on our drug escape dataset, and none fell in the gate with which we define drug escape variants. However, T259I and S170F variants displayed leaky expression and clustered near M208W (Fig. **4f**).

### The impact of Met208 on PLpro structure and function

The position of Met208 in the PLpro structure is interesting in that it sits in the interface between the largely β-sheet-containing fingers domain and the α-helix-containing palm domain, most closely packing against Arg166, a position where we found several inactive variants without a structural explanation. Inspection of PLpro crystal structures with ubiquitin and ISG15 (Klemm et al. 2020; Wydorski et al. 2023) indicates that Arg166 may make polar contacts with ubiquitin residue Gln49 and ISG15 residue Asn151, but neither appears central enough to the interface to be critical for binding. Furthermore, it is unclear whether Arg166 would play a role in recognition of our cellular FRET-based biosensor. Ma et al., through docking simulations, predict that Arg166 may be mobile and able to create a salt-bridge with Asp164 (Ma et al. 2021). Asp164 is critical to substrate recognition, making contacts to the backbone of the P4 substrate residue (Leu) in the L(R/K)GG recognition motif that is shared among all PLpro substrates. Thus, perturbation of Arg166 may impair PLpro activity via impacts on Asp164’s key role in substrate recognition. Met208, while close to Arg166, sits back from the interface with substrate and does not make intimate contacts with ubiquitin, ISG15 or 3k (Extended data Fig. 16).

We decided to compare M208A and M208W PLpro Michaelis-Menten kinetics with wildtype PLpro against Z-RLRGG-AMC, Ub-Rhodamine and ISG15-Rhodamine substrates (Fig. **5a**). Surprisingly, M208A significantly boosted enzymatic activity of PLpro against Z-RLRGG-AMC, while decreasing Ub-Rhodamine and, to a lesser extent, ISG15-Rhodamine cleavage efficiency. M208W performed similarly to wildtype PLpro against each substrate. We were not able to deconvolute *k_cat_* from *K_M_* as affinity is predicted to be in the µM range, and we could not achieve these concentrations in our assays. Wishing to further explore the differences between M208A and M208W, we measured thermal stability (Fig. **5b**). M208A appeared similarly stable to wildtype PLpro, while M208W strikingly increases the protein melting temperature by over 5℃, indicating a substantial improvement in thermal stability. Increased stability, and thus reduced turnover in cells, may provide a mechanism to explain leaky expression in our cellular assay and increased yield of recombinant protein for *E.coli* expression.

**Fig. 5.**
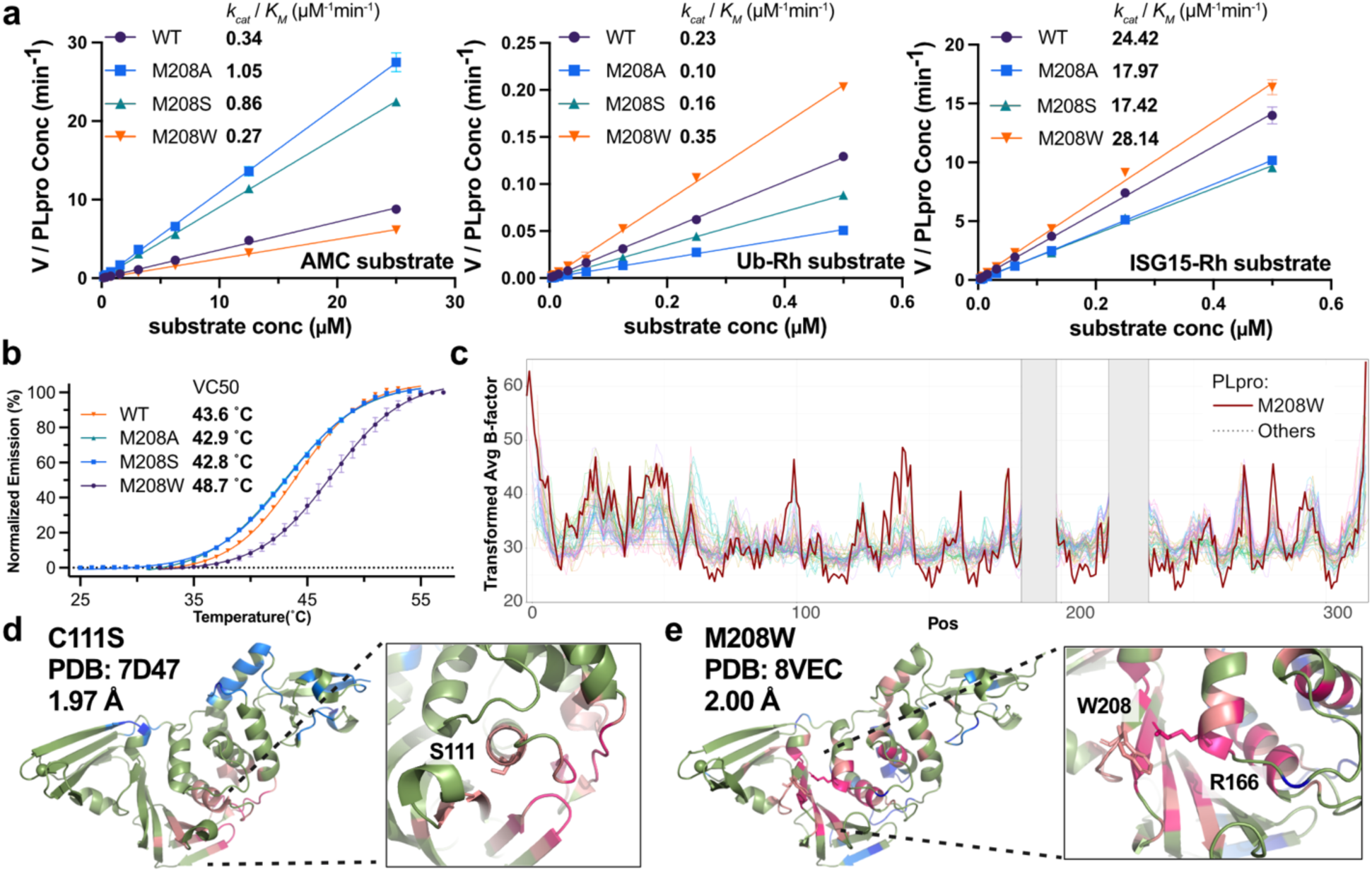
Characterization of M208 variants. **a)** Michaelis-Menten kinetics of M208 variants using substrates Z-RLRGG-AMC (left), Ubiquitin-Rhodamine (middle) and ISG15-Rhodamine (right). x-axis is substrate concentration in µM; y-axis is velocity of substrate cleavage divided by PLpro concentration used in the assay. The *k_cat_*/*K_M_* is approximated from the slope of the curves and makes the following assumption that the substrate concentration is substantially lower than the Km. Representative experiment from 2 biological replicates. Error bars denotes mean ± SD from technical replicates. *k_cat_*/*K_M_* is the mean from both biological replicates. **b)** M208W displays increased thermal stability. Wildtype, M208A and M208W variants were subjected to a Thermal Shift Assay that measures fluorescence from SYPRO orange dye as protein denatures with increasing temperature. Fluorescence at room temperature defines 0% denaturation and the plateau of each curve defines 100% on the y-axis. The midpoints are calculated with Boltzman Sigmoid Regression in Prism 9. Representative experiment from 2 biological replicates. Error bars denotes mean ± SD from technical replicates. The reported midpoints are the mean from both biological replicates. **c)** B factors from different 54 PLpro structures (light colors) scaled to the M208W PLpro structure (bold and red line). x-axis shows residue position and y-axis is the average of the backbone and Cβ atom B factors after scaling. The grey box selects out the zinc-coordinating regions which are disordered in most PLpro structures and were excluded from our analysis. **d)** C111S PLpro (PDB: 7D47) (unpublished) is stabilized at residues surrounding the active site. To test our B-factor analysis we applied the same process in (c) using PLpro C111S as the reference structure. The structure is presented in ribbon and colors indicate regions of stabilization that were less than 2 (pink) and 3 (magenta) standard deviations from the mean of all data and regions of destabilization that were more than 2 (light blue) or 3 (dark blue) standard deviations from the mean. The zoomed view shows the region surrounding C111S where stabilization is most prominent. e) M208W PLpro (PDB: 8VEC) (this study, *Supplementary Table* **1**) is stabilized at the interface of the palm and thumb domain. The structure is presented in ribbon and colors indicate regions of stabilization that were less than 2 (pink) and 3 (magenta) standard deviations from the mean of all data and regions of destabilization that were more than 2 (light blue) or 3 (dark blue) standard deviations from the mean. The zoomed view shows the region surrounding M208W and shows a potential role for R166 stacking against the introduced tryptophan at position 208. Both residues are represented in stick.

To determine whether M208A and M208W altered protein structure, we attempted to crystallize both proteins. Most crystal structures of PLpro contain inhibitor and/or bear the C111S mutation to improve crystal quality. We set up parallel crystallization screens with M208A, M208W and wildtype PLpro, all with intact catalytic sites. M208W readily crystallized (in 14/96 conditions), but we were unable to recover diffraction-grade crystals of M208A or wildtype PLpro. After minimal optimization of screening conditions, we collected a 2.00 Å dataset of M208W with unique space group and unit cell dimensions (P2_1_2_1_2; unit cell 58 Å, 85.5 Å, 83 Å, 90°, 90°, 90°) (*Supplementary Table* 1) compared to other PLpro structures in the PDB.

The resulting structure was overlaid with the crystal structure of C111S PLpro (PDB: 7D47, deposited without publication) and showed an identical global structure, with an RMSD of 0.495 Å over Cα atoms. The Trp at position 208 had clear electron density and was accommodated without disturbing neighboring residues. Overlay of the M208W structure with PLpro:Ub, PLpro:ISG15 and PLpro:3k structures indicated that this mutation does not introduce steric clashes with ISG15 or 3k (Extended data Fig. 16). Val70 of ubiquitin is within 2.5 Å of Trp208, potentially creating a mild steric clash. Nevertheless, M208W cleaved Ub-Rh more efficiently than wildtype PLpro indicating substrate engagement still occurs.

We examined B-factors to determine the extent to which the M208W mutation stabilized local PLpro structure. We collected all PLpro structures in the PDB with a resolution better than 3.5 Å and calculated the average of backbone and Cβ atom B-factors residue-by-residue along the length of the protein. Residues 186-197 and 219-232 were not considered during the analysis, as these regions have poor density in most reported structures. We plotted B-factor against residue number and calculated scaling factors to normalize the traces using a Broyden– Fletcher–Goldfarb–Shanno (BFGS) iterative algorithm (Fletcher 2000) from the “optim” function in R (R studio v4.2.0), which calculates optimal scaling and translation factors to minimize the differences between complex curves. We visually inspected the scaled plots for good overlay (Fig. **5c**). If successful, the scaling factors should also similarly normalize the overall Wilson B-factor, and this was the case (Extended data Table **2**).

We looked for local changes in stability as defined by regions where scaled B-factors were 2 (weak) or 3 (strong) standard deviations from the mean of all structures. We first assessed the C111S structure, predicting there would be observable stabilization around the active site. We colored regions of weak or strong stabilization (red tones) and destabilization (blue tones) on the C111S PLpro structure and saw stabilization around the active site as expected (Fig. **5d**). In contrast, increased stability in M208W PLpro extended along the whole β-sheet of the fingers domain and propagated into the α-helices of the palm domain that pack against this sheet (Fig. **5e**). The area of stabilization encompassed the 3k binding site and may provide a mechanism for improved 3k sensitivity by reducing flexibility without perturbing structure. While we were unable to determine the structure of M208A, we predict that Ala at this position would be destabilizing compared to methionine. We did not see this manifest in the thermal shift assay measuring the temperature at which PLpro denatures, but this prediction is consistent with lowered yield of recombinant protein when this variant was expressed in *E. coli*.

## Discussion

In this study, we comprehensively examined the mutational effects of every PLpro residue on the proteolytic activity and abundance of SARS-CoV-2 PLpro. Our data are validated by multiple observations. We see that mutation of the active site ablates PLpro activity, as does introduction of premature stop codons anywhere in the sequence except the last 10 residues. We also confirmed the role of the zinc-coordinating cysteines in the fingers domain as critical for protein integrity (Barretto et al. 2005), as variants at the four zinc-coordinating cysteines (with the exception of C224H) destroy proteolytic activity and reduce PLpro expression. By comparing activity and abundance datasets, we identified 22 positions where the function of PLpro is impaired without impacting structure. The nature of these positions is discussed below.

### Features of the Active site

PLpro is a member of papain-like thiol proteases superfamily. Papain itself is the model protein on which the catalytic mechanism of proteolytic cleavage can be understood for this family. Papain is globular protein with L- and R-domains. Papain shares very little sequence homology with PLpro, but when the structures are overlaid based on the arrangement of residues in the active site, parts of the L- and R-domains, and particularly the interface between them, overlay well on the thumb and palm domains of PLpro (Fig. **3b**). Like PLpro, the active site of papain is split between the L- and R-domains, with Cys25 on the L-domain and His159 found on the R-domain, which correspond to C111 and His272 in PLpro, respectively. The catalytic dyad formed by Cys25 and His159 is the minimal requirement for papain-mediated catalysis and results in a Cys^-^-His^+^ ion pair that acts as a nucleophile during protein cleavage. Three additional residues play prominent roles in catalytic activity. Asn175, often thought of as part of the catalytic triad, is not strictly necessary for catalysis, but rather plays a role in positioning His159 and modulating the ionization state of the catalytic dyad by influencing the pKa of His159 (Vernet et al. 1995). Trp177 also modulates the ionization state and plays a part in protecting the Asn175-His159 interaction from solvent (Gul et al. 2008). Glu19 stabilizes a transition state intermediate of the substrate during cleavage and forms part of the oxyanion hole that releases charge buildup during substrate cleavage (Ménard et al. 1991). In the PLpro active site, Asp286 plays the role of Asn175 in papain, Trp106 occupies the same relative position as the oxyanion-hole residue Gln19 in the papain active site, and there is no equivalent for papain residue Trp177.

Our DMS data confirm the absolute requirement for Cys111 and His272 in catalytic activity. Given that Asp286 is equivalent to Asn175 in papain, we were surprised that not even Asp is tolerated at this position in PLpro. Our alignments indicate that Asp is used exclusively in viral PLpro sequences. Indeed, D286N showed lower expression levels in our abundance DMS dataset, indicating that the explanation for this might be structural. Asp instead of Asn may also be preferred because of the lack of a PLpro residue analogous to Trp177, which shields the papain Asn175-His159 interaction from solvent.

PLpro residue Trp106 occupies the same position as oxyanion hole residue Gln19 and mutation in SARS-CoV PLpro impairs PLpro function (Baez 2012). While this Trp can be mutated to almost any other residue without affecting PLpro abundance, PLpro activity is only maintained with the substitution of polar residues Gln and Asn, other aromatic residues Tyr and Phe, and Ala. Although Trp is preferred, the nature of the mutations tolerated at this position is consistent with its ascribed role in stabilizing an oxyanion intermediate as suggested by Baez-Santos et al. in their analysis of SARS-CoV PLpro (Báez-Santos, St. John, and Mesecar 2015). MERS PLpro has a Leu at this position (Lee et al. 2015), which is unable to behave as a hydrogen-bond donor and our results indicate a similar mutation would impair SARS-CoV-2 PLpro. One study mutated MERS PLpro Leu106 to Trp and saw an increase in MERS PLpro activity, concluding that the properties of the oxyanion hole of MERS PLpro limit catalytic efficiency (Lei et al. 2014).

### Features of substrate binding

Our assays identify the residues required for cleavage of the Nsp2/3 site defined as L_P4_K_P3_G_P2_G_P1_↓A_P1’_. We identify features of substrate binding pockets and the sequence requirements of the blocking loop, which lines the narrow channel that feeds the P2 and P1 glycines of the substrate to the active site.

The blocking loop is flanked by Gly266 and Gly271. Both glycines are highly conserved among coronaviruses and Gly271 is completely intolerant of mutation in our system. Addition of a sidechain to Gly271 would impact blocking loop conformation and accommodation of the P2 Gly of substrate. Being adjacent to the catalytic His272 means perturbation of Gly271 would impact the active site. We thus see Gly271 is both functionally and structurally important. Gly266 variants are partially functionally impaired, but abundance is not reduced, indicating a likely role for flexibility of the blocking loop for substrate binding. The sequence between Gly266 and Gly271 is tolerant to almost any mutation, indicating that movement of the blocking loop is likely required, but the sidechains of the blocking loop are dispensable for substrate binding and PLpro activity.

The S4 pocket that binds the essential P4 Leu is larger than what is needed to accommodate Leu and has been exploited for drug-discovery purposes instead of the shallow active site. The S4 pocket is lined by Asp164, Arg166, Met208, Pro247, Pro248, Tyr264, Tyr273, Thr301 and Asp302; but only Asp164, Pro248, Tyr264 and Tyr273 and Thr301 are in close proximity to the P4 Leu. The blocking loop flips up and, with Leu162, forms a roof over the S4 pocket and the P3 residue of the substrate. Among the residues near the P4 Leu, we find functional roles for all except Thr301. We also see a functional role for Arg166 and Asp302, despite their distance from the site of P4 engagement, that cannot be explained by a structural impairment in our datasets. The importance of these residues towards substrate binding have been described elsewhere (Rut et al. 2020; Shin et al. 2020; Klemm et al. 2020; Fu et al. 2021; Perlinska et al. 2022; Ratia et al. 2008a; Shiraishi and Shimada 2023).

We also identified that Arg183, Val184, Val242, Met243, and Tyr207 are important for PLpro function, and with Arg166 and Asp302, they form a connected pathway from the back of the fingers domain all the way to the S4 pocket. Arg166 has been proposed to move upon substrate binding and stabilize Asp164, a key residue for substrate binding (Ma et al. 2021). It is possible that these residues, with Met208 (see below), play an allosteric role in modulating PLpro engagement with substrate.

### Escape from inhibition of the 3k/5c scaffold

PLpro is a validated drug target for COVID-19 treatment (Klemm et al. 2020). We took advantage of our robust mammalian cell DMS platform to explore how PLpro might evade inhibition from the 3k/5c scaffold, which were the only reported compounds we found that could be used sensibly in our cellular assay. In doing so, we provide the framework for assessing other more potent lead compounds as they emerge. SARS-CoV-2 PLpro has an active patent landscape with 45 patents (Brian Chia and Pheng Lim 2023) filed since the start of the pandemic. We attempted to measure resistance to GRL0617, Jun9-84-3 and XR8-89 but could not achieve high enough concentrations in our cellular assays without encountering toxicity and precipitation issues. We therefore limit the findings reported here to the two related compounds 3k and 5c.

The PDB contains many entries of PLpro in complex with various inhibitors, and almost all contact Tyr268 on the blocking loop. Our DMS datasets indicate that this residue is dispensable for PLpro proteolytic activity, although our experiments with recombinant PLpro measuring initial rates of substrate cleavage indicate that Y268R activity is substantially lower than wildtype. We were able to correlate our DMS Activity scores to viral fitness through assessment of circulating variants: C284R has been observed in patient samples, and raw DMS activity score indicates an approximately 15-fold reduction in activity compared to wildtype. Although rare in SARS-CoV-2 circulating variants, SARS-CoV PLpro contains an Arg at this position, providing support for this being a genuine circulating variant and indicating that even poor PLpro activity can result in functional virus.

Our initial drug escape data highlighted that variants at Met208, Tyr268 and Asp164 are important to drug binding. We validated several variants at these sites and were able to confirm all but one indeed resulted in drug escape. M208W was the exception, and we identified that poor dox control contributed to its misidentification. In response, we created a map of variants that had higher than expected expression in the absence of dox and surmise that these variants will either have increased catalytic activity or increased protein stability.

We were then able to exclude variants with high leaky expression to isolate true drug escape variants. Because several filters have been applied to generate our final datasets, and some variants are missing from the analysis, we do not claim that this is a complete list of drug escape variants. Nevertheless, of the variants we can confidently define, most are found with mutations at Tyr268 and Asp164, both of which directly contact the 3k scaffold: Tyr268 via a T-shaped π stack against the 3k naphthalene and Asp164 through a hydrogen bond to the piperidine nitrogen (Calleja et al. 2022). Aromatic residues at Tyr268 were found to retain sensitivity to drugs, while other mutations provide escape. At Asp164, only Cys, Ser, Asn and Gln variants are functionally tolerated, and of those, D164C and D164S cause drug escape.

The DMS also revealed Met208 as being important to 3k and 5c activity. The reason for this was unclear, since Met208 does not directly contact either inhibitor, with the terminal Cε atom of Met208 being 4.4 Å from the nearest carbon atom of the naphthyl ring of 3k (PDB: 7TZJ) (Calleja et al. 2022). By inspecting the crystal structure of PLpro with ubiquitin (6XAA) and full length ISG15 (7RBS) (Klemm et al. 2020; Wydorski et al. 2023), we conclude that Met208 also does not play an essential role in these interactions. Our functional DMS data additionally shows Met208 to be tolerant of almost any other substitution. Other coronaviruses have Phe and Ala at this position, which further demonstrates that Met208 is neither conserved nor required for proteolytic, deubiquitination or deISGylation activities. We confirmed M208A and M208S reduced drug sensitivity in both cellular and biochemical assays with recombinant protein, but we observed that M208W was sensitive to drug, which excludes a role for steric hindrance as a mechanism for drug escape. Instead, we propose that thermal motion around the S4 pocket is important. M208A and M208S both increased proteolytic activity of PLpro while modestly decreasing the rate of ubiquitin or ISG15 cleavage. M208W behaved similarly to wildtype PLpro in assays measuring catalysis, but it promoted thermal stability as measured by an increase of 5℃ of the PLpro melting temperature, and it readily crystallized, which through B-factor analysis showed stabilization of an area encompassing large parts of the palm and thumb domains. Together, mutations at Met208 suggest that there is role for thermal motion in substrate preference and may explain why mutations distant from the active site, and extending to the fingers domain, impact PLpro activity.

### Concluding Remarks

Similar to work performed by Bolon and colleagues (Flynn et al. 2022), who determined the fitness landscape of Mpro, we applied DMS to decipher the mutational landscape of PLpro and deepen our understanding of the biology of this crucial viral protease. Our results further provide knowledge of the potential escape routes open to the virus to evade developing therapeutics. Plasticity in the S4 binding pocket and blocking loop presents a hurdle that should be considered during the drug development process. This therefore benefits current and future efforts in developing PLpro inhibitors and opens the door for other research into the catalytic mechanism of PLpro.

## Supporting information

Supplementary Tabel 3

Supplementary Tabel 3

R script for B factor analysis

R script for PLpro PacBio data analysis

## Acknowledgements

This work has been supported by: a Wellcome Trust Innovator Award 222698/Z/21/Z, WEHI Innovator funding, MRFF grants MRF2002119 and MRF2016781 to DK, GL, MJC among others; NHMRC Investigator Grants (GNT2016461 to GL, GNT1178122 to DK); a donation from Hengyi Pacific Pty Ltd to support COVID-19 research; a donation from AWM Electrical to support Australian drug discovery research; and a donation from John and Tibby Peterson to MJC.

This research was undertaken in part using the MX2 beamline at the Australian Synchrotron, part of ANSTO, and made use of the Australian Cancer Research Foundation (ACRF) detector. The DMS libraries were created with the help of the WEHI Multiplexed Assay Technology Hub, which was founded with WEHI New Medicines and Advanced Technology Theme funding. We would like to thank the Bio21-WEHI Crystallisation Facility within Melbourne Protein Characterisation at The Bio21 Molecular Science and Biotechnology Institute, The University of Melbourne. We thank WEHI Flow Cytometry and Advanced Genomics Facilities as well as the National Drug Discovery Centre for technical resources and advice.

All DMS data have been provided in the Supplementary Materials as Resource files and have been deposited on MaveDB (mavedb:00000672). The crystal structure of M208W PLpro has been deposited to the PDB (8VEC). Work in the laboratories of the authors was made possible through Victorian State Government Operational Infrastructure Support (OIS) and Australian Government NHMRC Independent Research Institute Infrastructure Support (IRIIS) Scheme.

## Author Contribution

MJC conceived and with MEC and DK supervised the study. XW and MG designed the libraries. MJC and XW designed the cell assays and DMS strategies. XW, MG prepared the samples and carried out the Illumina sequencing. XW and MG prepared the samples for PacBio sequencing. All data were processed and analyzed by XW, under the guidance from MJC and MEC. XW performed all cell experiments, and with BGCL performed the biochemical assays. XW purified all proteins and with JVN performed the crystallization study. DJC provided advice on PLpro protein purification and crystallization. MJC and XW processed and analyzed the crystallization data. DK, GL, JPM, BGCL, KNL and NWK provided PLpro inhibitors and developed biochemical assays used in this study. XW and MJC drafted the manuscript, which was reviewed and edited by all co-authors.

## Materials and Methods

### Vectors

The FRET biosensor and wildtype SARS-CoV-2 PLpro coding sequences were ordered as G-blocks (IDT) and subsequentially installed into the Gateway donor vector pDONR221 (Thermo Fisher Scientific) with BP clonase (Invitrogen).

The FRET biosensor was shuttled into pMX-Gateway-IRES-Hygro (gift from Andrew Brooks, University of Queensland) with LR clonase (Invitrogen).

The PLpro recipient vector, FU-tetO-Gateway-rtTA-2A-Puro (Extended data Fig. 2c), was constructed from FU-tetO-Gateway, a gift from John Gearhart (Addgene plasmid #43914; http://n2t.net/addgene:43914; RRID:Addgene_43914), pMSCV-puro (Dickins et al. 2005) and pFTRE3G (Murphy et al. 2013). Briefly, the PGK driven Puromycin resistance cassette from pMSCV-puro was PCR amplified with oligos containing XmaI sites. The PCR product was cut with XmaI and ligated into AgeI cut FU-tetO-Gateway to generate FU-tetO-Gateway-Puro. The rtTA-2A-Puro was then installed by cutting FU-tetO-Gateway-Puro and pFTRE3G with AgeI and BsiWI. The final vector, FU-tetO-Gateway-rtTA-2A-Puro, was created by ligation of the vector fragment from FU-tetO-Gateway-Puro and the insert fragment of pFTRE3G. The pOPIN-B vector was used for recombinant PLpro expression and was a gift from Ray Owens (Addgene plasmid # 41142). All oligos used in this study were ordered from IDT. All individual PLpro mutations were introduced into pDONR221-PLpro using NEB Q5 Site-Direct Mutagenesis Kit (NEB) following manufacturer’s instructions and shuttled into FU-tetO-Gateway-rtTA-2A-Puro using LR clonase.

We also constructed pFGH1-UTG-mTagBFP2 as empty vectors by inserting PCR-amplified mTagBFP2 from pLKO.1 - TRC (a gift from Timothy Ryan; Addgene plasmid #191566) into digested backbone from pFGH1-UTG, a gift from Marco Herold (Addgene plasmid #70183) via HiFi assembly. This empty vector is used for lentivirus co-package for PLpro DMS.

### Cell line

Verified HEK293T cells were sourced from Cellbank Australia, (#12022001) and maintained in DMEM (GIBCO, #10313039) supplemented with glutamine and 10% FBS (GIBCO, #2526728RP).

### Retroviral and Lentiviral production and transduction

HEK293T cells were transfected with transfer vector and retroviral packaging vectors MMLV-gag-pol and VSVg or lentiviral packaging vectors RSV-REV, pMdi and VSVg with Calcium Phosphate. Supernatant was harvested 2 days later and filtered through a 0.45 µm filter. Polybrene was added to the viral supernatant to a final concentration of 4 µg/ml. Virus was diluted in complete DMEM + polybrene to achieve the desired MOI and added to HEK293T cells at 50% confluency. Cells were spun at 37°C for 45 mins at 700g to encourage transduction before overnight incubation at 37°C. Media was replaced the following day and cells assessed or placed on selection two days post transduction.

### Generation of HEK293T biosensor cells

pMX-mClover3-TLKGGAPTKV-mRuby3-IRES-Hygro was packaged in HEK293T cells with MMLV-gag-pol and VSV-g. HEK293T cells were transduced at a multiplicity of infection (MOI) of approximately 0.1 to avoid multiple integrations. Two days after transduction, the cell line was subjected to 180ng/ul Hygromycin (Merck, US1400052) treatment for 2 days to remove non-transfected cells and sorted twice based on medium FRET signal. The resulting cell line was aliquoted in cryoprotective medium containing 90% FBS + 10% DMSO and subsequentially stored in liquid nitrogen. A fresh batch of cells was thawed for every new experiment. The same procedure was repeated for biosensors with SARS-CoV and MERS PLpro cleavable linkers.

### Introduction of PLpro into HEK293T biosensor cells

FU-tetO-PLpro-rtTA-2A-Puro was packaged in HEK293T cells with RSV-REV, pMdi and VSVg. Retrovirus was transduced into HEK293T biosensor cells at a multiplicity of infection (MOI) of approximately 0.1 to avoid multiple integrations. Two days after transduction, the cell line was subjected to 2 ng/µl Puromycin (Thermo, #A1113803) treatment for 2 days to remove non-transfected cells. The resulting cell line was aliquoted in cryoprotective medium containing 90% FBS + 10% DMSO and subsequentially stored in liquid nitrogen. A fresh batch of cells was thawed for every new experiment. The same procedure was repeated for SARS-CoV and MERs PLpro.

### Inhibitor dose response assay

Wells of a 96-well flat-bottom plate were seeded with 2.5 x 10^5^ cells in 150 µl media. Compound was serially diluted in DMSO in a 7 point, 3-fold dilution starting at 10 mM. 1.5 µl each dilution was added to the appropriate wells and 300 ng/ml dox (#D5207, Sigma Aldrich) added to induce PLpro expression. Each compound was tested in triplicate. In every drug screen, 5c was used as benchmark. After overnight incubation at 37°C, 10% CO_2_, cells were detached and analyzed by flow cytometry (WEHI FACS facility) to determine the FRET+ percentage of cells at each inhibitor concentration.

To fit dose-response curves non-linear regression was used to fit the data from 5c using the following equation in Prism 9. The Hill slope was set to 1 and constants determined for the Top and Bottom of the curve:

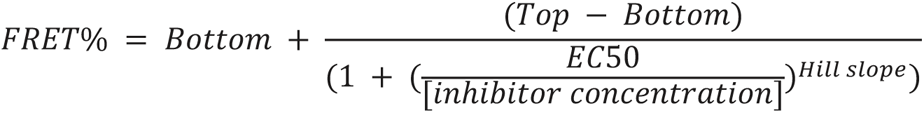

The data from all compounds in each experiment was then normalized by calculating the scaling factor (*k*) and translating factor (*b*) and correcting FRET+ % to PLpro activity using the following equations.

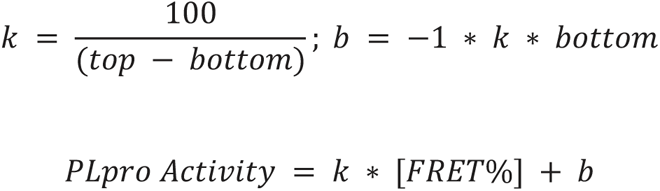

For other compounds non-linear regression performed on normalized data to determine EC50s with the top set at 100 and bottom as 0. Hill slope was varied from 1 as required.

### PLpro library construction

For residues close to the active site (encompassing residues 62, 69-70, 73-74, 77, 93, 104, 106-119, 151-174, 206-212, 243-253, 260-276, 285-286, 296-304) we ordered dsDNA with gateway adapters from Twist and directly cloned these fragments into our lentiviral vector FUV-tetO-Gateway-rtTA-2A-Puro vector. For the remaining residues, we adapted the MITE mutagenesis from (Melnikov et al. 2014) (Extended data Fig. 2a). The library was broken up into 6 cohorts of approximately 40 residues each. Three fragments were cloned in a single HiFi Assembly reaction (NEB). 1) The vector sequence was PCR amplified from pDONR221-PLpro and spanned a region 3’ to the barcode (installed 3’ to the PLpro stop codon) to 5’ to the region targeted for mutagenesis. 2) A PCR product was created that spanned 3’ to the region targeted for mutagenesis to the end of PLpro and contained 16 bp barcode. 3) The region targeted for mutagenesis as ordered as an IDT oPool and contained NNK variant codons. All fragments had overlaps of ∼25 bp to facilitate HiFi Assembly. The Assembly reactions were transformed into DH-10b electro-competent cell (NEB) and split into 2% and 98% before spreading on Agar plates containing 50ng/ul Kanamycin. Colonies on 2% plate were counted manually and used to estimate the total number of transformants. Collected transformants were determined by the cohort size aiming for approximately 10 colonies per DNA variant. The cohorts were moved to FU-tetO-Gateway-rtTA-2A-Puro by Gateway LR cloning (Invitrogen). All cohorts and the Twist library were pooled at equal molarity.

### PLpro Abundance library construction

We shuttled the PLpro library back to pDONR221 with BP clonase to create a complete library in the pDONR221 backbone. We designed a PLpro-mClover3 fusion separated by the linker PVGGSGGGSGGG and ordered this as an IDT G-block and cloned it into pDONR221 with BP clonase (Invitrogen) to be the basis of our new abundance library.

To construct this library, we cut our existing PLpro library (in the pDONR221 backbone), with PpuMI (at G298) and XmnI (between the PLpro stop codon and barcode) and recovered the vector fragment. Meanwhile, we PCR amplified the pDONR221-PLpro-linker-mClover3 with oligos annealing to the region 25bp upstream G298 and the region 25bp downstream of the stop codon. The PCR product was mixed with the vector fragment and assembled with HiFi assembly (NEB). This strategy maintains variant barcode matching from residues for the library up until L290. We then created new variants between L290 and K315 with an additional round of MITE mutagenesis as described in the previous section, but now using pDONR221-PLpro-linker-mClover3 as the source of vector.

### PacBio characterization of DMS Libraries

The PLpro library was cut with NotI and NdeI (NEB) to release a 3908bp fragment containing PLpro and the barcode; For the PLpro abundance library, we used NotI and BlpI (NEB) to generate a 3841bp fragment. Both fragments were gel purified and isolated with a ZymoClean Gel extraction kit (Integrated Science). The samples were lyophilized and sequenced on the Revio system at AGRF (University of Queensland).

We obtained approximately 4 million high-fidelity consensus reads for the PLpro library and approximately 2 million reads for the PLpro abundance library. After trimming to the region of interest and correcting orientation of reverse reads with cutadapt version 3.4 (Martin 2011), reads were further filtered with cutadapt to remove reads with greater than 100 mutations. Then barcode error correction was performed with UMI tools version 1.1.4 (Smith, Heger, and Sudbery 2017) with an error-correct-threshold setting of 2 and only sequences with more than one read were kept. To generate the barcode look up table, the barcode corrected fastq file was loaded to R studio v4.2.0 with readfastq from the “ShortRead” pacakge (Morgan et al. 2009). The reads were separated by barcode and global pairwise alignment (Coghlan 2011) was performed. If 50% or more of the reads for any given barcode had an insertion or deletion at the same position, that barcode was discarded. For a variant to be called for a particular barcode, greater than 90% of the reads had to contain the same mutation. Barcodes attached to reads without mutations were classified as wildtype. Barcodes with reads that didn’t meet these criteria were discarded. Finally, barcode lookup tables for both libraries were constructed from the reads that passed the criteria. The linker and mClover3 sequences were also checked during PLpro abundance library data processing. We identified 102,363 and 31,698 unique barcodes for the PLpro and PLpro abundance libraries respectively. The source code has been provided in the supplementary materials.

### DMS Library Selection

To ensure cells received a single copy of PLpro we mixed the libraries with an unrelated lentiviral vector pFGH1-UTG-mTagBFP2 at a 1:2 ratio. This strategy has previously been reported and minimizes barcode swapping during viral production and transduction (Hill et al. 2018). Virus containing the PLpro library was then transduced into enough HEK293T biosensor cells, or in the case of the mClover3-containing abundance library, parental HEK293T cells, to ensure 10-fold replication of barcodes at an MOI of approximately 0.2. Two days after transduction, transduced cells were enriched in 2 ng/µl Puromycin (Thermo, #A1113803) for two days. Live cells were then harvested and seeded for respective selections.

For the activity screen, we treated cells with 300 ng/ml dox for 24 hours and sorted FRET positive from FRET negative cells for mRNA extraction (Fig **1g**). For the abundance screen we similarly treated with dox but sorted on mClover3 high and mClover3 low cells (Extended data Fig. 4). To select for drug-escape variants, cells were similarly treated with dox and 3k and 5c were added at to a concentration of 65 µM corresponding to an effective concentration of ∼80%. 24 hours later cells were sorted from the FRET positive and FRET negative gates for mRNA extraction. To measure the efficiency of dox control (leaky expression), cells were not treated with dox prior to sorting from the FRET positive and FRET negative gates but were instead treated with dox for 4 hours after sorting to induce mRNA for extraction.

### mRNA extraction and Next-Generation Sequencing

RNA was extracted with RNeasy Mini Kit (Qiagen) following the manufacturer’s instructions. The RNA was reverse transcribed with a primer that annealed 3’ to the barcode and contained an overhang adapter for index primer annealing. After reverse transcription the cDNA was amplified by a second overhang primer annealing 5’ to the barcode and 3’ to the PLpro stop codon and a reverse overhang primer. The resulting PCR product was indexed in triplicate using the flanking overhangs to allow multiplexed sequencing. The indexed samples were sent for NextSeq 2000 (illumina), single-end sequencing at the WEHI Genomics Hub. The average reads per barcode for each sample was typically more than 30.

### Illumina Data Processing and variant scoring

The data were demultiplexed and trimmed with cutadapt version 3.4 (Martin 2011). The final data consisted of fastq files containing barcodes representing the frequency of each variant’s mRNA in the selected population. Variant fitness scores from various selection conditions (see below) were calculated with DimSum (Faure et al. 2020) using Illumina-derived barcode fastq files in combination with PacBio-derived barcode lookup tables to deconvolute barcodes into variants. The quality filter was set at 15 and no minimum read number was set. Technical replicates were used to combine samples with triplicate indexes after checking for concordance. Experimental replicates were defined as selections from independent transductions. DimSum automatically normalizes scores such that wildtype is 0. Instead, we wanted to normalize data to center the wildtype to a score of 1 (F_WT_) and the population of nonsense variants to a score of −1 (F_avg-nonsense(res1-305)_) for easy visualization of variants and comparison of datasets. To accomplish this, raw DimSum fitness scores F_raw_ were normalized with a scaling factor (*k*) and translation constant (*b*) according to the following equations.

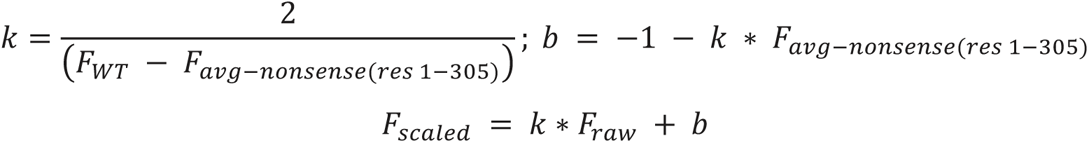

Errors were scaled with *k*.

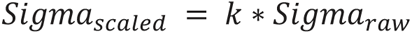

### Activity scoring

Samples from the FRET positive gate (inactive PLpro) were treated as time-point 0 in DimSum, while those from the FRET negative gate (active PLpro) were treated as timepoint 1. After normalization, data was filtered to remove variants with no more than 10 reads at time-point 0 and sigma no less than 0.4 (Extended data Fig. 3). Positive fitness scores indicate activity and negative fitness scores indicate impairment.

### Abundance scoring

Samples from the mClover3 low gate (misfolded PLpro) were treated as time-point 0, while those from the mClover3 high gate (folded PLpro) were treated as timepoint 1. After normalization, data was filtered to remove variants with no more than 13 reads at time-point 0 and sigma no less than 0.75 (Extended data Fig. 4). Positive fitness scores indicate expression and negative fitness scores indicate impaired expression.

### 3k and 5c drug-escape scoring

Samples from the FRET positive gate (sensitive and functionally impaired PLpro) were treated as time-point 0, while those from the FRET negative gate (insensitive PLpro) were treated as timepoint 1. After normalization, data was filtered to remove variants with no more than 5.5 (3k) and 9 (5c) average reads and errors no less than 0.5. Fitness scores greater than 1 indicate reduced sensitivity to drug, while scores below 1 indicate inhibition and/or impaired PLpro activity.

### Characterizing each variant’s leaky expression (poor dox control) profile

Samples from the FRET positive gate (dox-controlled or inactive PLpro) were treated as time-point 0 in DimSum, while those from the FRET negative gate (poor dox control) were treated as timepoint 1. After normalization, data was filtered to remove variants with no more than 18 reads at time-point 0 and sigma no less than 1 (Extended data Fig. 11). Positive fitness scores indicate leaky expression and negative fitness scores indicate good dox control or impaired PLpro.

### Calculation of Sequence-Function maps

Sequence-Function maps with blue-white-red gradients representing scores and lines representing errors were prepared in R studio v4.2.0 with ggplot2. For activity scores the white point was set to 0 (midway between synonymous wildtype and nonsense variant scores). For abundance data the white point was set at 0.5 to bisect the two main peaks of the library distribution. For drug-escape and leaky expression scores the white point was set at 1 (centered on synonymous wildtype scores). The code can be provided upon request.

### Identification of functionally important variants

We plotted normalized abundance scores against normalized activity scores to find functionally important variants that remained abundant but had impaired activity. Based on the distribution of wildtype and nonsense variants with synonymous mutations, we classified variants with abundance scores exceeding 0.5 and activity scores below −0.4 as functionally important. Given the proximity of many variants to the threshold and variations in error magnitudes, we additionally mandated that variants adhere to these criteria even after adjusting their fitness scores by subtracting (for abundance) or adding (for activity) error (*Sigma_scaled_* as above). Variants passing both criteria are colored green in Fig. **3c**.

To assess other variants at functionally important positions we further gated this plot as indicated in Extended data Fig. 17, grouping remaining variants into 5 categories: low abundance, but active; abundant and active (wildtype-like); partially abundant and partially active; abundant, but partially active; and low abundance, but active. The results of this are reported in Table **1**.

### Identification of 3k and 5c drug escape hotspots

We first attempted to extract drug-escape variants from a plot of drug escape scores versus activity scores, however in follow up experiments we found our dataset was confounded by poor dox control of some variants. We instead identified drug-escape variants by plotting drug-escape scores versus leaky expression scores. We noted disperse populations for synonymous wildtype and nonsense variants in these plots and performed additional cleanup of the data (Extended data Fig. 12). Variants defined with less than 5 barcodes were discarded as were rare variants with much higher errors than the main population. When these filters were applied to synonymous wildtype variants the population dispersion was considerably narrowed. We used surviving synonymous wildtype variants to determine a final minimum counts filter that was applied to the whole dataset.

The filtered data was then re-plotted (drug-escape vs leaky expression) to identify drug-escape variants that were defined as follows. We drew a density border around synonymous wildtype variants using geom_density2d from ggplot2 with a 0.2 contour cutoff. At the top and bottom borders of this contour we extended the linear relationship between drug-escape and leaky expression (Fig. **4c**). Variants the had drug-scape scores greater than the right contour boundary of the synonymous wildtype variants and were below the bottom linear correlation line were considered true escape variants.

### Recombinant PLpro expression and purification

The procedure was adapted from (Calleja et al. 2022). Wildtype PLpro and variants were transformed into BL21/DE3 (NEB) and grown in Super Broth (WEHI Bioservices) with 50 µg/ml Kanamycin. Once an optical density of 0.8 was reached, the temperature was dropped to 18°C and expression was induced with 300 µM IPTG. Cells were harvested 16h-18h after induction and pelleted. Pellets were resuspended in lysis buffer (50mM Tris.HCl pH 7.5, 500 mM NaCl, 10 mM Imidazole pH 8.0, 5 mM beta-mercaptoethanol, freshly supplemented with lysozyme, DNaseI and cOmplete protease inhibitor cocktail (Roche)) and sonicated. Supernatants after high-speed centrifugation were loaded onto His-Tag Purific Resin (Roche), washed and eluted with 50 mM Tris.HCl pH 7.5, 500mM NaCl, 300mM Imidazole pH 8.0, 5mM beta-mercaptoethanol. The eluate was desalted with a PD-10 column (GE healthcare) into 50 mM Tris.HCl pH 7.5, 500 mM NaCl, 10 mM Imidazole pH 8.0, 5 mM beta-mercaptoethanol and cleaved with 3C protease overnight. The next day, protein was again passed over Nickel resin to remove His-tagged protein and further purified by size exclusion chromatography (Superdex 75 10/300g, GE healthcare) into 20mM Tris.HCl 7.5, 50mM NaCl, 1mM TCEP. Fractions were analyzed with SDS-PAGE and correct fractions were pooled, concentrated, aliquoted, flash-frozen and stored at −80°C.

### Recombinant PLpro Activity Assays

We used the PLpro substrate Z-RLRGG-AMC acetate (Sigma Aldrich) for inhibitor dose response assays. Inhibitors were prediluted in DMSO at 50-fold their final concentrations and tested in a 10-point, 3-fold dilution series with 100 µM as the top final concentration. 120 nL of inhibitor was first spotted in wells (Echo^®^ acoustic dispenser, LabCyte) of a 384-well black plate (Corning #3820) and PLpro and Z-RLRGG-AMC added to initiate the reaction (6 µl of 10 nM PLpro and 300 nM Z-RLRGG-AMC in 50 mM HEPES pH 7.5, 0.1 mg/ml bovine serum albumin, 150 mM NaCl, 2.5 mM dithiothreitol). The mixture was incubated at room temperature for two hours. AMC fluorescence with excitation at 380 nm and emission of 445 nm was measured on a ClarioStar Plus (BMG labtech).

For *k_cat_*/*K_M_* measurements, an 8-point, 3-fold serial dilution of Z-RLRGG-AMC starting at 25 µM was incubated with 5 nM PLpro in the same assay buffer as above (reaction volume 6 µl). To measure the fluorescence of completely cleaved substrate we incubated the same Z-RLRGG-AMC dilution series with 100 nM PLpro. Base measurements of the dilution series in the absence of PLpro were also taken to measure the fluorescence from un-cleaved substrate. The reaction was scanned every 5 min for 2 h to measure initial rates. The same strategy was used for Ubiquitin-Rhodamine (Ub-Rd110Gly, UbiQ) and ISG15-Rhodamine (ISG15-Rd110Gly, UbiQ) substrates with the following alterations. The dilution series began at 500 nM, the assay buffer was 20 mM Tris pH 8.0, 0.03% BSA, 0.01% Triton X-100, 1 mM GSH and fluorescence was measured at an excitation of 487 nm and an emission of 535 nm. 5 nM PLpro was used in the Ub-Rhodamine assay while 100 nM PLpro was used to capture the complete cleavage fluorescence; 0.5 nM PLpro was used in the ISG15-Rhodamine assay while 10 nM PLpro with for ISG15-Rhomdaine was sufficient for complete cleavage in 2 h. As the substrate concentrations used in these assays were well below the estimated *K_M_* for each substrate, we approximated the *k_cat_*/*K_M_* from the slope of plots of initial rate/PLpro concentration vs substrate concentration with linear regression. A similar approach has been used previously (Rut et al. 2020)

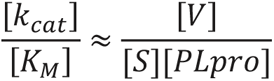

The equation derived from the conventional Michaelis-Menten equation: 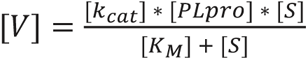. As [*K_m_*] >> [*S*], the denominator was approximated to [*K_m_*].

### Thermal Shift Assay

20 nM PLpro in 20 mM Tris-HCl, 50 mM NaCl, 1 mM TCEP with 4% SYPRO Orange dye (Merck, #S5692) were filtered in SpinX tubes (Corning) and 25 µl added to a 384-well PCR plate (Thermo, AB-1384/W). Samples were heated for 30s at the temperature starting at 25°C to 95°C by 1°C increment and SYPRO Orange fluorescence monitored with a Biorad CFX384 Real Time System, C1000 Thermal Cycler. Data were normalized with initial fluorescence dictating 0 and maximum fluorescence as 100. Melting temperatures (V50) were extracted with regression using the following the formula in Prism 9: Y=Bottom+(Top-Bottom)/(1+exp((V50-X)/Slope)).

### M208W PLpro structure determination

PLpro M208W protein was concentrated to 10 mg/ml and sent to the WEHI-Bio21-Crystallisation Facility for SG1 (shotgun) screening in a vapor diffusion sitting drop plate (96-well plate, 100 nL reservoir buffer + 100 nL protein). The best condition was identified as 0.1 M tri-sodium citrate pH 5.5 + 20% w/v PEG3000. Crystals from this well were harvested and made into seeds with Hampton Research’s Seed Bead Kit and used in a hanging drop setup with 1 µl 10 mg/ml M208W PLpro plus 1 µl mother liquor (0.1 M trisodium citrate pH 5.5, 20% w/v PEG3000) and 0.2 µl seeds. Crystals from this drop were fished and cryoprotected with 0.1 M trisodium citrate pH 5.5, 20% w/v PEG3000, 30% glycerol before vitrification. Data were collected from the Australian Synchrotron (Australian Nuclear Science and Technology Organization, ANSTO) beamline, MX2 (Aragão et al. 2018) at a wavelength of 0.9537 Å and a temperature of 100 K. Data were indexed with using XDS (Kabsch 2010) and scaled with Aimless (Evans and Murshudov 2013) and processed with Truncate (French and Wilson 1978). The 2.00 Å structure of M208W PLpro (PDB: 8VEC) was solved using Phaser (McCoy et al. 2007) by molecular replacement with PDB 8FWN (unpublished). R-free flags were selected in Phenix (Adams et al. 2011). The refinement in PHENIX started with a round of rigid body refinement followed by simulated annealing. Subsequently, each refinement cycle consisted of XYZ (reciprocal, real-space) and individual B-factor refinement, and model building in Coot (Emsley et al. 2010). Zinc was positioned in the density center near the zinc-finger and water molecules were added in the final stages of refinement.

### B-factor comparison

We compared the B-factors of C111S (7D47) and M208W (8VEC) against 53 other PLpro structures in the PDB with resolution lower than 3.5 Å to look for regions that are stabilized by the respective mutations. Because crystallization conditions and data quality alter B-factors in ways that are not intrinsic to the protein of interest we scaled each PDB’s B-factors to either C111S or M208W data before looking for regions of altered stability when compared to the population.

Tables were made for each PDB file and contained columns for the residue position and its average B-factor calculated from C, Cα, N, O and, when not Glycine, Cβ atoms. The true intrinsic B-factors (*B_ture_resi_N_*) of PLpro were estimated from the average of all 55 PDB files, which had multiple spacegroups and crystallization conditions.

To determine regions of stabilization and destabilization in M208W (8VEC) and C111S (7D47) we scaled B-factors with the “optim” function in R (R studio v4.2.0), using the Broyden–Fletcher– Goldfarb–Shanno (BFGS) method (Fletcher 2000) to find scaling (k) and translating (b) constants.

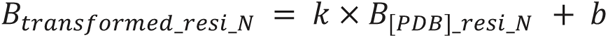

The BFGS method is an iterative process that converges on appropriate scaling values by minimizing the sum-of-squares total differences between B-factor tables.

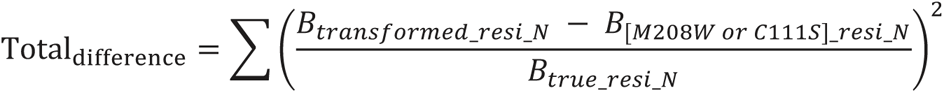

* The Zinc finger residues 186 to 197 and 219 to 232 were excluded in this procedure as they are disordered in many PLpro structures.

where *B_transformed_resi_N_* is the average residue B-factor at position N of the PDB file upon applying transformation, *B_[M208W or C111S]_resi_N_* is the average residue B-factor at position N of the PDB file being assessed, while *B_ture_resi_N_* is the average residue B-factor at position N from the average of all PDB files.

The success of this approach is dependent on the initial values of *k* and *b*. *k* was specified as the average of all residue B-factors of M208W or C111S divided by the average of all residue B-factors of the PDB structure of interest and b was initiated from zero:

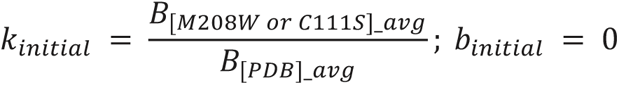

Where *B_[M208W or C111S]_avg_* is the average B-factor of M208W or C111S; *B_[PDB]_avg_* is the average B-factor of the PDB file without applying any transformation.

Once optimal *k* and *b* values were determined we plotted data from all scaled PDB B-factors to ensure scaling was successful (Fig. **5d**). We also applied the same normalization parameters to the Wilson B-factor for each structure (Supplementary table. **2**). The transformed Wilson B-factors are within 5Å^2^ of the Wilson B-factor of our M208W structure, showing that our transformation is appropriate. Residues from M208W or C111S PLpro with B-factors more than 2 standard deviations higher than the mean of all other data were reported as destabilized and those more than 2 standard deviations below were classified as stabilized residues.

### PLpro sequence alignment

The alignment of PLpros from SARS-CoV-1, HKU1, 229E, NL63, OC43, BtSARS, Bt273, Bt133, BtHKU9, MHVJ, BCoV, TGEV, aIBV was taken from (Chaudhuri et al. 2011), and we applied the same approach to include MERS and SARS-CoV-2 PLpro in the final alignment, which was used to generate a WebLogo representation (Crooks et al. 2004). The alignment has also been supplied in the supplementary materials.

### Identification of circulating variants

The data is obtained from the literature (Portelli et al. 2020). We first chose the variants on Nsp3 PLpro region (aa746 to 1060) and found the variant with the maximum observations, which is A145D with 24688 sequences registered. In our analysis we only considered variants that had observations of more than 1% of the most observed variant (i.e 247) to exclude rare variants that might be due to sequencing errors.

## Supplementary Figures

**Extended Data Fig. 1.**
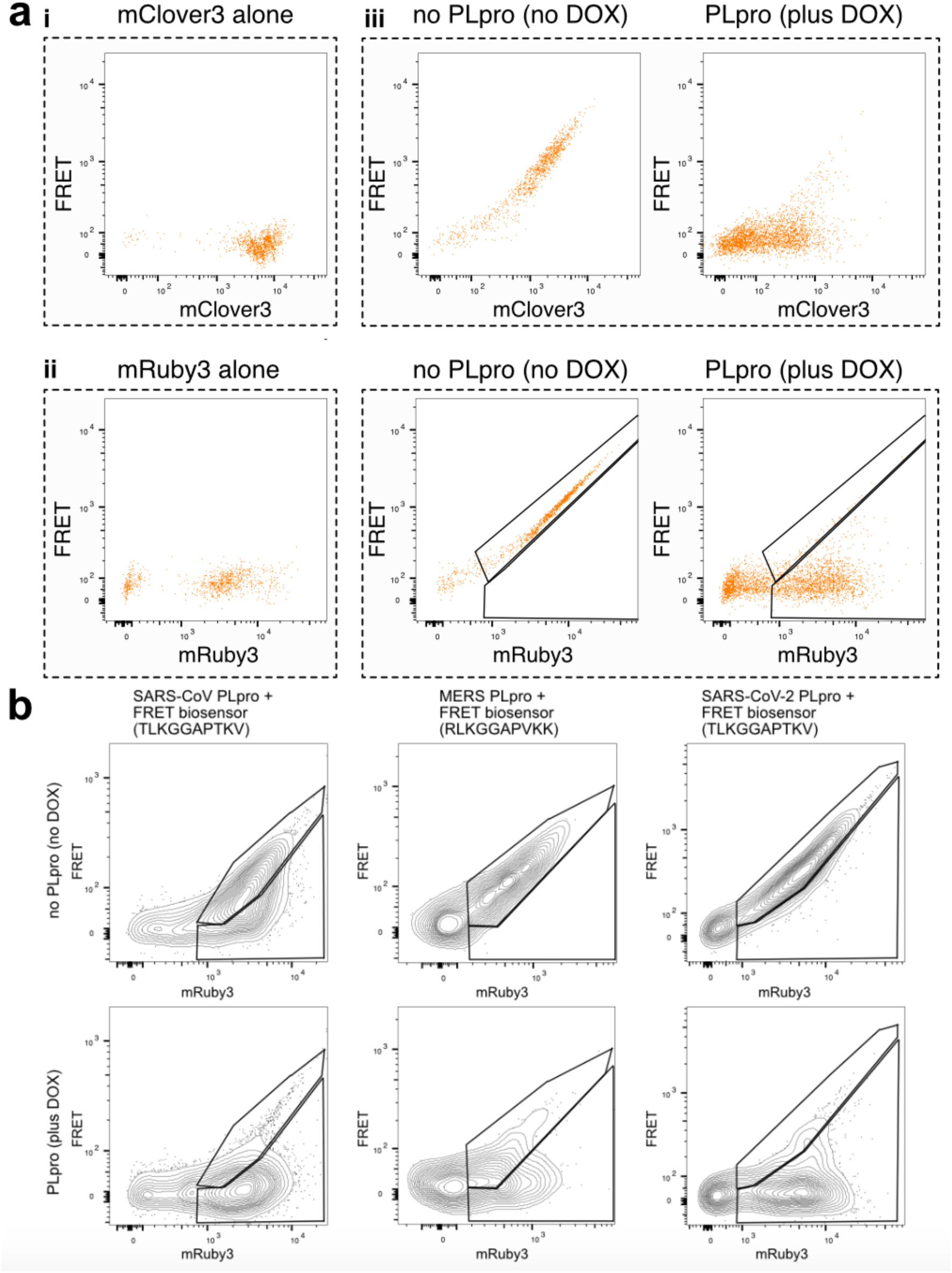
PLpro FRET assay and other coronavirus biosensors. **a)** Characterization of the SARS-CoV-2 PLpro biosensor. i) Flow cytometry plot of 293T cells expressing mClover3 (donor). ii) Flow cytometry plot of 293T cells expressing mRuby3 (acceptor). No signal is seen in the FRET channel when donor and acceptor are expressed independently. iii) 293T biosensor cells in the absence (left) and presence (right) of PLpro with FRET on the y-axis and mClover3 on the x-axis. iv) shows the same cells as iii), with mRuby3 on the x-axis. Biosensor cleavage reduced FRET and resulted in unexpected drop in mClover3 fluorescence so gating for analysis and selection was performed on mRuby3 vs FRET flow cytometry plots. **b)** Flow cytometry contour plots illustrating the sensitivity of biosensors to SARS-CoV (left), MERS (middle) and SARS-CoV-2 (right) PLpro. Upper row shows biosensor fluorescence in the absence of PLpro, lower row in the presence.

**Extended Data Fig. 2.**
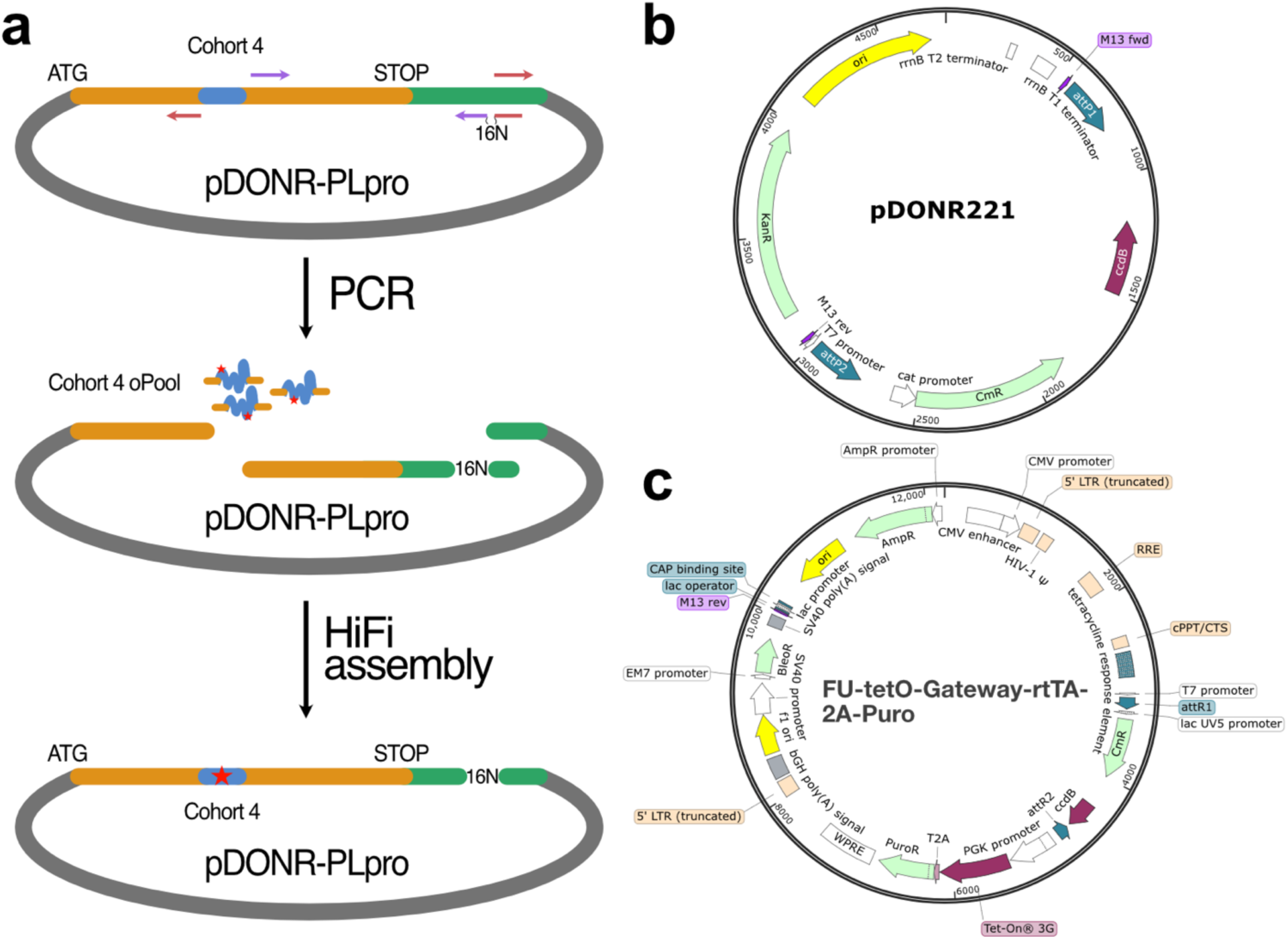
Mutagenesis strategy and vector maps. **a)** Mutagenesis strategy of PLpro DMS library, cohort 4 showing primer binding sites to generate the vector fragment and 3’ barcoded PLpro fragment with PCR. These two fragments were mixed with cohort 4 IDT oPools containing degenerate codons and assembled with NEB HiFi assembly. **b)** Vector map of pDONR221 **c)** Vector map of FU-tetO-Gateway-rtTA-2A-Puro.

**Extended Data Fig. 3.**
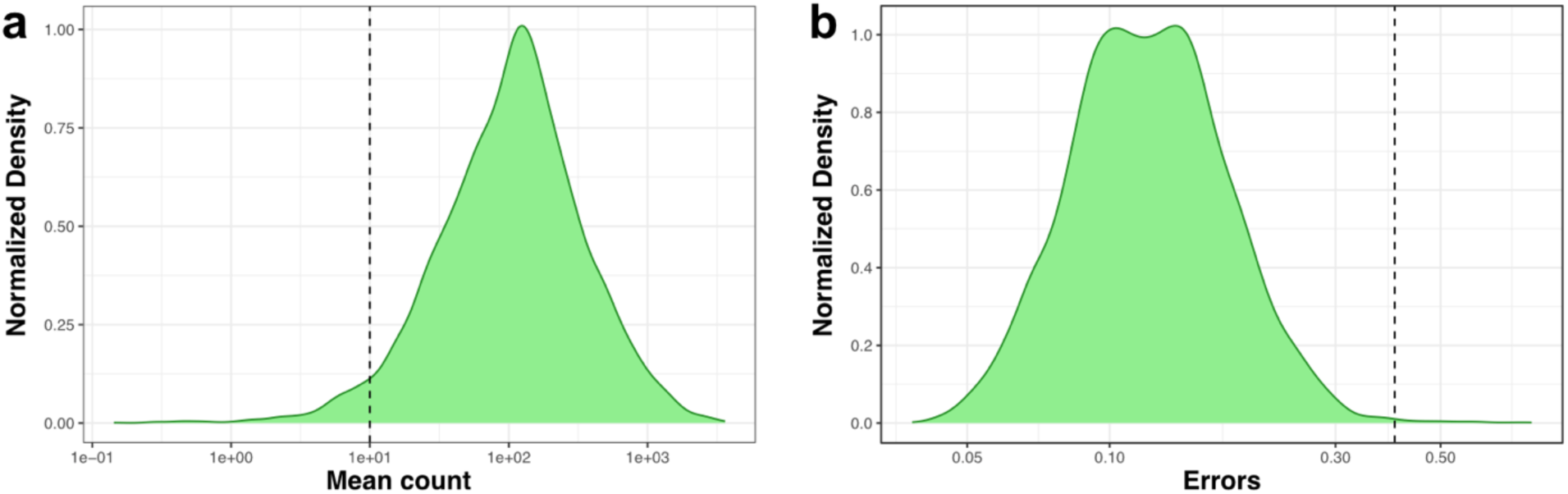
Activity Data filters. **a)** The average reads per variant in the activity dataset depicted as a density distribution plot. Reads with less than 10 mean counts were filtered. **b)** The distribution of errors for variants in the activity dataset. Variants with errors greater than 0.4 were filtered.

**Extended Data Fig. 4.**
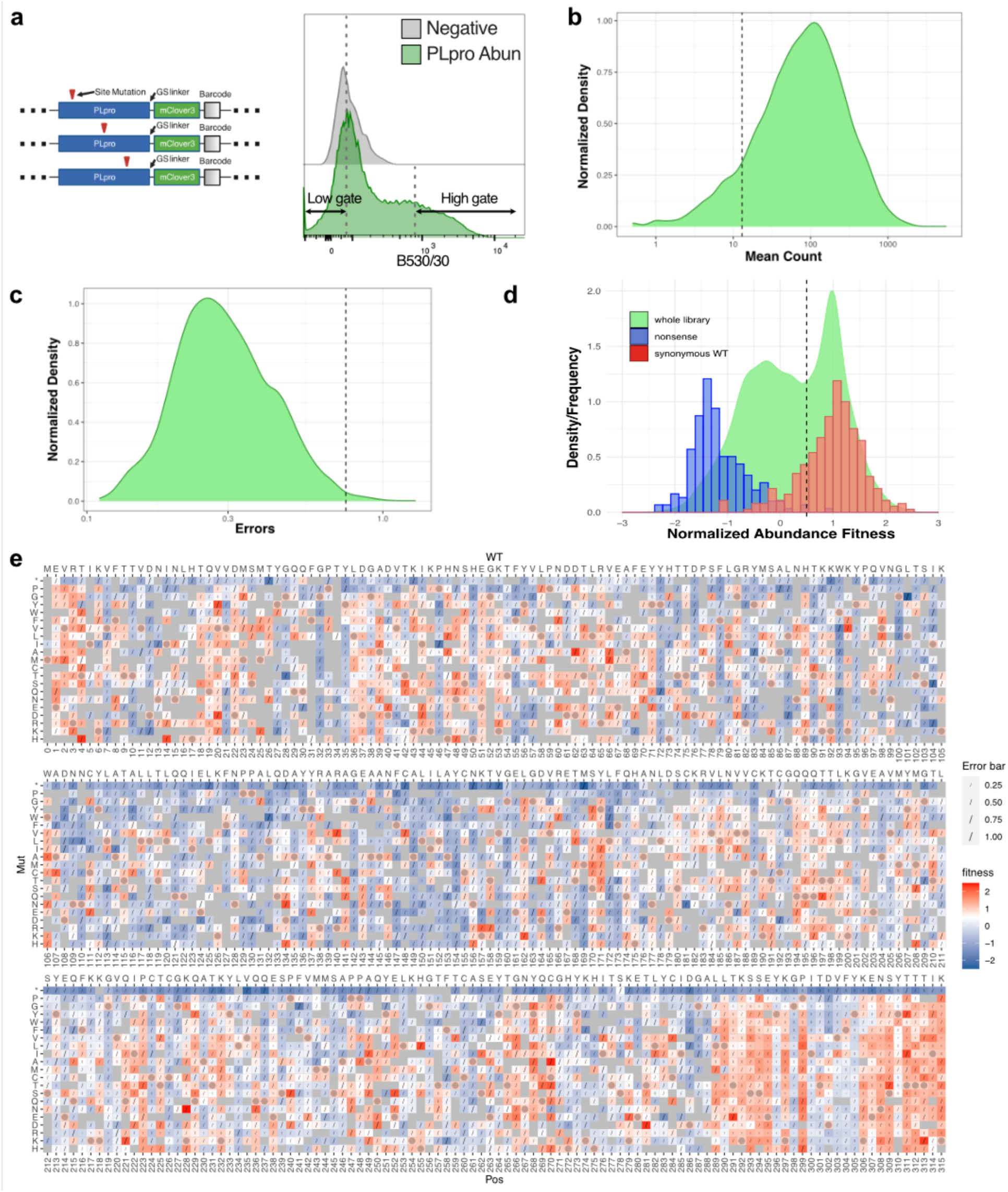
Abundance assay design and Sequence-Function map. **a)** Cartoon of the PLpro-mClover3 fusion library (*left*) and flow cytometry plot (*right*) of mClover3 fluorescence in 293T cells before (grey) and after (green) introduction of PLpro abundance library. Cells were sorted according to the high and low gates marked. **b)** The average reads per variant in the abundance dataset depicted as a density distribution plot. Reads with less than 13 mean counts were filtered. **c)** The distribution of errors for variants in the abundance dataset. Variants with errors greater than 0.75 were filtered. **d)** The distribution of normalized abundance dataset fitness scores for the whole library (density; green), overlaid with the frequency of scores from synonymous wildtype variants (red; set at 1) and nonsense variants at positions 1-305 (blue; set at −1). **e)** The normalized sequence-function map of abundance data from 2 independent transductions. The bottom x-axis indicates the position; the top x-axis indicates WT sequence; and the y-axis indicates the mutation. Fitness is shown in a two-color gradient with red indicating abundant variants, blue low abundant variants, and grey missing variants. The white point of the gradient was set at 0.5 to discriminate distinct populations seen is d). Wildtype variants are marked with a circle. Errors are indicated with a diagonal line or if above 1, an X.

**Extended Data Fig. 5.**
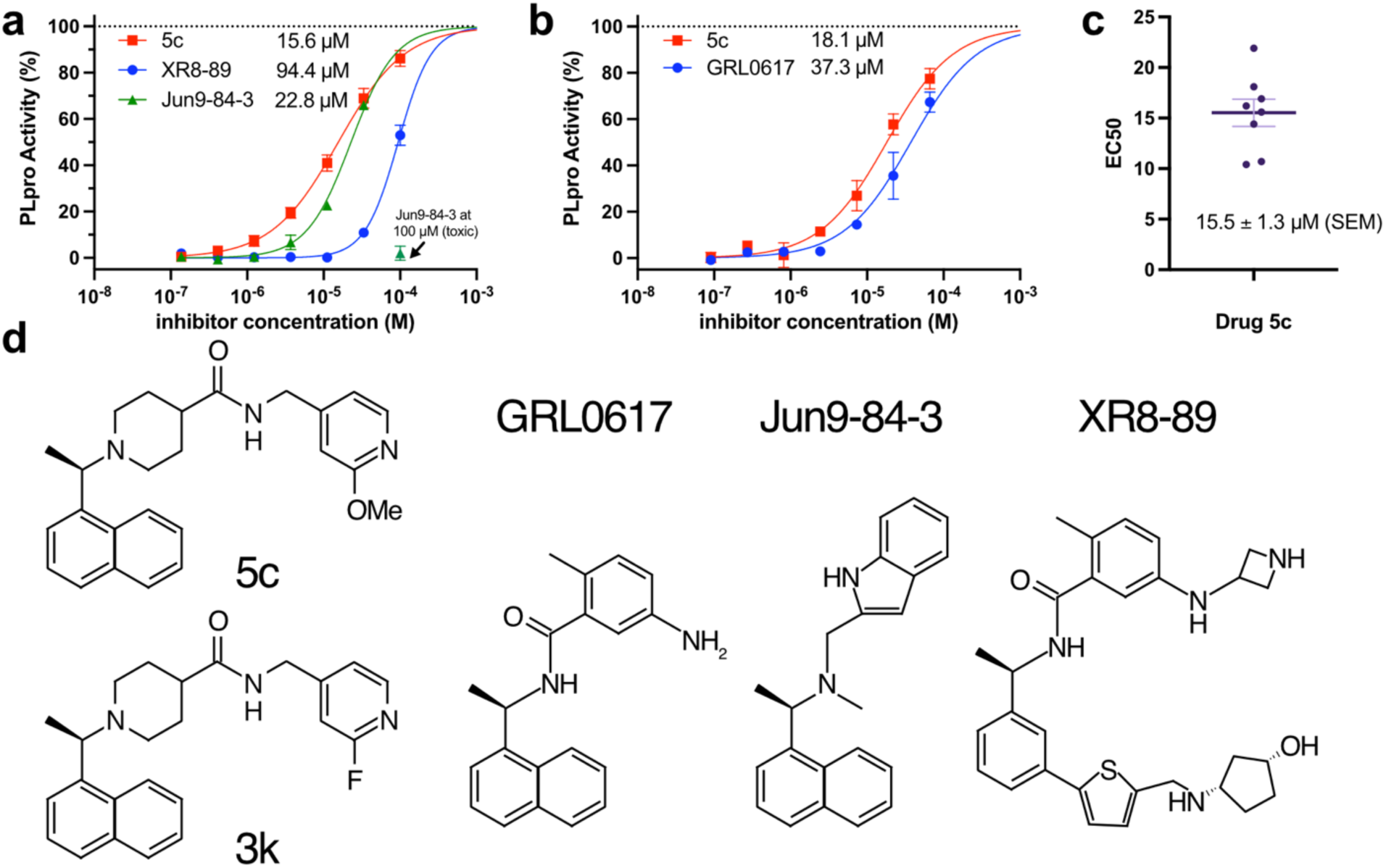
Dose response of lead PLpro compounds in the cellular FRET assay. **a)** Dose response of 5c, Jun9-84-3 and XR8-89 (technical triplicates). Error bars: mean ± SD **b)** Dose response of 5c and GRL0617 (technical triplicates). Error bar: mean ± SD; Data was normalized so the top was 100 and bottom was 0 after fitting the 5c dose response curve that had a hillslope of 1. XR8-89 and Jun9-84-3 had hillslopes of 2 and 1.5 respectively. EC50 values for each experiment are marked, N = 1. **c)** Plot C shows the EC50s of eight independent measurements of 5c, which had a mean of 15.5 µM, and standard error of the mean (SEM) of 1.3 µM. **d)** Chemical structures of compounds referred to in this study.

**Extended Data Fig. 6.**
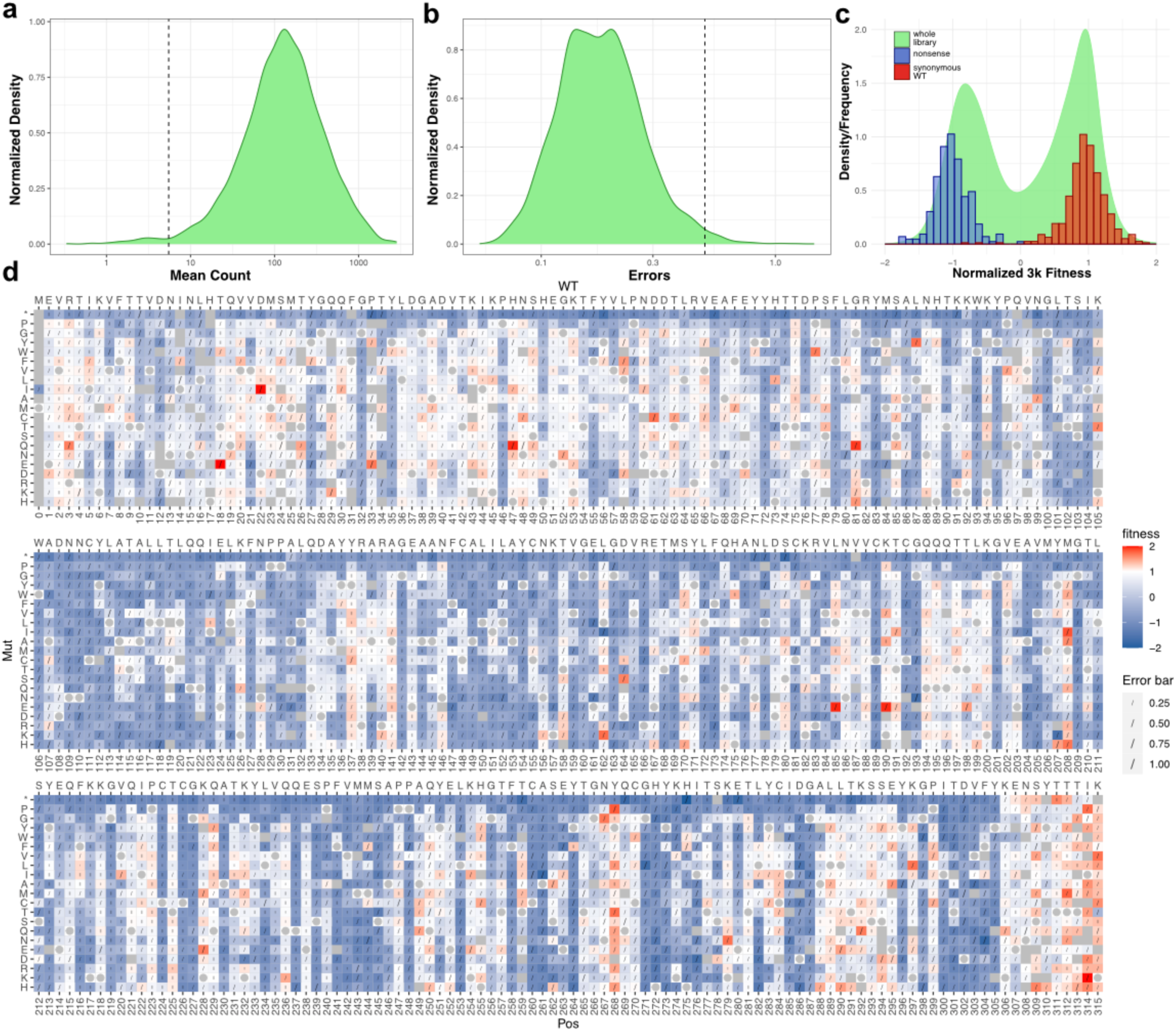
Sequence-function map of normalized 3k drug-escape fitness scores. **a)** The average reads per variant in the 3k drug-escape dataset depicted as a density distribution plot. Reads with less than 5.5 mean counts were filtered. **b)** The distribution of errors for variants in the 3k drug-escape dataset. Variants with errors greater than 0.5 were filtered. **c)** The distribution of normalized 3k drug-escape fitness scores for the whole library (density; green), overlaid with the frequency of scores from synonymous wildtype variants (red; set at 1) and nonsense variants at positions 1-305 (blue; set at −1). **d)** The normalized sequence-function map of the data from 3 independent transductions. The bottom x-axis indicates the position; the top x-axis indicates WT sequence; and the y-axis indicates the mutation. Fitness is shown in a three-color gradient with red indicating escape variants, blue non-escape variants, and the white point set at the wild-type score of 1. Grey indicates missing variants. Wildtype variants are marked with a circle. Errors are indicated with a line.

**Extended Data Fig. 7.**
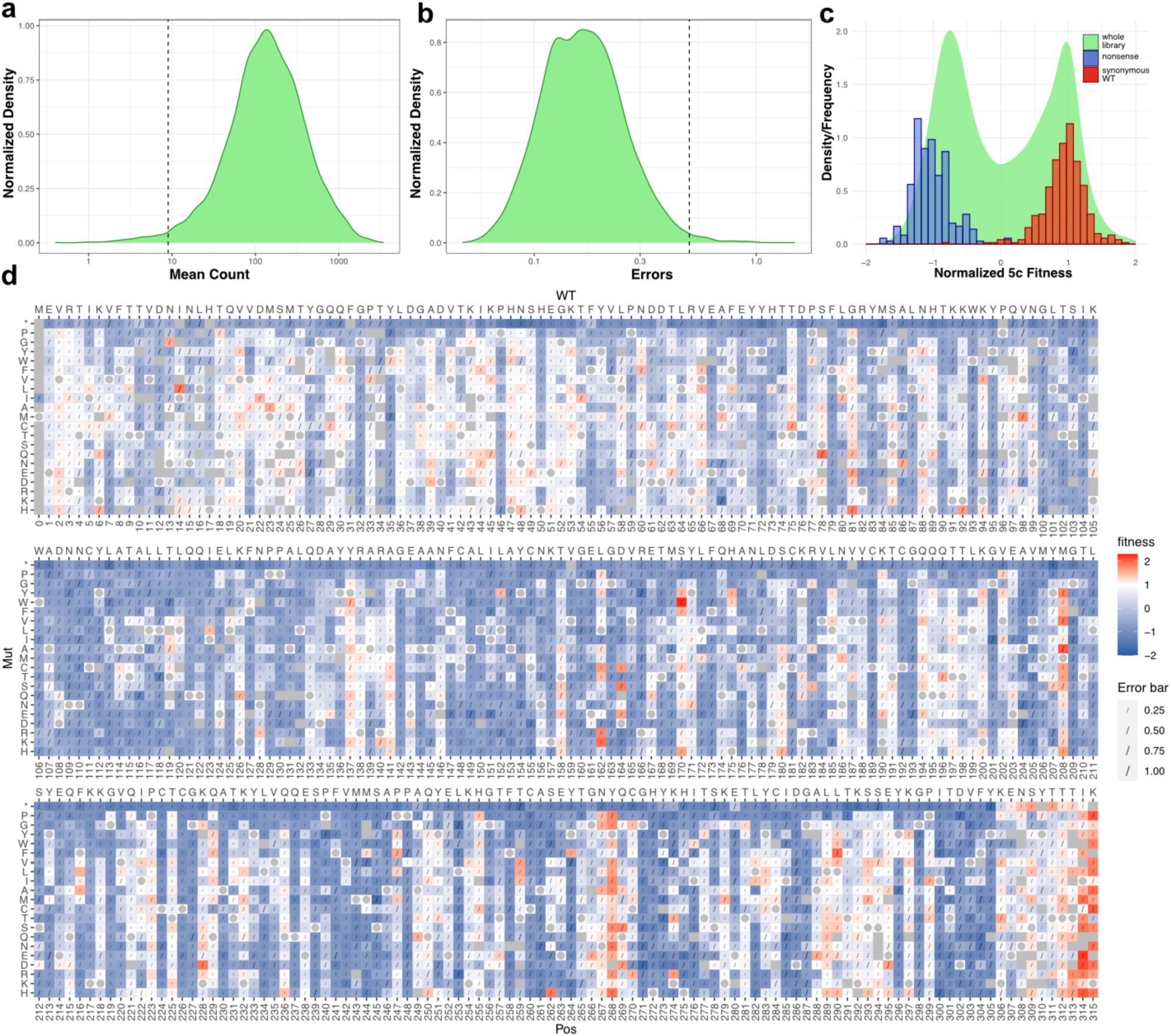
Sequence-function map of normalized 5c drug-escape fitness scores. **a)** The average reads per variant in the 5c drug-escape dataset depicted as a density distribution plot. Reads with less than 9 mean counts were filtered. **b)** The distribution of errors for variants in the 5c drug-escape dataset. Variants with errors greater than 0.5 were filtered. **c)** The distribution of normalized 5c drug-escape fitness scores for the whole library (density; green), overlaid with the frequency of scores from synonymous wildtype variants (red; set at 1) and nonsense variants at positions 1-305 (blue; set at −1). **d)** The normalized sequence-function map of the data from 5 independent transductions. The bottom x-axis indicates the position; the top x-axis indicates WT sequence; and the y-axis indicates the mutation. Fitness is shown in a three-color gradient with red indicating escape variants, blue non-escape variants, and the white point set at the wild-type score of 1. Grey indicates missing variants. Wildtype variants are marked with a circle. Errors (sigma) are indicated with a line.

**Extended Data Fig. 8.**
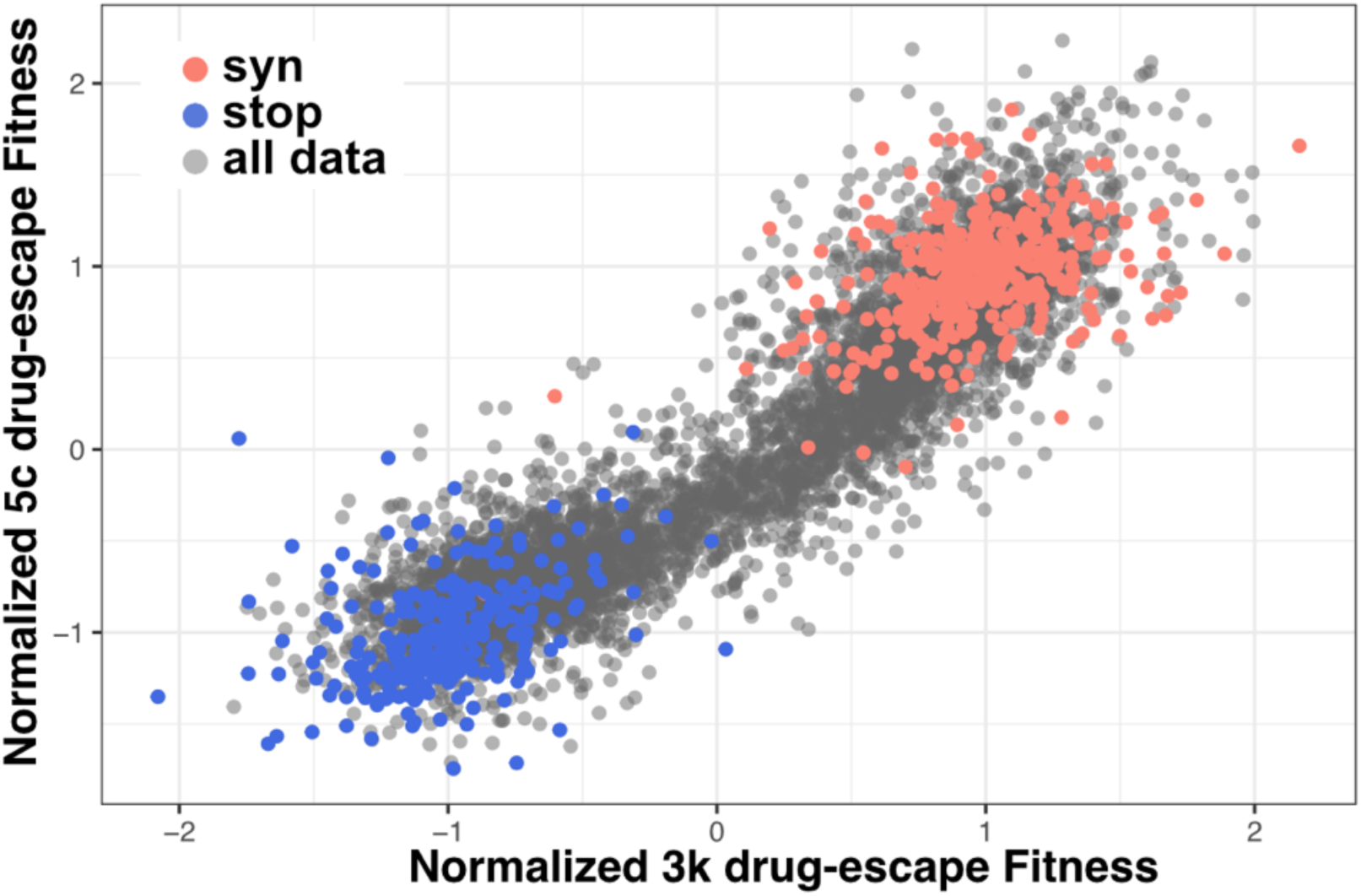
Correlation between normalized drug-escape scores between 3k and 5c and structures of both compounds. **a)** Plot of Normalized 3k (x-axis) and 5c (y-axis) drug-escape fitness scores. Nonsense variants (positions 1-305) are shown in blue, synonymous WT are shown in red and the rest shown in grey.

**Extended Data Fig. 9.**
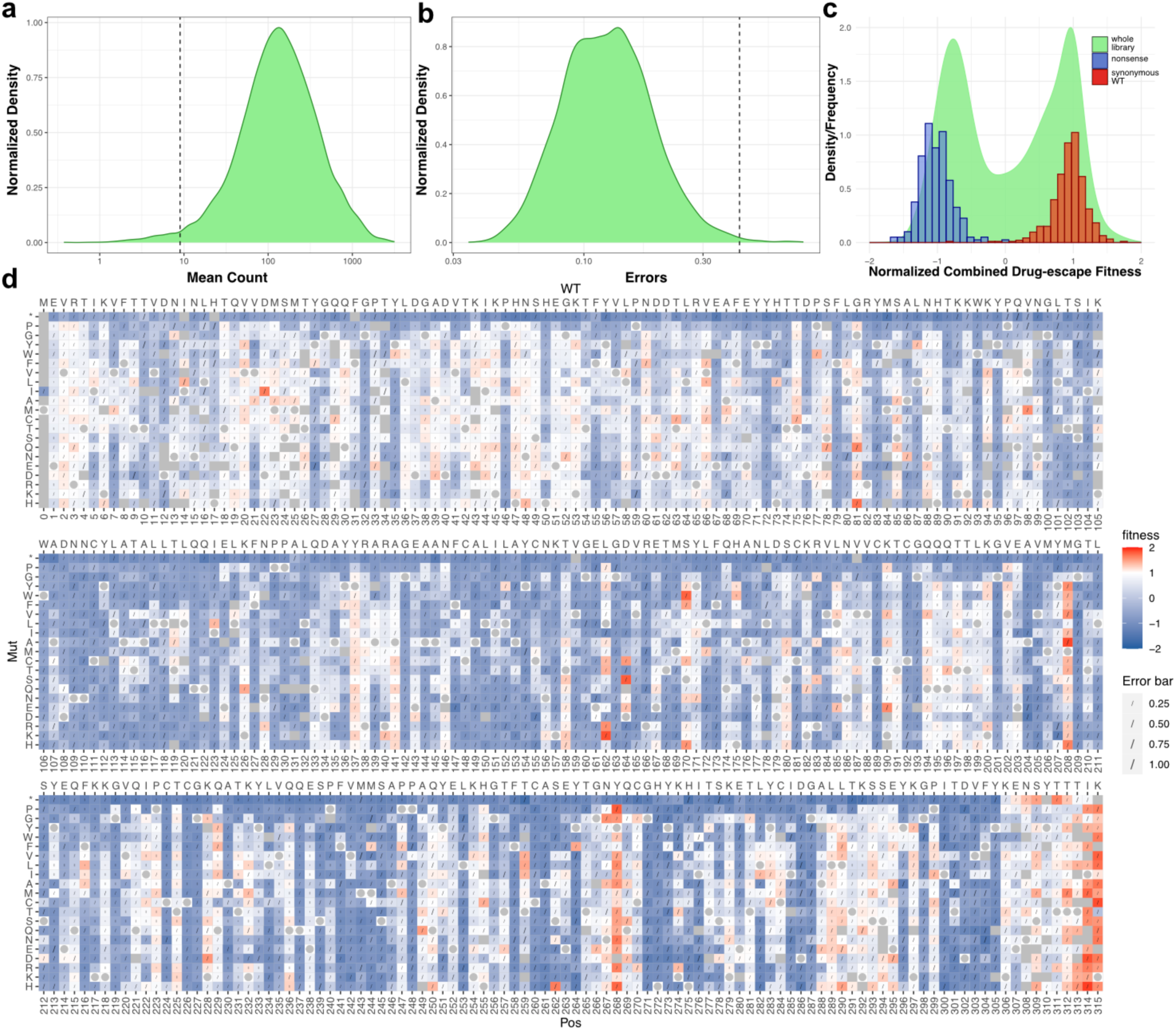
Sequence-function map of normalized 3k & 5c combined drug-escape fitness scores. **a)** The average reads per variant in the combined drug-escape dataset depicted as a density distribution plot. Reads with less than 9 mean counts were filtered. **b)** The distribution of errors for variants in the combined drug-escape dataset. Variants with errors greater than 0.42 were filtered. **c)** The distribution of normalized combined drug-escape fitness scores for the whole library (density; green), overlaid with the frequency of scores from synonymous wildtype variants (red; set at 1) and nonsense variants at positions 1-305 (blue; set at −1). **d)** The normalized sequence-function map of the data. The bottom x-axis indicates the position; the top x-axis indicates WT sequence; and the y-axis indicates the mutation. Fitness is shown in a three-color gradient with red indicating escape variants, blue non-escape variants, and the white point set at the wild-type score of 1. Grey indicates missing variants. Wildtype variants are marked with a circle. Errors (sigma) are indicated with a line.

**Extended Data Fig. 10.**
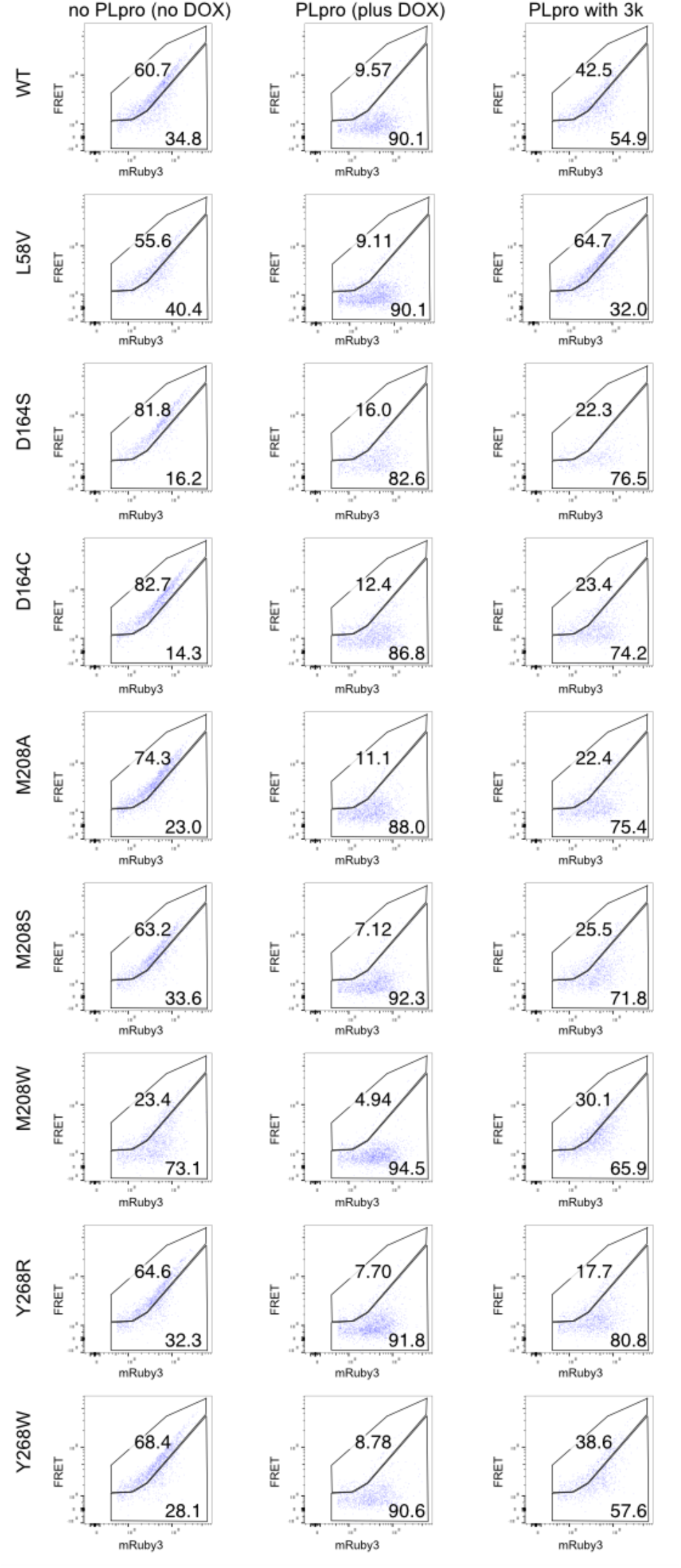
Flow cytometry plots of PLpro variants. Each row represents the PLpro variant indicated; Conditions are shown by columns. The first column contains cells with no dox treatment; the second column contains cells treated with DOX; the third column contains cells treated with DOX and 3k. The x-axis is mRuby3 (YG610/20) and y-axis is FRET (B610/20). In each plot, the upper gate shows FRET positive cells (cells with no PLpro activity) and the lower gate FRET negative (cells with PLpro activity). Percentages of cells in each gate are shown. One independent transduction was tested twice with similar results.

**Extended Data Fig. 11.**
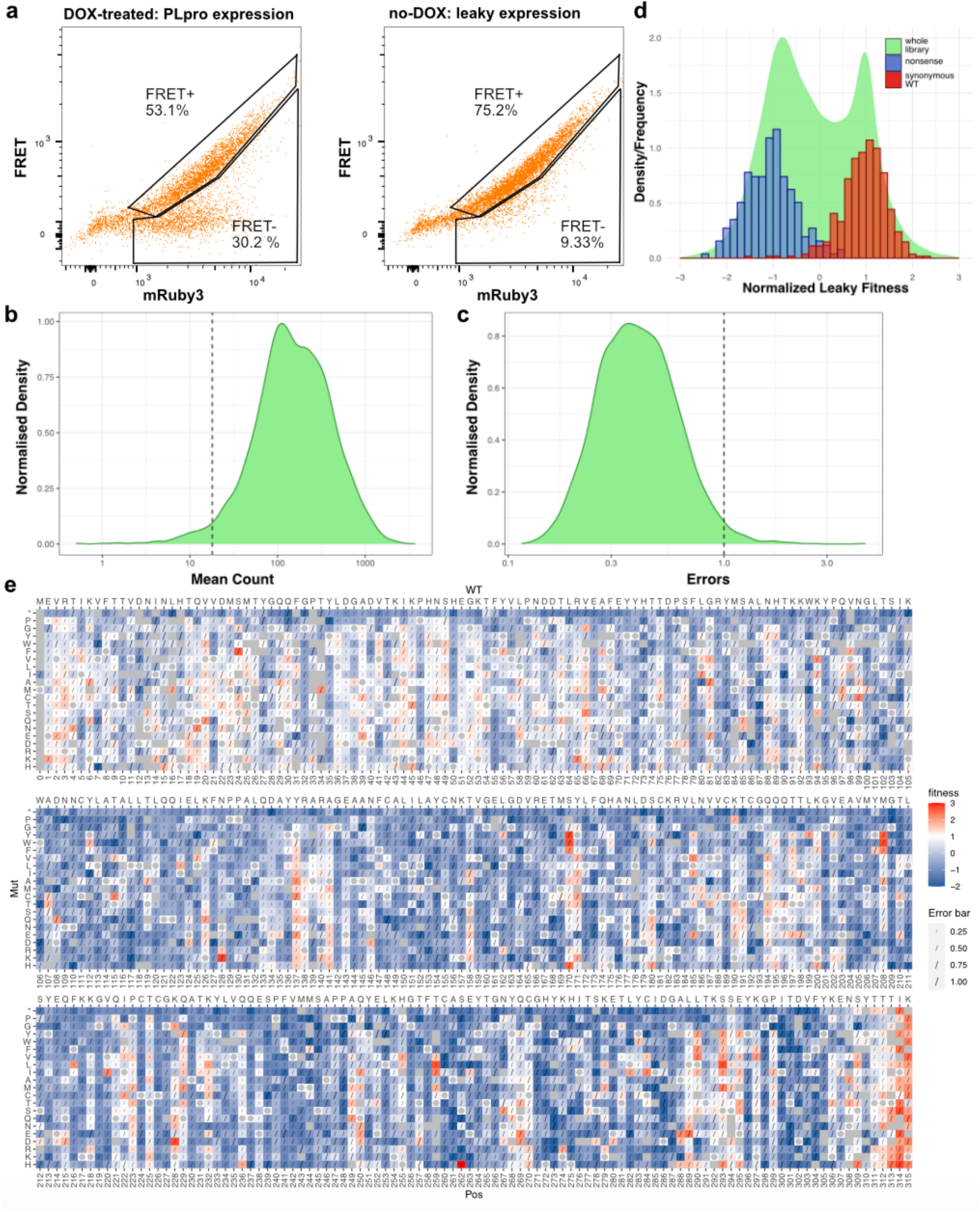
Normalized PLpro leaky expression sequence-function map. **a)** Flow cytometry plots of the PLpro activity library in the presence of dox (left) and in the absence (right). The x-axis is mRuby3 (YG610/20) and y-axis is FRET (B610/20). **b)** The average reads per variant in the leaky expression dataset depicted as a density distribution plot. Reads with less than 18 mean counts were filtered. **c)** The distribution of errors for variants in the leaky expression dataset. Variants with errors greater than 1 were filtered. **d)** The distribution of normalized leaky expression fitness scores for the whole library (density; green), overlaid with the frequency of scores from synonymous wildtype variants (red; set at 1) and nonsense variants at positions 1-305 (blue; set at −1). **e)** The normalized sequence-function map of the data from 2 independent transductions. The bottom x-axis indicates the position; the top x-axis indicates WT sequence; and the y-axis indicates the mutation. Fitness is shown in a three-color gradient with red indicating active and leaky variants, blue not leaky or inactive variants, and the white point set at the wild-type score of 1. Grey indicates missing variants. Wildtype variants are marked with a circle. Errors (sigma) are indicated with a line.

**Extended Data Fig. 12.**
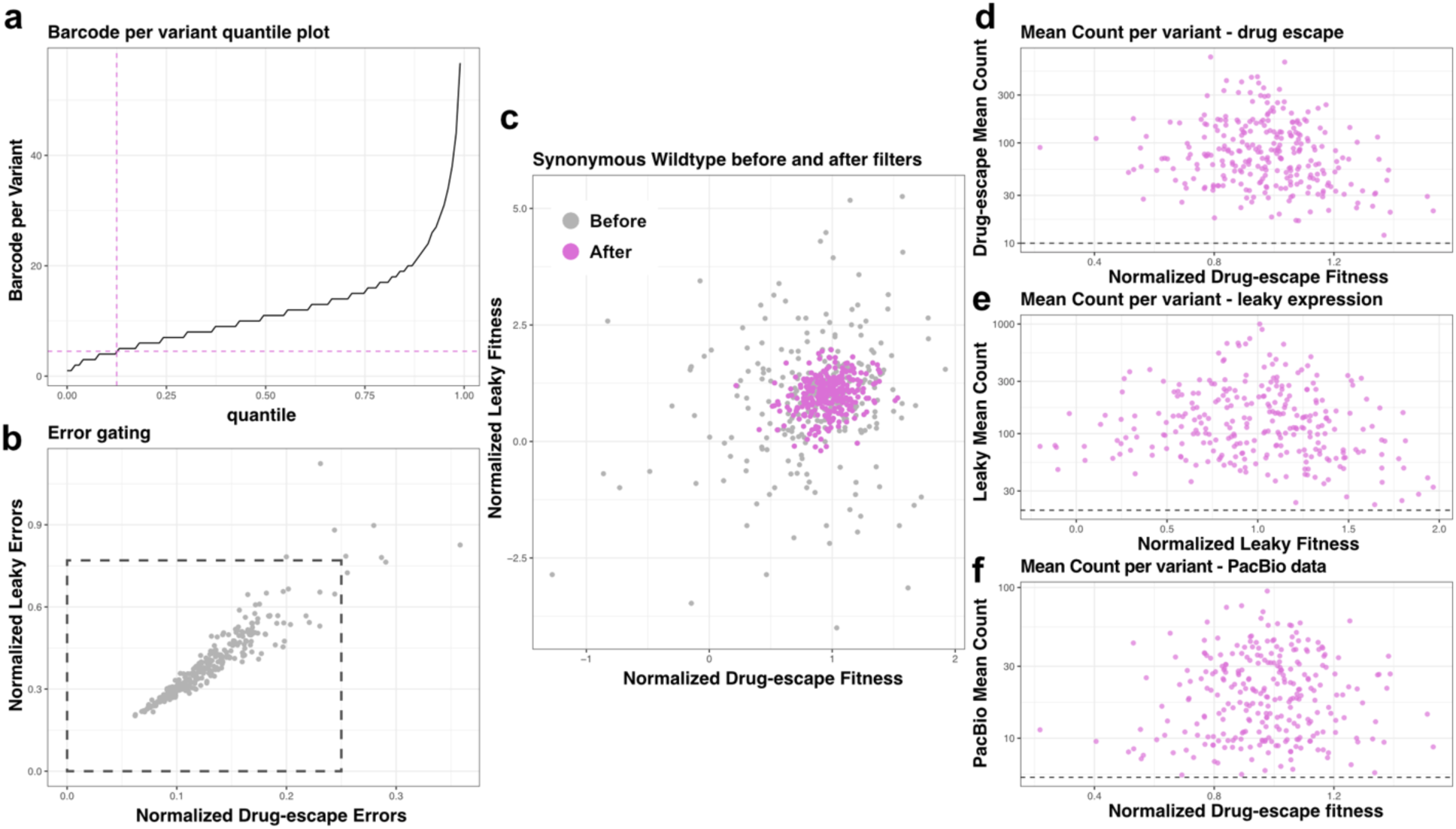
Noise reduction to allow comparison between combined drug-escape and leaky expression fitness scores guided by synonymous wildtype fitness scores. **a)** Barcodes per variant quantile plot. Based on this plot, variants with less than 5 barcodes were filtered removing ∼12% of the data. **b)** Plot of errors seen in drug-escape and leaky-expression fitness scores. Variants with high errors in both datasets were removed. The dashed rectangle indicates data that passed selection. **c)** The distribution of synonymous wildtype drug-escape and leaky-expression fitness scores after filters from (a) and (b) were applied. Data before filtering is shown in grey and after filtering is shown in magenta. **d-e)** The synonymous wildtype variants were assessed for fitness score versus mean count. **d)** A minimum requirement of 10 read counts per variant was set for the drug-escape dataset. **e)** A minimum requirement of 20 read counts per variant was set for the drug-escape dataset. **f)** The average number of reads per variant from PacBio data was determined from the distribution of synonymous wildtype variant drug-escape fitness scores against the mean read count in PacBio sequencing data. Variants with less than 5.5 reads were discarded.

**Extended Data Fig. 13.**
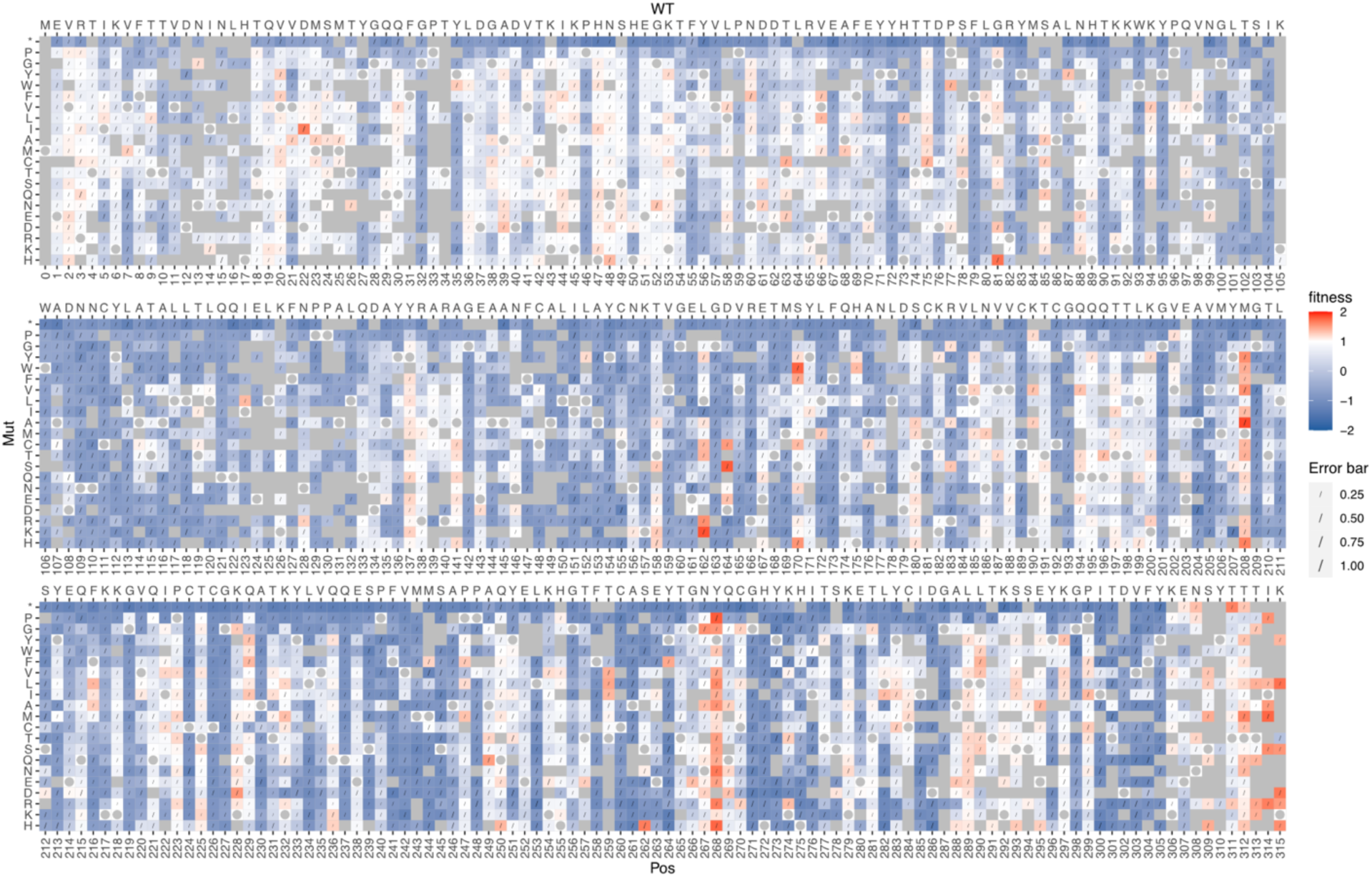
Sequence-function map of normalized combined drug-escape fitness scores after applying the filters outlined in Extended Data Fig. 12. The bottom x-axis indicates the position; the top x-axis indicates WT sequence; and the y-axis indicates the mutation. Fitness is shown in a three-color gradient with red indicating escape variants, blue non-escape variants, and the white point set at the wild-type score of 1. Grey indicates missing variants. Wildtype variants are marked with a circle. Errors (sigma) are indicated with a line.

**Extended Data Fig. 14.**
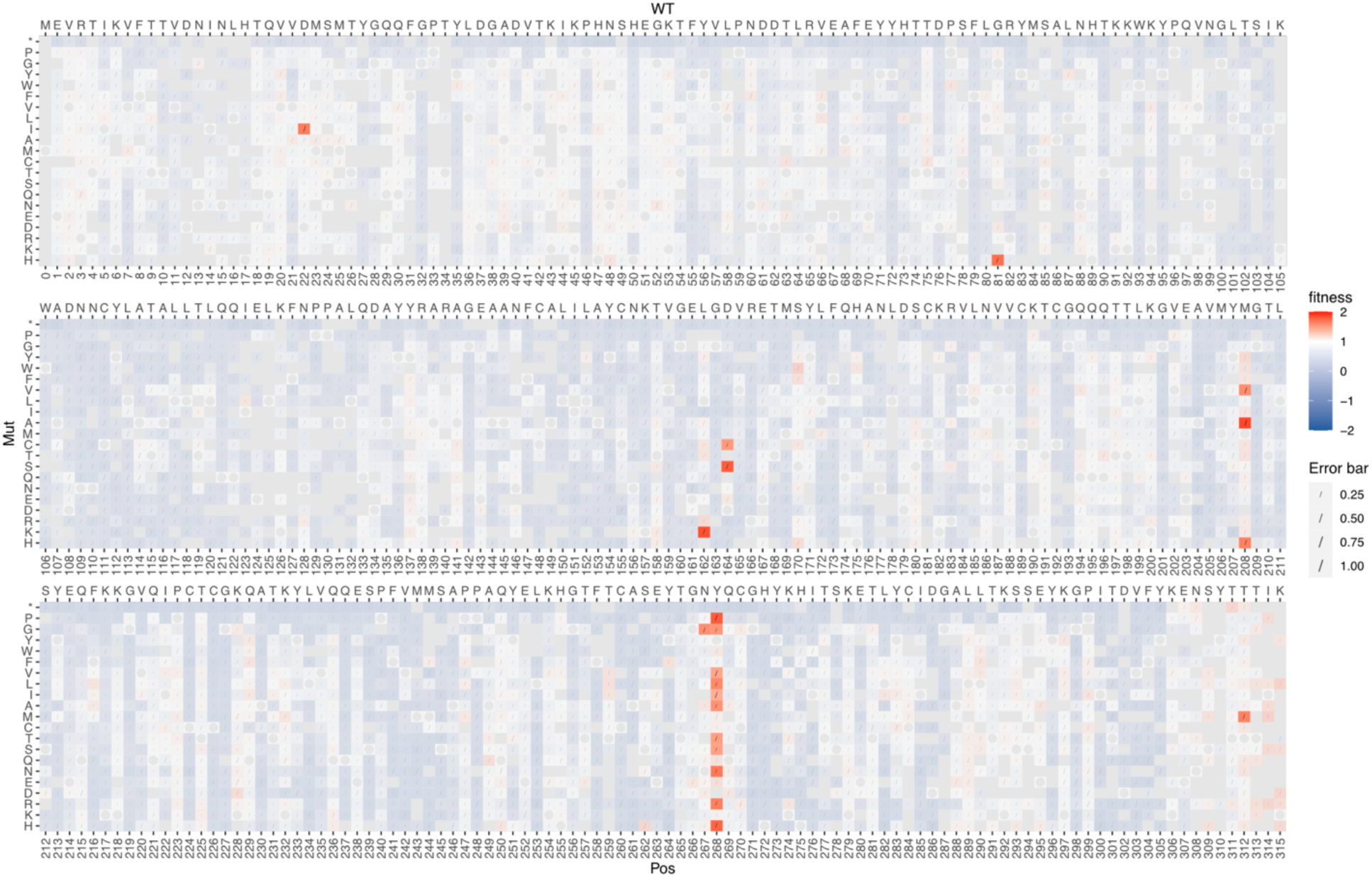
Drug-escape variants falling in the gate defined in Fig. 4c. The sequence function map in *Extended Data Fig. 13* was masked with a semi-transparent white layer except for variants identified as candidate escape variants, shown in shades of red.

**Extended Data Fig. 15.**
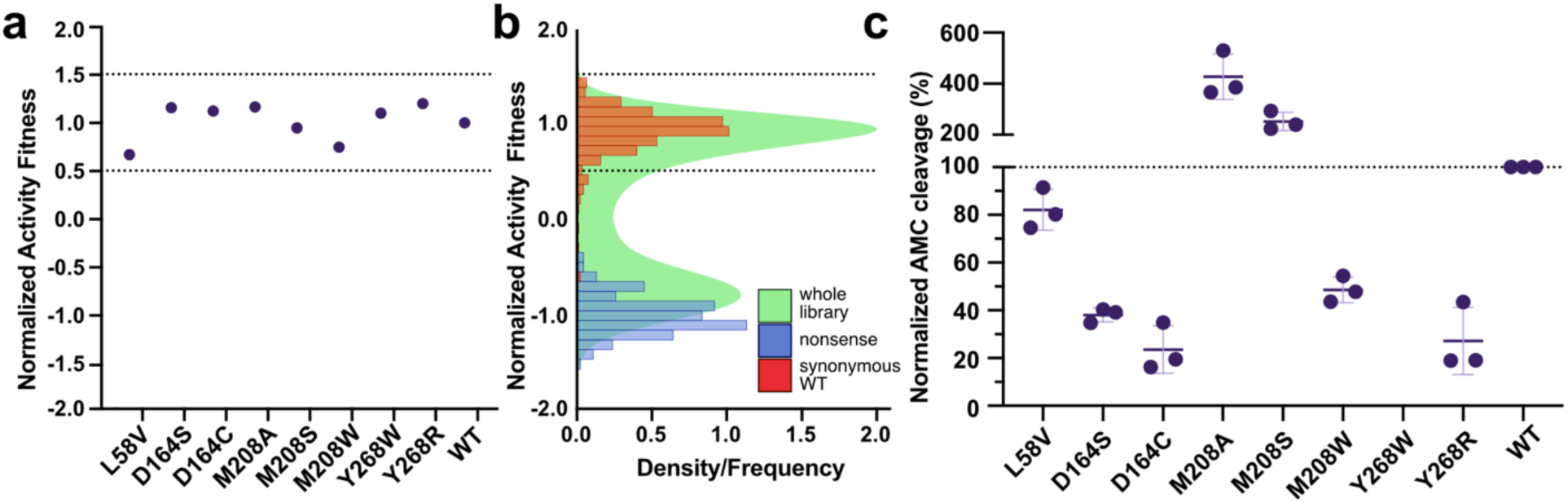
Comparison of DMS activity fitness scores (at steady state) with activity of recombinant proteins (initial rates). **a**) Normalized activity fitness scores extracted from the DMS activity dataset. Dashed lines indicate wildtype-like activity determined from the graph in (b). **b)** The distribution of normalized PLpro activity fitness scores for the whole library (density; green), overlaid with the frequency of scores from synonymous wildtype variants (red; set at 1) and nonsense variants at positions 1-305 (blue; set at −1). **c)** Z-RLRGG-AMC cleavage by recombinant PLpro variants after 2 h incubation at room temperature. All data is normalized to wildtype (100%). Data is from three independent experiments. Error bars indicates mean ± SD.

**Extended Data Fig. 16.**
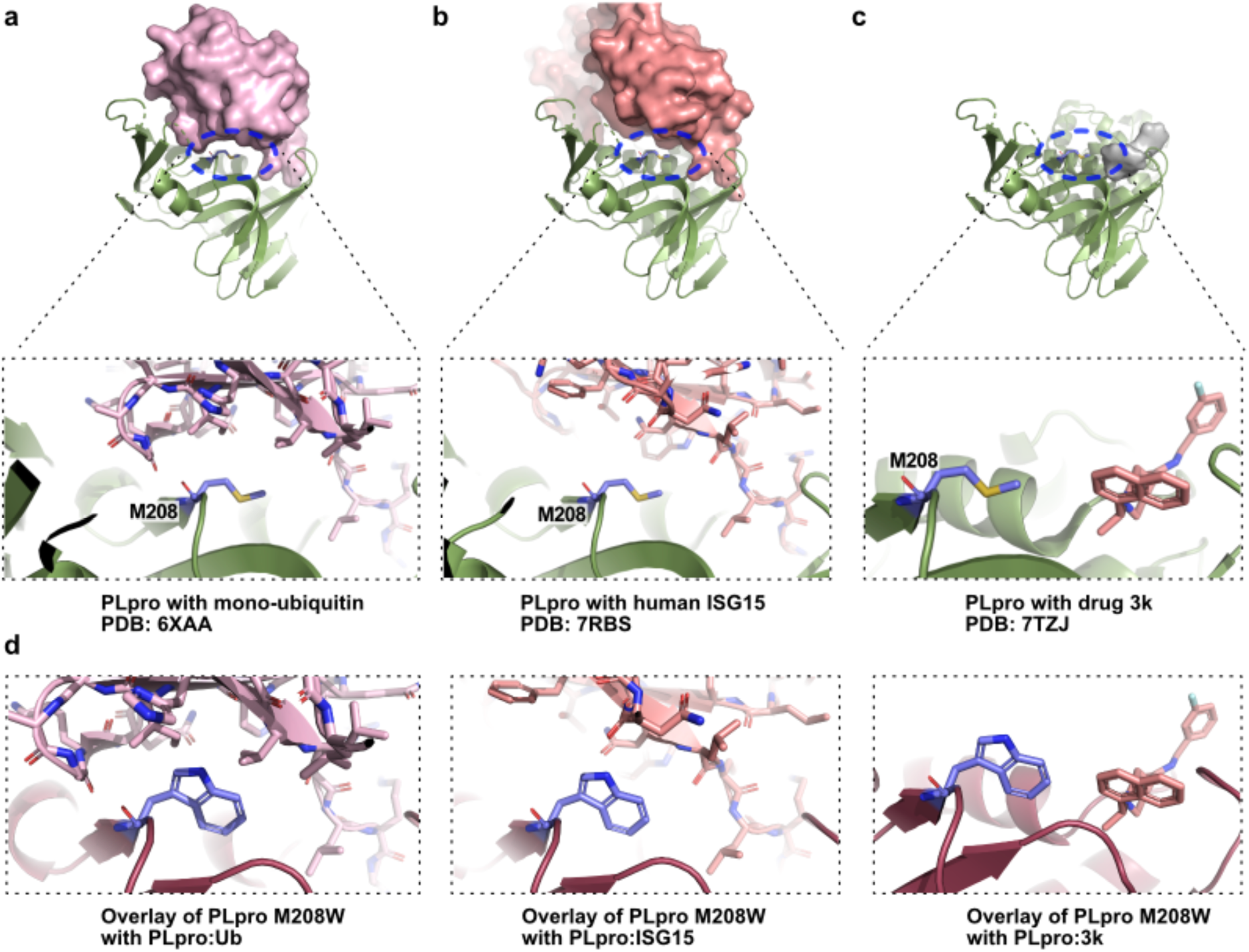
Met208 does not directly contact substrate or inhibitor. **a)** PLpro (green) with Ub (pink) (PDB: 6XAA) (Klemm et al. 2020) **b)** PLpro (green) with human ISG15 (red) (PDB: 7RBS) (Wydorski et al. 2023) **c)** PLpro (green) with 3k (grey) (PDB: 7TZJ) (Calleja et al. 2022). The location of Met208 is highlighted in the blue circle. Insets zoom in on Met208 and surrounding residues on substrates (in stick). **d)** Overlay of PLpro M208W with PLpro:Ub, PLpro:ISG15 and PLpro:3k.

**Extended Data Fig. 17.**
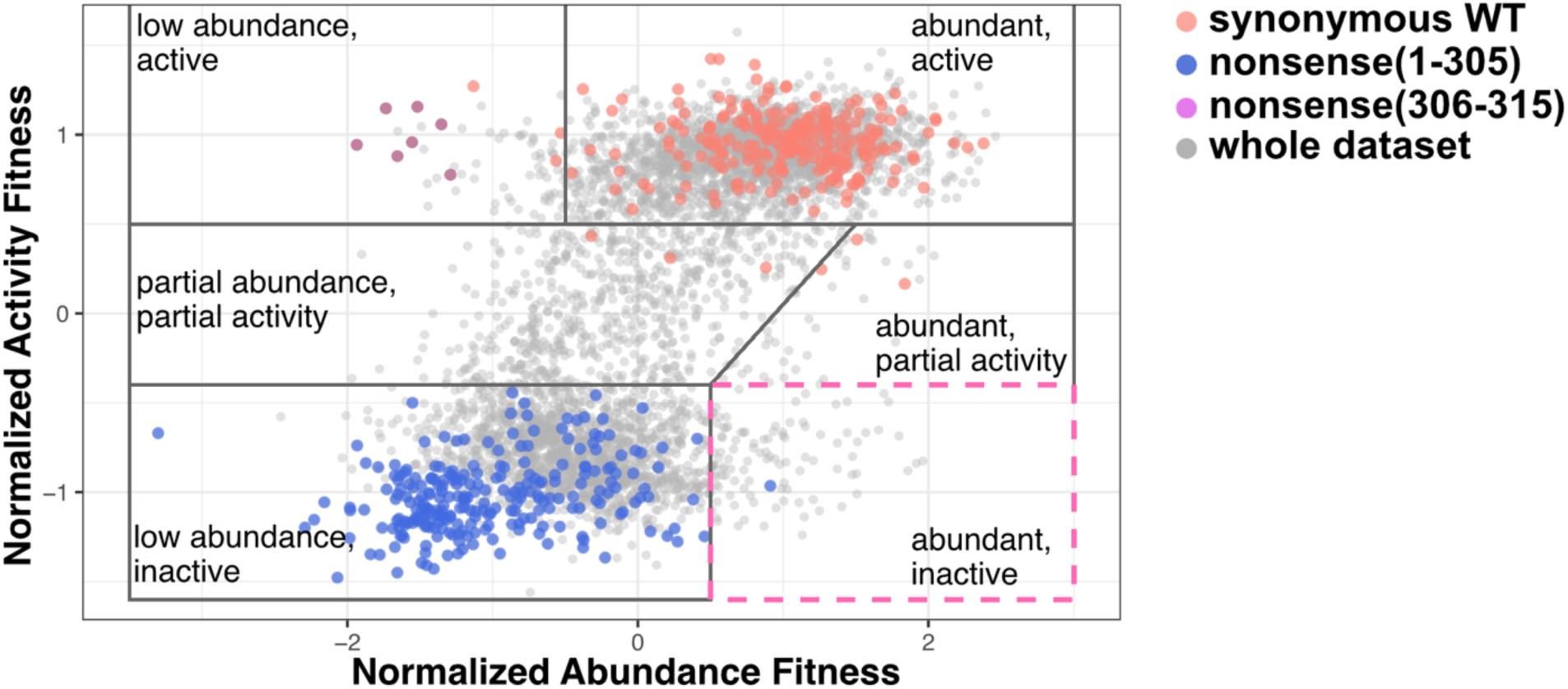
Gating strategy for classification of variants in Table 1. The scatterplot shows the correlation between normalized abundance fitness scores (x-axis) and normalized activity fitness scores (y-axis). The library is depicted in grey, synonymous wildtype are shown in red and nonsense variants (positions 1-305) are shown in blue. Nonsense variants (positions 306-315), which are active, but do not make a GFP fusion protein are labelled in magenta. The gates are chosen based on the distribution of synonymous wildtype and nonsense variant datapoints.

**Supplementary Table 1.** Data collection and refinement statistics. Statistics for the highest-resolution shell are shown in parentheses.

|  |  |
| --- | --- |
|  | M208W (PDB: 8VEC) |
| **Wavelength** | 0.9537 Å |
| **Resolution range** | 47.5 - 2.002 (2.073 - 2.002) |
| **Space group** | P 21 21 2 |
| **Unit cell** | 57.958 85.47 82.88 90 90 90 |
| **Total reflections** | 210621 (20864) |
| **Unique reflections** | 28423 (2601) |
| **Multiplicity** | 7.4 (7.4) |
| **Completeness (%)** | 99.19 (92.63) |
| **Mean I/sigma(I)** | 13.08 (1.52) |
| **Wilson B-factor** | 32.92 |
| **R-merge** | 0.1138 (1.305) |
| **R-meas** | 0.1225 (1.406) |
| **R-pim** | 0.0447 (0.516) |
| **CC1/2** | 0.999 (0.495) |
| **CC\*** | 1 (0.814) |
| **Reflections used in refinement** | 28209 (2602) |
| **Reflections used for R-free** | 2003 (180) |
| **R-work** | 0.1738 (0.2616) |
| **R-free** | 0.2170 (0.2932) |
| **CC(work)** | 0.968 (0.750) |
| **CC(free)** | 0.945 (0.718) |
| **Number of non-hydrogen atoms** | 2750 |
| **macromolecules** | 2530 |
| **ligands** | 1 |
| **solvent** | 219 |
| **Protein residues** | 318 |
| **RMS(bonds)** | 0.008 |
| **RMS(angles)** | 0.86 |
| **Ramachandran favored (%)** | 97.15 |
| **Ramachandran allowed (%)** | 2.85 |
| **Ramachandran outliers (%)** | 0.00 |
| **Rotamer outliers (%)** | 0.36 |
| **Clashscore** | 3.40 |
| **Average B-factor** | 36.74 |
| **macromolecules** | 36.24 |
| **ligands** | 75.70 |
| **solvent** | 42.34 |

**Supplementary Table 2.** PLpro structures used for B-factor analysis. Columns are as indicated. The last column contains the transformed Wilson B-factor after applying the transformation factors in columns three and four.

| **PDB** | **Resolution (Å)** | **scaling factor (k)** | **translating factor (b)** | **Wilson b factor** | **transformed Wilson b factor** |
| --- | --- | --- | --- | --- | --- |
| 8fwn | 1.5 | 0.90402689 | 12.5245887 | 26.06 | 36.0835295 |
| 6wrh | 1.6 | 0.8172159 | 10.3372938 | 28.53 | 33.6524634 |
| 7d6h | 1.6 | 0.72627769 | 12.9846059 | 30.39 | 35.0561849 |
| 7qcg | 1.75 | 0.28777225 | 20.2202365 | 45.3 | 33.2563194 |
| 7qci | 1.76 | 0.24398085 | 20.9486587 | 49.39 | 32.9988727 |
| 7qcm | 1.77 | 0.27977192 | 20.200233 | 47.23 | 33.4138608 |
| 6wzu | 1.79 | 0.7306924 | 5.84982769 | 36.07 | 32.2059025 |
| 7sgu | 1.79 | 0.48337376 | 9.81652723 | 45.22 | 31.6746886 |
| 7qcj | 1.84 | 0.28343885 | 19.9374688 | 47.37 | 33.363967 |
| 7qch | 1.88 | 0.20223123 | 20.5359937 | 60.97 | 32.8660317 |
| 7rbr | 1.88 | 0.40421242 | 14.7488056 | 43.87 | 32.4816043 |
| 7d7k | 1.9 | 0.54241448 | 9.96422679 | 47.83 | 35.9079114 |
| 7ofs | 1.9 | 0.3172899 | 16.5898207 | 51.79 | 33.0222647 |
| 7ybg | 1.9 | 0.88976412 | 6.29254421 | 34.69 | 37.1584617 |
| 7qck | 1.92 | 0.28873131 | 18.7618954 | 49.1 | 32.9386026 |
| 7jn2 | 1.93 | 0.30021555 | 15.2052852 | 54.38 | 31.5310069 |
| 7jit | 1.95 | 0.29041825 | 18.7411431 | 45.53 | 31.963886 |
| 7oft | 1.95 | 0.43989436 | 12.7550473 | 46.8 | 33.3421035 |
| 7sgw | 1.95 | 0.3973123 | 13.5003948 | 46.74 | 32.0707716 |
| 7d47 | 1.97 | 0.17318532 | 24.3997685 | 40.95 | 31.4917072 |
| 7kok | 2 | 0.17626932 | 20.6933588 | 61.07 | 31.4581265 |
| 7sgv | 2 | 0.2824366 | 14.8957274 | 58.97 | 31.5510135 |
| 7sqe | 2 | 0.39100083 | 11.5748566 | 54.07 | 32.7162717 |
| 8vec (M208W) | 2 | 1 | 0 | 32.92 | 32.92 |
| 7koj | 2.02 | 0.14335867 | 21.5478645 | 67.66 | 31.2475118 |
| 7m1y | 2.02 | 0.47744884 | 5.3308902 | 56.59 | 32.3497202 |
| 7rzc | 2.04 | 0.37184378 | 14.1975608 | 48.75 | 32.3249453 |
| 7jiv | 2.05 | 0.18424148 | 19.9304935 | 61.08 | 31.1839633 |
| 7jir | 2.09 | 0.17895473 | 20.1513364 | 62.68 | 31.3682191 |
| 7d7l | 2.11 | 0.37771096 | 15.5739496 | 52.43 | 35.3773352 |
| 8g62 | 2.17 | 0.33712231 | 14.1694767 | 55.62 | 32.9202195 |
| 7lbr | 2.2 | 0.46925852 | 11.5522862 | 43.63 | 32.0260353 |
| 7jiw | 2.3 | 0.13961797 | 19.2291798 | 82.63 | 30.765813 |
| 7llf | 2.3 | 0.43913362 | 8.25369361 | 52.87 | 31.4706882 |
| 7e35 | 2.4 | 0.06503379 | 28.1157246 | 72.89 | 32.8560372 |
| 6xg3 | 2.48 | 0.42301488 | 8.34495564 | 55.08 | 31.6446152 |
| 7jrn | 2.48 | 0.47797673 | 14.5753361 | 37.16 | 32.3369515 |
| 7cjd | 2.5 | 0.30854741 | 11.9386062 | 75.1 | 35.1105164 |
| 7kol | 2.58 | 0.19851585 | 12.402111 | 99.78 | 32.2100224 |
| 7cmd | 2.59 | 0.30114077 | 18.0708433 | 54.8 | 34.5733576 |
| 7tzj | 2.66 | 0.20445981 | 19.6145001 | 57.63 | 31.397519 |
| 7nt4 | 2.68 | 0.0552252 | 27.7951372 | 42.66 | 30.1510442 |
| 6w9c | 2.7 | 0.17810203 | 20.8143003 | 56.13 | 30.8111672 |
| 6xaa | 2.7 | 0.15537445 | 22.1539149 | 58.05 | 31.1734015 |
| 7krx | 2.72 | 0.10741612 | 20.0301615 | 103.1 | 31.1047631 |
| 7sdr | 2.72 | 0.30539089 | 13.4497592 | 59.66 | 31.6693799 |
| 6wuu | 2.79 | 0.24733127 | 12.3122497 | 80.91 | 32.323823 |
| 7lbs | 2.8 | 0.41229441 | 22.9090872 | 21.67 | 31.8435071 |
| 6xa9 | 2.9 | 0.16993909 | 17.8465475 | 102 | 35.1803343 |
| 7llz | 2.9 | 0.36917044 | 19.4338057 | 35.05 | 32.3732294 |
| 7los | 2.9 | 0.3580477 | 12.9197543 | 50.48 | 30.9940023 |
| 7rbs | 2.98 | 0.50526758 | -3.0652243 | 76.1 | 35.3856386 |
| 8eua | 3.1 | 0.21673041 | 12.7173902 | 90.65 | 32.3640022 |
| 6yva | 3.18 | 0.0517231 | 26.1909569 | 87.6 | 30.7219001 |
| 8cx9 | 3.5 | 0.26036956 | 17.8264929 | 63.1 | 34.2558121 |

## References

Adams, P. D., P. V. Afonine, G. Bunkóczi, V. B. Chen, N. Echols, J. J. Headd, L. W. Hung, S. Jain, G. J. Kapral, R. W. Grosse Kunstleve, A. J. McCoy, N. W. Moriarty, R. D. Oeffner, R. J. Read, D. C. Richardson, J. S. Richardson, T. C. Terwilliger, and P. H. Zwart. 2011. ‘The Phenix software for automated determination of macromolecular structures’, Methods, 55: 94–106.

Addis, Russell C., Fu-Chun Hsu, Rebecca L. Wright, Marc A. Dichter, Douglas A. Coulter, and John D. Gearhart. 2011. ‘Efficient Conversion of Astrocytes to Functional Midbrain Dopaminergic Neurons Using a Single Polycistronic Vector’, PLOS ONE, 6: e28719.

Aragão, D., J. Aishima, H. Cherukuvada, R. Clarken, M. Clift, N. P. Cowieson, D. J. Ericsson, C. L. Gee, S. Macedo, N. Mudie, S. Panjikar, J. R. Price, A. Riboldi-Tunnicliffe, R. Rostan, R. Williamson, and T. T. Caradoc-Davies. 2018. ‘MX2: a high-flux undulator microfocus beamline serving both the chemical and macromolecular crystallography communities at the Australian Synchrotron’, J Synchrotron Radiat, 25: 885–91.

Baez, Yahira. 2012. ’Insight into the Substrate Specificity and Inhibition of Human Coronavirus Papain-Like Proteases’, University of Illinois at Chicago.

Báez-Santos, Y. M., A. M. Mielech, X. Deng, S. Baker, and A. D. Mesecar. 2014. ‘Catalytic function and substrate specificity of the papain-like protease domain of nsp3 from the Middle East respiratory syndrome coronavirus’, J Virol, 88: 12511–27.

Báez-Santos, Y. M., S. E. St John, and A. D. Mesecar. 2015. ‘The SARS-coronavirus papain-like protease: structure, function and inhibition by designed antiviral compounds’, Antiviral Res, 115: 21–38.

Báez-Santos, Yahira M., Scott J. Barraza, Michael W. Wilson, Michael P. Agius, Anna M. Mielech, Nicole M. Davis, Susan C. Baker, Scott D. Larsen, and Andrew D. Mesecar. 2014. ‘X-ray Structural and Biological Evaluation of a Series of Potent and Highly Selective Inhibitors of Human Coronavirus Papain-like Proteases’, Journal of Medicinal Chemistry, 57: 2393–412.

Báez-Santos, Yahira M., Sarah E. St. John, and Andrew D. Mesecar. 2015. ‘The SARS-coronavirus papain-like protease: Structure, function and inhibition by designed antiviral compounds’, Antiviral Research, 115: 21–38.

Barretto, N., D. Jukneliene, K. Ratia, Z. Chen, A. D. Mesecar, and S. C. Baker. 2005. ‘The papain-like protease of severe acute respiratory syndrome coronavirus has deubiquitinating activity’, J Virol, 79: 15189–98.

Bloom, J. D. 2014. ‘An experimentally determined evolutionary model dramatically improves phylogenetic fit’, Mol Biol Evol, 31: 1956–78.

Brian Chia, C. S., and S. Pheng Lim. 2023. ‘A Patent Review on SARS Coronavirus Papain-Like Protease (PL(pro)) Inhibitors’, ChemMedChem, 18: e202300216.

Calleja, D. J., N. Kuchel, B. G. C. Lu, R. W. Birkinshaw, T. Klemm, M. Doerflinger, J. P. Cooney, L. Mackiewicz, A. E. Au, Y. Q. Yap, T. R. Blackmore, K. Katneni, E. Crighton, J. Newman, K. E. Jarman, M. J. Call, B. C. Lechtenberg, P. E. Czabotar, M. Pellegrini, S. A. Charman, K. N. Lowes, J. P. Mitchell, U. Nachbur, G. Lessene, and D. Komander. 2022. ‘Insights Into Drug Repurposing, as Well as Specificity and Compound Properties of Piperidine-Based SARS-CoV-2 PLpro Inhibitors’, Front Chem, 10: 861209.

Calleja, Dale J., Guillaume Lessene, and David Komander. 2022. ‘Inhibitors of SARS-CoV-2 PLpro’, Frontiers in Chemistry, 10.

Cassandri, Matteo, Artem Smirnov, Flavia Novelli, Consuelo Pitolli, Massimiliano Agostini, Michal Malewicz, Gerry Melino, and Giuseppe Raschellà. 2017. ‘Zinc-finger proteins in health and disease’, Cell Death Discovery, 3: 17071.

Chaudhuri, Rima, Sishi Tang, Guijun Zhao, Hui Lu, David A. Case, and Michael E. Johnson. 2011. ‘Comparison of SARS and NL63 Papain-Like Protease Binding Sites and Binding Site Dynamics: Inhibitor Design Implications’, Journal of Molecular Biology, 414: 272–88.

Choudhury, Debi, Sampa Biswas, Sumana Roy, and J.K. Dattagupta. 2010. ’Improving thermostability of papain through structure-based protein engineering’, *Protein Engineering*, Design and Selection, 23: 457–67.

Coghlan, Avril. 2011. ’A Little Book of R For Bioinformatics’, Cambridge, UK: *Welcome Trust Sanger Institute*.

Crooks, G. E., G. Hon, J. M. Chandonia, and S. E. Brenner. 2004. ‘WebLogo: a sequence logo generator’, Genome Res, 14: 1188–90.

Csibra, Eszter, and Guy-Bart Stan. 2022. ‘Absolute protein quantification using fluorescence measurements with FPCountR’, Nature Communications, 13: 6600.

Dickins, Ross A., Michael T. Hemann, Jack T. Zilfou, David R. Simpson, Ingrid Ibarra, Gregory J. Hannon, and Scott W. Lowe. 2005. ‘Probing tumor phenotypes using stable and regulated synthetic microRNA precursors’, Nature Genetics, 37: 1289–95.

Durfee, L. A., N. Lyon, K. Seo, and J. M. Huibregtse. 2010. ‘The ISG15 conjugation system broadly targets newly synthesized proteins: implications for the antiviral function of ISG15’, Mol Cell, 38: 722–32.

Emsley, P., B. Lohkamp, W. G. Scott, and K. Cowtan. 2010. ‘Features and development of Coot’, Acta Crystallogr D Biol Crystallogr, 66: 486–501.

Evans, P. R., and G. N. Murshudov. 2013. ‘How good are my data and what is the resolution?’, Acta Crystallogr D Biol Crystallogr, 69: 1204–14.

Faure, Andre J., Jörn M. Schmiedel, Pablo Baeza-Centurion, and Ben Lehner. 2020. ‘DiMSum: an error model and pipeline for analyzing deep mutational scanning data and diagnosing common experimental pathologies’, Genome Biology, 21: 207.

Fletcher, Roger. 2000. Practical methods of optimization (John Wiley & Sons).

Flynn, J. M., N. Samant, G. Schneider-Nachum, D. T. Barkan, N. K. Yilmaz, C. A. Schiffer, S. A. Moquin, D. Dovala, and D. N. A. Bolon. 2022. ‘Comprehensive fitness landscape of SARS-CoV-2 M(pro) reveals insights into viral resistance mechanisms’, Elife, 11.

Fowler, Douglas M., and Stanley Fields. 2014. ‘Deep mutational scanning: a new style of protein science’, Nature Methods, 11: 801–07.

French, S., and K. Wilson. 1978. ‘On the treatment of negative intensity observations’, Acta Crystallographica Section A, 34: 517–25.

Fu, Ziyang, Bin Huang, Jinle Tang, Shuyan Liu, Ming Liu, Yuxin Ye, Zhihong Liu, Yuxian Xiong, Wenning Zhu, Dan Cao, Jihui Li, Xiaogang Niu, Huan Zhou, Yong Juan Zhao, Guoliang Zhang, and Hao Huang. 2021. ‘The complex structure of GRL0617 and SARS-CoV-2 PLpro reveals a hot spot for antiviral drug discovery’, Nature Communications, 12: 488.

Gold, I. M., N. Reis, F. Glaser, and M. H. Glickman. 2022. ‘Coronaviral PLpro proteases and the immunomodulatory roles of conjugated versus free Interferon Stimulated Gene product-15 (ISG15)’, Semin Cell Dev Biol, 132: 16–26.

Gul, Sheraz, Syeed Hussain, Mark P. Thomas, Marina Resmini, Chandra S. Verma, Emrys W. Thomas, and Keith Brocklehurst. 2008. ‘Generation of Nucleophilic Character in the Cys25/His159 Ion Pair of Papain Involves Trp177 but Not Asp158’, Biochemistry, 47: 2025–35.

Harcourt, B. H., D. Jukneliene, A. Kanjanahaluethai, J. Bechill, K. M. Severson, C. M. Smith, P. A. Rota, and S. C. Baker. 2004. ‘Identification of severe acute respiratory syndrome coronavirus replicase products and characterization of papain-like protease activity’, J Virol, 78: 13600–12.

Hill, Andrew J., José L. McFaline-Figueroa, Lea M. Starita, Molly J. Gasperini, Kenneth A. Matreyek, Jonathan Packer, Dana Jackson, Jay Shendure, and Cole Trapnell. 2018. ‘On the design of CRISPR-based single-cell molecular screens’, Nature Methods, 15: 271–74.

Kabsch, W. 2010. ‘XDS’, Acta Crystallogr D Biol Crystallogr, 66: 125–32.

Klemm, Theresa, Gregor Ebert, Dale J Calleja, Cody C Allison, Lachlan W Richardson, Jonathan P Bernardini, Bernadine GC Lu, Nathan W Kuchel, Christoph Grohmann, Yuri Shibata, Zhong Yan Gan, James P Cooney, Marcel Doerflinger, Amanda E Au, Timothy R Blackmore, Gerbrand J van der Heden van Noort, Paul P Geurink, Huib Ovaa, Janet Newman, Alan Riboldi-Tunnicliffe, Peter E Czabotar, Jeffrey P Mitchell, Rebecca Feltham, Bernhard C Lechtenberg, Kym N Lowes, Grant Dewson, Marc Pellegrini, Guillaume Lessene, and David Komander. 2020. ‘Mechanism and inhibition of the papain-like protease, PLpro, of SARS-CoV-2’, The EMBO Journal, 39: e106275.

Lee, Hyun, Hao Lei, Bernard D. Santarsiero, Joseph L. Gatuz, Shuyi Cao, Amy J. Rice, Kavankumar Patel, Michael Z. Szypulinski, Isabel Ojeda, Arun K. Ghosh, and Michael E. Johnson. 2015. ‘Inhibitor Recognition Specificity of MERS-CoV Papain-like Protease May Differ from That of SARS-CoV’, ACS Chemical Biology, 10: 1456–65.

Lei, Jian, Jeroen R Mesters, Christian Drosten, Stefan Anemüller, Qingjun Ma, and Rolf Hilgenfeld. 2014. ‘Crystal structure of the papain-like protease of MERS coronavirus reveals unusual, potentially druggable active-site features’, Antiviral Research, 109: 72–82.

Lindner, H. A., N. Fotouhi-Ardakani, V. Lytvyn, P. Lachance, T. Sulea, and R. Ménard. 2005. ‘The papain-like protease from the severe acute respiratory syndrome coronavirus is a deubiquitinating enzyme’, J Virol, 79: 15199–208.

Lindner, H. A., V. Lytvyn, H. Qi, P. Lachance, E. Ziomek, and R. Ménard. 2007. ‘Selectivity in ISG15 and ubiquitin recognition by the SARS coronavirus papain-like protease’, Arch Biochem Biophys, 466: 8–14.

Lv, Z., K. E. Cano, L. Jia, M. Drag, T. T. Huang, and S. K. Olsen. 2021. ‘Targeting SARS-CoV-2 Proteases for COVID-19 Antiviral Development’, Front Chem, 9: 819165.

Ma, Chunlong, Michael Dominic Sacco, Zilei Xia, George Lambrinidis, Julia Alma Townsend, Yanmei Hu, Xiangzhi Meng, Tommy Szeto, Mandy Ba, Xiujun Zhang, Maura Gongora, Fushun Zhang, Michael Thomas Marty, Yan Xiang, Antonios Kolocouris, Yu Chen, and Jun Wang. 2021. ‘Discovery of SARS-CoV-2 Papain-like Protease Inhibitors through a Combination of High-Throughput Screening and a FlipGFP-Based Reporter Assay’, ACS Central Science, 7: 1245–60.

Martin, Marcel. 2011. ‘Cutadapt removes adapter sequences from high-throughput sequencing reads’, 2011, 17: 3.

Matreyek, K. A., L. M. Starita, J. J. Stephany, B. Martin, M. A. Chiasson, V. E. Gray, M. Kircher, A. Khechaduri, J. N. Dines, R. J. Hause, S. Bhatia, W. E. Evans, M. V. Relling, W. Yang, J. Shendure, and D. M. Fowler. 2018. ‘Multiplex assessment of protein variant abundance by massively parallel sequencing’, Nat Genet, 50: 874–82.

McCoy, A. J., R. W. Grosse-Kunstleve, P. D. Adams, M. D. Winn, L. C. Storoni, and R. J. Read. 2007. ‘Phaser crystallographic software’, J Appl Crystallogr, 40: 658–74.

Melnikov, A., P. Rogov, L. Wang, A. Gnirke, and T. S. Mikkelsen. 2014. ‘Comprehensive mutational scanning of a kinase in vivo reveals substrate-dependent fitness landscapes’, Nucleic Acids Res, 42: e112.

Ménard, R., J. Carrière, P. Laflamme, C. Plouffe, H. E. Khouri, T. Vernet, D. C. Tessier, D. Y. Thomas, and A. C. Storer. 1991. ‘Contribution of the glutamine 19 side chain to transition-state stabilization in the oxyanion hole of papain’, Biochemistry, 30: 8924–8.

Morgan, M., S. Anders, M. Lawrence, P. Aboyoun, H. Pagès, and R. Gentleman. 2009. ‘ShortRead: a bioconductor package for input, quality assessment and exploration of high-throughput sequence data’, Bioinformatics, 25: 2607–8.

Murphy, James M, Peter E Czabotar, Joanne M Hildebrand, Isabelle S Lucet, Jian-Guo Zhang, Silvia Alvarez-Diaz, Rowena Lewis, Najoua Lalaoui, Donald Metcalf, Andrew I Webb, Samuel N Young, Leila N Varghese, Gillian M Tannahill, Esme C Hatchell, Ian J Majewski, Toru Okamoto, Renwick C J. Dobson, Douglas J Hilton, Jeffrey J Babon, Nicos A Nicola, Andreas Strasser, John Silke, and Warren S Alexander. 2013. ‘The Pseudokinase MLKL Mediates Necroptosis via a Molecular Switch Mechanism’, Immunity, 39: 443–53.

Owen, Dafydd R., Charlotte M. N. Allerton, Annaliesa S. Anderson, Lisa Aschenbrenner, Melissa Avery, Simon Berritt, Britton Boras, Rhonda D. Cardin, Anthony Carlo, Karen J. Coffman, Alyssa Dantonio, Li Di, Heather Eng, RoseAnn Ferre, Ketan S. Gajiwala, Scott A. Gibson, Samantha E. Greasley, Brett L. Hurst, Eugene P. Kadar, Amit S. Kalgutkar, Jack C. Lee, Jisun Lee, Wei Liu, Stephen W. Mason, Stephen Noell, Jonathan J. Novak, R. Scott Obach, Kevin Ogilvie, Nandini C. Patel, Martin Pettersson, Devendra K. Rai, Matthew R. Reese, Matthew F. Sammons, Jean G. Sathish, Ravi Shankar P. Singh, Claire M. Steppan, Al E. Stewart, Jamison B. Tuttle, Lawrence Updyke, Patrick R. Verhoest, Liuqing Wei, Qingyi Yang, and Yuao Zhu. 2021. ‘An oral SARS-CoV-2 M^pro^ inhibitor clinical candidate for the treatment of COVID-19’, Science, 374: 1586–93.

Perlinska, Agata P., Adam Stasiulewicz, Mai Lan Nguyen, Karolina Swiderska, Mikolaj Zmudzinski, Alicja W. Maksymiuk, Marcin Drag, and Joanna I. Sulkowska. 2022. ‘Amino acid variants of SARS-CoV-2 papain-like protease have impact on drug binding’, PLOS Computational Biology, 18: e1010667.

Perng, Yi-Chieh, and Deborah J. Lenschow. 2018. ‘ISG15 in antiviral immunity and beyond’, Nature Reviews Microbiology, 16: 423–39.

Portelli, S., M. Olshansky, C. H. M. Rodrigues, E. N. D’Souza, Y. Myung, M. Silk, A. Alavi, D. E. V. Pires, and D. B. Ascher. 2020. ‘Exploring the structural distribution of genetic variation in SARS-CoV-2 with the COVID-3D online resource’, Nat Genet, 52: 999–1001.

Ratia, K., S. Pegan, J. Takayama, K. Sleeman, M. Coughlin, S. Baliji, R. Chaudhuri, W. Fu, B. S. Prabhakar, M. E. Johnson, S. C. Baker, A. K. Ghosh, and A. D. Mesecar. 2008a. ‘A noncovalent class of papain-like protease/deubiquitinase inhibitors blocks SARS virus replication’, Proc Natl Acad Sci U S A, 105: 16119–24.

Ratia, Kiira, Scott Pegan, Jun Takayama, Katrina Sleeman, Melissa Coughlin, Surendranath Baliji, Rima Chaudhuri, Wentao Fu, Bellur S. Prabhakar, Michael E. Johnson, Susan C. Baker, Arun K. Ghosh, and Andrew D. Mesecar. 2008b. ‘A noncovalent class of papain-like protease/deubiquitinase inhibitors blocks SARS virus replication’, Proceedings of the National Academy of Sciences, 105: 16119–24.

Ratia, Kiira, Kumar Singh Saikatendu, Bernard D. Santarsiero, Naina Barretto, Susan C. Baker, Raymond C. Stevens, and Andrew D. Mesecar. 2006. ‘Severe acute respiratory syndrome coronavirus papain-like protease: Structure of a viral deubiquitinating enzyme’, Proceedings of the National Academy of Sciences, 103: 5717–22.

Rut, Wioletta, Zongyang Lv, Mikolaj Zmudzinski, Stephanie Patchett, Digant Nayak, Scott J. Snipas, Farid El Oualid, Tony T. Huang, Miklos Bekes, Marcin Drag, and Shaun K. Olsen. 2020. ‘Activity profiling and crystal structures of inhibitor-bound SARS-CoV-2 papain-like protease: A framework for anti–COVID-19 drug design’, Science Advances, 6: eabd4596.

Shao, Qiang, Muya Xiong, Jiameng Li, Hangchen Hu, Haixia Su, and Yechun Xu. 2023. ‘Unraveling the catalytic mechanism of SARS-CoV-2 papain-like protease with allosteric modulation of C270 mutation using multiscale computational approaches’, Chemical Science, 14: 4681–96.

Shen, Z., K. Ratia, L. Cooper, D. Kong, H. Lee, Y. Kwon, Y. Li, S. Alqarni, F. Huang, O. Dubrovskyi, L. Rong, G. R. Thatcher, and R. Xiong. 2021. ’Potent, Novel SARS-CoV-2 PLpro Inhibitors Block Viral Replication in Monkey and Human Cell Cultures’, bioRxiv.

Shen, Zhengnan, Kiira Ratia, Laura Cooper, Deyu Kong, Hyun Lee, Youngjin Kwon, Yangfeng Li, Saad Alqarni, Fei Huang, Oleksii Dubrovskyi, Lijun Rong, Gregory R. J. Thatcher, and Rui Xiong. 2022. ‘Design of SARS-CoV-2 PLpro Inhibitors for COVID-19 Antiviral Therapy Leveraging Binding Cooperativity’, Journal of Medicinal Chemistry, 65: 2940–55.

Shenoy, A. R., and S. S. Visweswariah. 2003. ‘Site-directed mutagenesis using a single mutagenic oligonucleotide and DpnI digestion of template DNA’, Anal Biochem, 319: 335–6.

Shin, D., R. Mukherjee, D. Grewe, D. Bojkova, K. Baek, A. Bhattacharya, L. Schulz, M. Widera, A. R. Mehdipour, G. Tascher, P. P. Geurink, A. Wilhelm, G. J. van der Heden van Noort, H. Ovaa, S. Müller, K. P. Knobeloch, K. Rajalingam, B. A. Schulman, J. Cinatl, G. Hummer, S. Ciesek, and I. Dikic. 2020. ‘Papain-like protease regulates SARS-CoV-2 viral spread and innate immunity’, Nature, 587: 657–62.

Shiraishi, Yutaro, and Ichio Shimada. 2023. ‘NMR Characterization of the Papain-like Protease from SARS-CoV-2 Identifies the Conformational Heterogeneity in Its Inhibitor-Binding Site’, Journal of the American Chemical Society, 145: 16669–77.

Skaug, Brian, and Zhijian J. Chen. 2010. ‘Emerging Role of ISG15 in Antiviral Immunity’, Cell, 143: 187–90.

Smith, T., A. Heger, and I. Sudbery. 2017. ‘UMI-tools: modeling sequencing errors in Unique Molecular Identifiers to improve quantification accuracy’, Genome Res, 27: 491–99.

Starr, T. N., A. J. Greaney, S. K. Hilton, D. Ellis, K. H. D. Crawford, A. S. Dingens, M. J. Navarro, J. E. Bowen, M. A. Tortorici, A. C. Walls, N. P. King, D. Veesler, and J. D. Bloom. 2020. ‘Deep Mutational Scanning of SARS-CoV-2 Receptor Binding Domain Reveals Constraints on Folding and ACE2 Binding’, Cell, 182: 1295–310.e20.

Steiner, P. J., Z. T. Baumer, and T. A. Whitehead. 2020. ‘A Method for User-defined Mutagenesis by Integrating Oligo Pool Synthesis Technology with Nicking Mutagenesis’, Bio Protoc, 10: e3697.

Thorn, K. 2017. ‘Genetically encoded fluorescent tags’, Mol Biol Cell, 28: 848–57.

Ton, A. T., M. Pandey, J. R. Smith, F. Ban, M. Fernandez, and A. Cherkasov. 2022. ‘Targeting SARS-CoV-2 papain-like protease in the postvaccine era’, Trends Pharmacol Sci, 43: 906–19.

Trenker, R., X. Wu, J. V. Nguyen, S. Wilcox, A. F. Rubin, M. E. Call, and M. J. Call. 2021. ‘Human and viral membrane-associated E3 ubiquitin ligases MARCH1 and MIR2 recognize different features of CD86 to downregulate surface expression’, J Biol Chem, 297: 100900.

Vernet, Thierry, Daniel C. Tessier, Jean Chatellier, Céline Plouffe, Tak Sing Lee, David Y. Thomas, Andrew C. Storer, and Robert Ménard. 1995. ‘Structural and Functional Roles of Asparagine 175 in the Cysteine Protease Papain (&#x2217;)’, Journal of Biological Chemistry, 270: 16645–52.

Wrenbeck, E. E., J. R. Klesmith, J. A. Stapleton, A. Adeniran, K. E. Tyo, and T. A. Whitehead. 2016. ‘Plasmid-based one-pot saturation mutagenesis’, Nat Methods, 13: 928–30.

Wydorski, Pawel M., Jerzy Osipiuk, Benjamin T. Lanham, Christine Tesar, Michael Endres, Elizabeth Engle, Robert Jedrzejczak, Vishruth Mullapudi, Karolina Michalska, Krzysztof Fidelis, David Fushman, Andrzej Joachimiak, and Lukasz A. Joachimiak. 2023. ‘Dual domain recognition determines SARS-CoV-2 PLpro selectivity for human ISG15 and K48-linked di-ubiquitin’, Nature Communications, 14: 2366.

Xia, Z., M. D. Sacco, C. Ma, J. A. Townsend, N. Kitamura, Y. Hu, M. Ba, T. Szeto, X. Zhang, X. Meng, F. Zhang, Y. Xiang, M. T. Marty, Y. Chen, and J. Wang. 2021. ‘Discovery of SARS-CoV-2 papain-like protease inhibitors through a combination of high-throughput screening and FlipGFP-based reporter assay’, bioRxiv.

Zhang, Mengdi, Jingxian Li, Haiyan Yan, Jun Huang, Fangwei Wang, Ting Liu, Linghui Zeng, and Fangfang Zhou. 2021. ‘ISGylation in Innate Antiviral Immunity and Pathogen Defense Responses: A Review’, Frontiers in Cell and Developmental Biology, 9.

Zhang, Yuan, Xianliang Ke, Caishang Zheng, Yan Liu, Li Xie, Zhenhua Zheng, and Hanzhong Wang. 2017. ‘Development of a luciferase-based biosensor to assess enterovirus 71 3C protease activity in living cells’, Scientific Reports, 7: 10385.

Zhao, Yao, Xiaoyu Du, Yinkai Duan, Xiaoyan Pan, Yifang Sun, Tian You, Lin Han, Zhenming Jin, Weijuan Shang, Jing Yu, Hangtian Guo, Qianying Liu, Yan Wu, Chao Peng, Jun Wang, Chenghao Zhu, Xiuna Yang, Kailin Yang, Ying Lei, Luke W. Guddat, Wenqing Xu, Gengfu Xiao, Lei Sun, Leike Zhang, Zihe Rao, and Haitao Yang. 2021. ‘High-throughput screening identifies established drugs as SARS-CoV-2 PLpro inhibitors’, Protein & Cell, 12: 877–88.

## References

Armstrong, L. A., S. M. Lange, V. Dee Cesare, S. P. Matthews, R. S. Nirujogi, I. Cole, A. Hope, F. Cunningham, R. Toth, R. Mukherjee, D. Bojkova, F. Gruber, D. Gray, P. G. Wyatt, J. Cinatl, I. Dikic, P. Davies, and Y. Kulathu. 2021. ‘Biochemical characterization of protease activity of Nsp3 from SARS-CoV-2 and its inhibition by nanobodies’, PLOS ONE, 16: e0253364.

Kamphuis, I. G., K. H. Kalk, M. B. A. Swarte, and J. Drenth. 1984. ‘Structure of papain refined at 1.65 Å resolution’, Journal of Molecular Biology, 179: 233–56.

