## Supplementary material for "Mutational Profiling of SARS-CoV-2 PLpro in human cells reveals requirements for function, structure, and drug escape": R script for B factor analysis

library('tidyverse')
library('gridExtra')
#the files are downloaded from pdb 
#the filter is made with PLpro sequence
#Only SARS-CoV-2 PLpro are chosen
#loop everything and output the final results
pdb_b_factor <- lapply(list.files('~/batch-download-structures-1697694010604/'), function(pdb_name) {
  #build up the connection
  one_file <- gzcon(file(paste0("~/batch-download-structures-1697694010604/", pdb_name), "rb")) #start by connecting with the file
  #read the file by lines output it if it reports the b-factor
  temp_df <- list() #empty list to hold the file 
  i <- 1 #indexing to input the results
  while(length(line <- readLines(one_file, n = 1)) > 0) {
    if (substr(line, 1, 4) == "ATOM") { #it it starts with ATOM (reports b-factor)
      temp_string <- unlist(strsplit(line, "\\s+")) %>% as.data.frame() #make a temp_strings 
      if (nrow(temp_string) == 12) {temp_df[[i]] <- temp_string} #if the output is standarised, report it
      i <- i + 1 #increase indexing to the end
    } else {i <- i + 1}
  }
  # Close the connection
  close(one_file)
  #remove NA if there is no data
  temp_df <- temp_df[!sapply(temp_df, is.null)]
  b_factor_df <- do.call(cbind, temp_df) %>% t(.) #merge data
  rownames(b_factor_df) <- seq(1, nrow(b_factor_df), 1) #make rowname numeric
  b_factor_df <- as.data.frame(b_factor_df) #make it a dataframe for tidyverse
  #make position and b-factor numberic
  b_factor_df <- b_factor_df %>% mutate(V6 = as.numeric(V6), V11 = as.numeric(V11))
  #make sure only chain A is kept
  b_factor_df <- b_factor_df %>% filter(V5 == "A")
  #make sure the numbering method is the same
  #D286 as active triads should be the same, 5 residues ahead/after are not D so can use it as probe
  true_pos <- 0
  for (i in seq(281, 291, 1)) {
    #Amino acid
    aa_at_pos <- b_factor_df %>% filter(V6 == i) %>% .[1, 'V4']
    if (is.na(aa_at_pos)) {true_pos_error <- "Error"; break}
    if (aa_at_pos == 'ASP') {true_pos <- i} else {true_pos_error <- "Error"}
  } #find the true D286 position, if not found, report error
  if (true_pos_error == "Error" & true_pos == 0) {return(NULL)}
  #otherwise, define move fatcor
  b_factor_df <- b_factor_df %>% mutate(V6 = V6 + 286 - true_pos)
  #Now, summarise the average b-factor and report it as final dataframe
  #final_df <- b_factor_df[, c('V4','V6','V11')] %>% 
  #  dplyr::group_by(V6) %>% dplyr::summarise(average_b_factor = mean(V11))
  #instead of calculating average b factor for all, just do backbone and Cb
  final_df <- b_factor_df[, c('V3', 'V4','V6','V11')] %>% 
    filter(V3 %in% c('N','CA','C','O','CB')) %>% dplyr::group_by(V6) %>% dplyr::summarise(average_b_factor = mean(V11))
  #return data
  return(final_df)
})
#rename it
names(pdb_b_factor) <- list.files('~/batch-download-structures-1697694010604/')
#remove null
pdb_b_factor <- pdb_b_factor[!sapply(pdb_b_factor, is.null)]

##add a new column
pdb_b_factor_name <- lapply(names(pdb_b_factor), function(list_name){
  #add new columns
  temp_df <- pdb_b_factor[[list_name]] %>% 
    mutate(PDB = str_split(list_name, '.pdb', simplify = T)[1])
})
names(pdb_b_factor_name) <- names(pdb_b_factor)
pdb_b_factor_merged <- do.call(rbind, pdb_b_factor_name)


#define weight based on the average b-factor for all PDB structures
weight_df <- pdb_b_factor_merged %>% 
  dplyr::group_by(V6) %>% 
  dplyr::summarise(average_b_factor = mean(average_b_factor)) %>% 
  as.data.frame()

#merge M208W and weighting df for quick calculation
function_df <- merge(weight_df, pdb_b_factor_merged %>% filter(PDB == 'M208W'),
                     by = c('V6'), suffixes = c('_weight','_M208W')) %>% dplyr::select(-c(PDB))


#define penalty function
norm_function <- function(params, target_Y, standard_df) {
  #assume params[1] is scaleing and params[2] is translating factor
  #transform the data
  target_Y$average_b_factor <- params[1] * target_Y$average_b_factor + params[2]
  #merge data
  norm_df <- merge(target_Y, standard_df, by = c('V6'),
                   suffixes = c('_sample', 'function_df'))
  #calculate distance in portion to peak height
  output_sum <- sum(((norm_df$average_b_factor - norm_df$average_b_factor_M208W) / norm_df$average_b_factor_weight)^2)
  return(output_sum)
}

#check every PDB files and perform optimum functions
scale_PDB <- lapply(unique(pdb_b_factor_merged$PDB)[-55], function(PDB_name){
  #find the dataframe
  temp_df <- pdb_b_factor_merged %>% filter(PDB == PDB_name)
  #propose initial values
  initial_scale_guess <- mean(function_df$average_b_factor_M208W) / mean(temp_df$average_b_factor)
  initial_translate_guess <- 0 #this is the initial guess after applying the initial scale guess and calculated with: mean(function_df$average_b_factor_M208W) - mean(temp_df$average_b_factor * initial_scale_guess)
  params <- c(initial_scale_guess, initial_translate_guess) #this is the initial value
  #remove Cysteine region for better performance
  temp_df <- temp_df %>% filter(V6 <= 185 | (V6 >= 198 & V6 <= 218) |
                                  V6 >= 233)
  #perform optimisation
  result <- optim(
    par = params,
    fn = norm_function,
    target_Y = temp_df,
    standard_df = function_df,
    method = "BFGS"
  )
  #return the extract parameters and rescale
  optimized_scale <- result$par[1]
  optimized_translation <- result$par[2]
  #rescale data
  temp_df$average_b_factor <- optimized_scale * temp_df$average_b_factor + optimized_translation
  return(list(temp_df, optimized_scale, optimized_translation))
})


#####from this part, all factors have been output#####
#extract the scaling factor matrix
scaling_factors <- lapply(scale_PDB, `[[`, 2)
translating_factors <- lapply(scale_PDB, `[[`, 3)

factors_dataframe <- data.frame(
  PDB = unique(pdb_b_factor_merged$PDB)[-55],
  scaling = unlist(scaling_factors),
  translating = unlist(translating_factors)
) #create this dataframe for final report

#sort by resolution and wilson's b factor
#loop everything and output the final results
#only reports the data that have been used in previous studies
used_files_index <- list.files('~/batch-download-structures-1697694010604/') %>% 
  str_split(., '\\.', simplify = T) %>% .[-58, 1] %>% as.vector() %>% 
  {which(. %in% unique(pdb_b_factor_merged$PDB)[-55])}

used_files <- list.files('~/batch-download-structures-1697694010604/')[used_files_index]

resolution_pdb <- lapply(used_files, function(pdb_name) {
  #build up the connection
  one_file <- gzcon(file(paste0("~/batch-download-structures-1697694010604/", pdb_name), "rb")) #start by connecting with the file
  #read the file by lines output it if it reports resolution
  i <- 1 #indexing to input the results
  while(length(line <- readLines(one_file, n = 1)) > 0) {
    if (grepl('RESOLUTION', line)) { #find the line with resolution
      temp_df <- line
      break
      i <- i + 1 #increase indexing to the end
    } else {i <- i + 1}
  }
  # Close the connection
  close(one_file)
  return(temp_df)
})
#loop everything and output the final results
wilson_b <- lapply(used_files, function(pdb_name) {
  #build up the connection
  one_file <- gzcon(file(paste0("~/PDB_PLpro/batch-download-structures-1697694010604/", pdb_name), "rb")) #start by connecting with the file
  wilson <- grep("MEAN B VALUE", readLines(one_file), value = TRUE)
  # Close the connection
  close(one_file)
  return(wilson)
})

#rename it
names(resolution_pdb) <- used_files
names(wilson_b) <- used_files

#remove null
resolution_pdb <- resolution_pdb[!sapply(resolution_pdb, is.null)]
#make it a dataframe
resolution_df_list <- lapply(names(resolution_pdb), function(resolution_str) {
  temp_str <- str_split(resolution_pdb[[resolution_str]], 'RESOLUTION.', simplify = T)[2] %>% 
    str_split(., 'ANGSTROMS', simplify = T) %>% as.vector() %>% .[1] %>% as.numeric(gsub('[^[:digit:]]','',.))
  temp_df <- data.frame(PDB = str_split(resolution_str,"\\.", simplify = T)[1],
                        Resolution = temp_str)
  return(temp_df)
})
resolution_df <- do.call(rbind, resolution_df_list) %>% 
  rbind(data.frame(PDB = 'M208W', Resolution = 2.00)) %>% 
  arrange(Resolution)

#merge wilson's b
#remove null
wilson_b <- wilson_b[!sapply(wilson_b, is.null)]
#make it a dataframe
wilson_df_list <- lapply(names(wilson_b), function(resolution_str) {
  temp_str <- as.numeric(sub(".*: ([0-9.]+).*", "\\1", wilson_b[[resolution_str]]))
  temp_df <- data.frame(PDB = str_split(resolution_str,"\\.", simplify = T)[1],
                        wilson = temp_str)
  return(temp_df)
})
wilson_df <- do.call(rbind, wilson_df_list) %>% 
  rbind(data.frame(PDB = 'M208W', wilson = 32.92)) %>% 
  arrange(wilson)

###merge all dfs
final_df <- merge(
  wilson_df, resolution_df, by = c('PDB'), all = T
) %>% merge(., factors_dataframe, by = c('PDB'), all = T)
#introduce M208W data
final_df[55, c(4,5)] <- c(1, 0)

#calculate moved wilson factor
final_df <- final_df %>% mutate(moved_wilson = scaling * wilson + translating)
write_tsv(final_df, 
          '~/parameters_for_alignment_with_M208W.txt'
)


#####this part is to apply the same algorithm to analysis all other PLpro structures#####
#####this is done by choosing any PLpro as standard and to align the b factors on it#####

weight_df <- pdb_b_factor_merged %>% 
  dplyr::group_by(V6) %>% dplyr::summarise(average_b_factor = mean(average_b_factor)) %>% 
  as.data.frame()

#penalty function
norm_function <- function(params, target_Y, standard_df) {
  #assume params[1] is scaleing and params[2] is translating factor
  #transform the data
  target_Y$average_b_factor <- params[1] * target_Y$average_b_factor + params[2]
  #merge data
  norm_df <- merge(target_Y, standard_df, by = c('V6'),
                   suffixes = c('_sample', 'function_df'))
  #calculate distance in portion to peak height
  output_sum <- sum(((norm_df$average_b_factor - norm_df$average_b_factor_sample) / norm_df$average_b_factor_weight)^2)
  return(output_sum)
}

#search every PDB
check_all_PDB <- lapply(unique(pdb_b_factor_merged$PDB), function(PDB_main_name) {
  #make the standard df
  function_df <- merge(weight_df, pdb_b_factor_merged %>% filter(PDB == PDB_main_name),
                       by = c('V6'), suffixes = c('_weight', '_sample')) %>% dplyr::select(-PDB)
  #optimise all other PDBs
  #check every PDB files and perform optimum functions
  scale_PDB <- lapply(unique(pdb_b_factor_merged$PDB)[-which(unique(pdb_b_factor_merged$PDB) == PDB_main_name)], function(PDB_name){
    #find the dataframe
    temp_df <- pdb_b_factor_merged %>% filter(PDB == PDB_name)
    #propose initial values
    initial_scale_guess <- mean(function_df$average_b_factor_sample) / mean(temp_df$average_b_factor)
    initial_translate_guess <- 0
    params <- c(initial_scale_guess, initial_translate_guess) #this is the initial value
    #remove Cysteine region for better performance
    temp_df <- temp_df %>% filter(V6 <= 185 | (V6 >= 198 & V6 <= 218) |
                                    V6 >= 233)
    #perform optimisation
    result <- optim(
      par = params,
      fn = norm_function,
      target_Y = temp_df,
      standard_df = function_df,
      method = "BFGS"
    )
    #return the extract parameters and rescale
    optimized_scale <- result$par[1]
    optimized_translation <- result$par[2]
    #rescale data
    temp_df$average_b_factor <- optimized_scale * temp_df$average_b_factor + optimized_translation
    return(temp_df)
  })
  #merge dataframe and plot
  final_df <- do.call(rbind, scale_PDB) %>% rbind(., pdb_b_factor_merged %>% filter(PDB == PDB_main_name))
  #plot
  p <- ggplot(final_df) +
    geom_line(aes(x = V6, y = average_b_factor, color = PDB), alpha = 0.5) +
    geom_line(data = final_df %>% filter(PDB == PDB_main_name),
              aes(x = V6, y = average_b_factor), color = 'Darkred', size = 2, alpha = 1) +
    theme(strip.text = element_blank()) + #geom_vline(xintercept = Pos_num, linetype = 'dashed') +
    geom_vline(xintercept = 208) + geom_vline(xintercept = 166) +
    scale_y_continuous(limit = c(0,60)) +
    ylab('Average B factor') + xlab('Pos') + labs(title = paste0('normalised with ', PDB_main_name)) +
    xlim(-5,320) + coord_cartesian(expand = F)
  
  #Find the outliers utilising the means
  base_line <- final_df %>% filter(PDB != PDB_main_name) %>% 
    dplyr::group_by(V6) %>% 
    dplyr::summarise(Avg_b = mean(average_b_factor),
                     SD = sd(average_b_factor)) %>% as.data.frame()
  merged_base <- merge(base_line, final_df %>% filter(PDB == PDB_main_name),
                       by = c('V6'))
  merged_base <- na.omit(merged_base)
  merged_base <- merged_base %>% 
    mutate(distance = (average_b_factor - Avg_b) / SD)
  merged_base <- merged_base %>% 
    mutate(classification = case_when(
      distance >= 3 ~ 'Destabilising',
      distance >= 2 ~ 'Weak destabilising',
      distance <= -3 ~ 'Stabilising',
      distance <= -2 ~ 'Weak stabilising',
      TRUE ~ 'Within range'
    ))
  #stablising
  stablising <- merged_base %>% filter(classification == 'Stabilising') %>% .$V6 %>% .[order(.)]
  #destabilising
  destabilising <- merged_base %>% filter(classification == 'Destabilising') %>% .$V6 %>% .[order(.)]
  #Weak destabilising
  Weak_destabilising <- merged_base %>% filter(classification == 'Weak destabilising') %>% .$V6 %>% .[order(.)]
  #Weak stabilising
  Weak_stabilising <- merged_base %>% filter(classification == 'Weak stabilising') %>% .$V6 %>% .[order(.)]
  #make a list
  report_resi <- list(stablising, destabilising, Weak_destabilising, Weak_stabilising)
  #return result
  return(list(report_resi, p))
})

#rename PDB
names(check_all_PDB) <- unique(pdb_b_factor_merged$PDB)

#extract classes
extract_class <- lapply(unique(pdb_b_factor_merged$PDB), function(sublist_name) {
  #report the length as dataframe elements
  temp_df <- data.frame(
    stablised = length(check_all_PDB[[sublist_name]][[1]]),
    destablised = length(check_all_PDB[[sublist_name]][[2]]),
    weak_destablised = length(check_all_PDB[[sublist_name]][[3]]),
    weak_stablised = length(check_all_PDB[[sublist_name]][[4]]),
    PDB_name = sublist_name
  )
})
extract_class_df <- do.call(rbind, extract_class)
